# A global map of the impact of deletion of Post-Translational Modification sites in genetic diseases

**DOI:** 10.1101/2020.12.20.423666

**Authors:** Perceval Vellosillo, Pablo Minguez

## Abstract

**Background:** There are >200 protein post-translational modification (PTMs) types described in eukaryotes, having diverse species conservation levels, proteome coverage, number of high-throughput experiments and functional roles. From a clinical perspective, a number of diseases have been associated to deregulated PTM sites and missense rare variants are globally enriched in PTMs. We hypothesize that some genetic diseases may be caused by the deregulation of particular functions produced by the removal of a specific PTM type by genomic variants.

**Results:** We collected >320,000 human PTMs of 59 types and cross them with >4M missense DNA variants annotated with pathogenic predictions and disease associations. We report >1.74M PTM-variant concurrences in >16,500 proteins that an enrichment analysis distributed in 217 pairwise significant associations between 18 PTM types and 150 genetic diseases. Around 23% of these associations are already described in the literature, 34% have partial evidences based on single variants, related diseases or regulatory evidences, and 43% are novel. Removal of acetylation presents the highest effect, still low studied PTM types like S-glutathionylation or S-nitrosylation show relevance. A network of PTM types and phenotypes associations is also discussed. Using pathogenicity predictions we identified potential PTM sites to produce particular diseases if genomic variants remove them.

**Conclusions:** Our results show an important impact of PTM removal producing genetic diseases and phenotypes that is PTM type specific. We describe for the first time a general scenario of PTM types and genetic diseases direct associations, many of them novel, that provides new capacities to understand and diagnose these disorders.

## Background

Post-translational modifications (PTMs) are an essential source of protein regulation. They are mostly reversible additions of small moieties or proteins, able to reprogram rapidly protein function. PTMs are ubiquitous in the eukaryotic proteome and have a large global stoichiometry anytime. There are several hundreds of PTM types, each defined by their own features, both chemical and functional [1,2], are conserved at different degrees [3], have been studied unequally by large scale experiments [4] and their occupancy and combination are responsible for protein activity and localization [3,5].

Apart from their specific regulation or enzymatic failures, together with the differential isoform transcription [6], genomic variation is the main source of PTM lack of occupancy. In particular, non synonymous single nucleotide variations (nsSNVs) in codons coding for a modifiable residue or within the enzyme recognized motif, impede permanently the modification. This effect has been largely studied in the context of somatic genomic variation in cancers, revealing a significant impact of at least the most abundant types (phosphorylation, acetylation or ubiquitination) in tumors development [7–9]. Regarding germline DNA variation, a large scale survey have described an enrichment of rare and disease annotated variants in PTM regions [10] suggesting that functions provided by PTMs are linked to disease phenotypes. Although not many other works are available for a global view, PTMs are gaining much attention in this context, as shown by the number of public resources providing mappings [4,11–14], tools for their usage for genetic diagnosis [15,16], or the application of genetic engineering to introduce PTMs in recombinant biopharmaceuticals favoring their functionality [17]. Not surprisingly, the main features of PTM sites, functionality, exposure or conservation, are also the main input for most variant pathogenicity/tolerance predictors [18,19].

The functional landscape of every type of PTM is still being decoded [3,20]. Meanwhile, their dysfunction have been associated to hundreds of diseases already. Major attention has caught phosphorylation, that besides of cancers [7], has been described to be associated to other disorders like Alzheimer [21], or cardiovascular diseases [22]. Other types of PTMs have also been linked to cancers: methylation and acetylation by their regulation of histone tails [23,24], ubiquitination through the alteration of protein degradation [25] or S-nitrosylation by its regulation of tumor proliferation [26]. Target residues of acetylation, methylation, glycosylation or S-nitrosylation are reported to be dysfunctional in cardiovascular and neurological disorders [27–31]. Glycation was found to be deregulated in diabetes mellitus [32] and N and O-Glycosylation in distinct genetic congenital disorders [33]. The ubiquitination writer system is found to be key in many rare congenital disorders like the Angelman or von Hippel-Lindau syndromes among many others [25].

In this work we hypothesize that the loss of specific protein functions defined by particular types of PTMs may converge in similar disease phenotypes. Thus, we performed a crossmatching between human PTMs of different types and nsSNVs with disease annotation and deleterous predictions, and found a large number of associations between disorders and PTM types, some of them with very low proteome coverage. We report herein the first systematic and global survey of associations between genetic diseases and the lack of specific PTM types. These results will encourage to gain attention on particular PTM types in the study of genetic disorders as well as to provide a massive volume of new hypothesis in the research of rare diseases.

## Results

### Crossmatching of PTMs and genomic variants

We collected 321,643 human PTMs of 59 types [4,34–42] (Figure 1A) and 4,464,412 nsSNVs [39,43–46], 392,185 (8.8%) of them classified as pathogenic using predictors [18,47–50] and 44,969 (1%) annotated with their disease-associations taken from public repositories [39,45,46,51,52] (Figure 1B, Tables S1-3). All PTMs and nsSNVs variants were crossmatched to identify residues with reported modifications that are prevented by certain nsSNVs. We name here an affected PTM, also a PTM-nsSNV co-occurrence, when a missense variant mutates: 1) the codon coding the modifiable amino acid or, 2) the codons coding a window of ±5 amino acids from the modifiable residue, for PTM types with motifs reported (see Materials and Methods, Figure S1). We identified a total of 1,743,207 PTM-nsSNVs co-occurrences (Table S4, Supplementary Data 1), 15,607 with nsSNVs involved in diseases (Table S5) and 133,292 with nsSNVs predicted to be pathogenic (Table S6). To highlight PTM types affected by DNA variation above the general trend we calculated the average proportion of a PTM of any type to be impacted by: i) any, ii) pathogenic, and iii) disease-associated variants, being 74.1%, 20.3% and 2.4% respectively (Figure 1C). Thus, up to 11 (18.6%) PTM types are above the baseline probability of being impacted by any type of nsSNV, including O-GlcNAc glycosylation, malonylation and neddylation and eight PTM types from the top 10 most abundant classes, only phosphorylation and ubiquitination left out. Interestingly, neddylation, with only 40 instances collected, shows the highest proportion, not only of variants matching modified sites (100%), but also those predicted as pathogenic (60%) and annotated as disease-associated (17.5%). Besides neddylation, other 11 types are above the threshold for the proportion of PTMs potentially removed by pathogenic variants, 3 types of glycosylations (NG, OG and OGl), malonylation, S-glutathionylation, S-nitrosylation, and the 5 most studied and abundant classes: phosphorylation, ubiquitination, acetylation, SUMOylation and methylation. In terms of variants associated to diseases, 13 types are above the baseline probability, with carboxylation as the second most impacted type.

**Figure 1.**
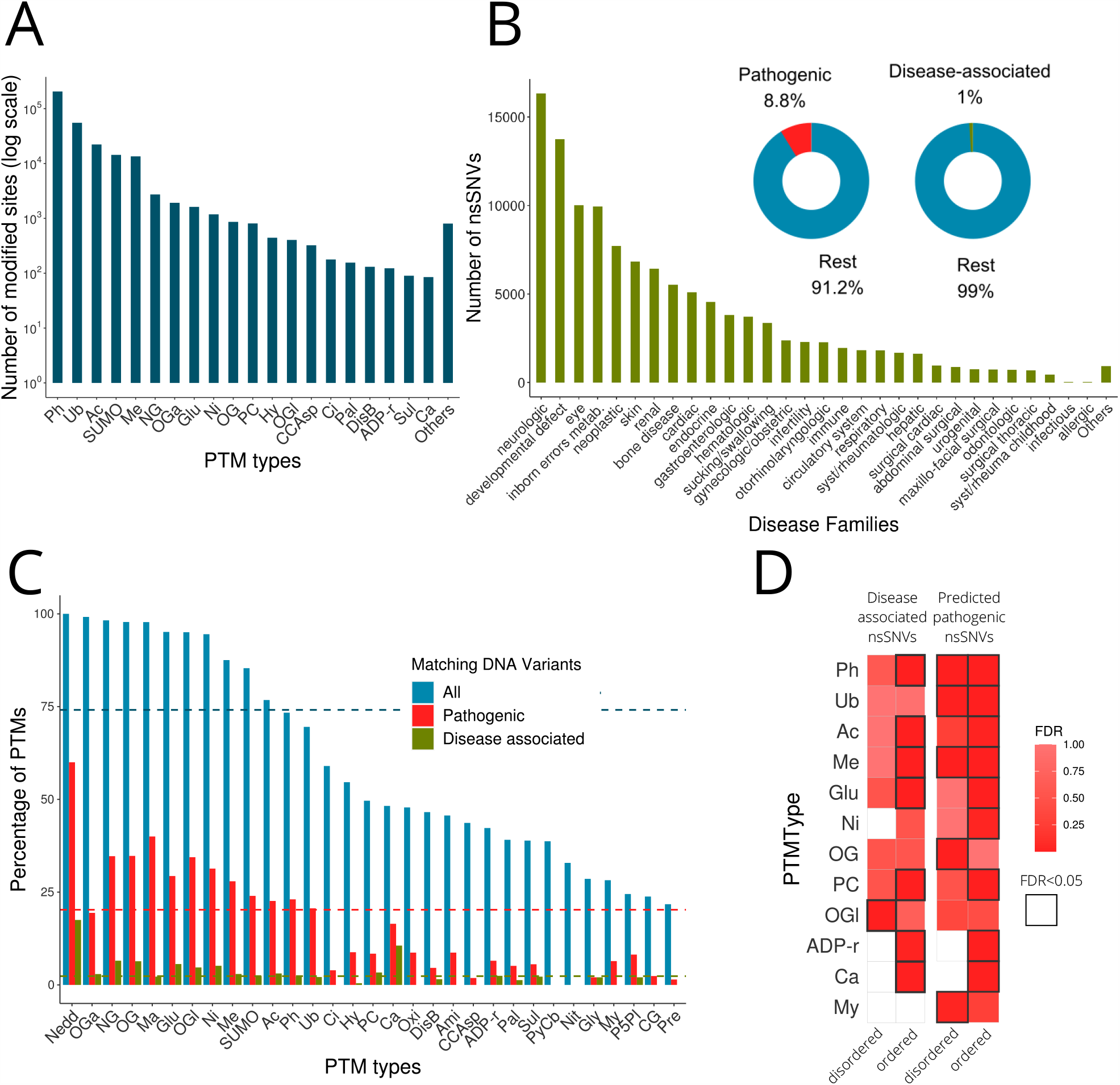
Human PTMs and nsSNVs compiled and details of their crossmatching and PTM types enriched in disease associated and predicted pathogenic nsSNVs. A. Number of PTMs classified by type. B. Number of nsSNVs classified in Orphanet families. Donut charts represent the percentage of variants predicted pathogenic and involved in diseases. C. Crossmatching between PTMs and nsSNVs per PTM type. Horizontal colored lines represent the average percentage of all PTM types to match all, pathogenic and disease-associated nsSNVs. D. Enrichment of pathogenic and disease causing variants affecting PTM types.

In order to assess whether there are PTM types whose removal is more deleterious than others, we tested if pathogenic and disease-associated nsSNVs impact recurrently not just on PTMs but on certain PTM types (Figure 1D, Tables S7,S8). As PTM functionality might depend on the protein region where the PTM is located, we tested independently ordered and disordered regions. First observation is that PTM types are, in general, more enriched in predicted pathogenic nsSNVs than in disease-associated. The removal of PTMs at ordered regions seems, also globally, more deleterious than those in disordered regions. Only phosphorylation, ubiquitination and methylation show a functionality that seems equally impacted by nsSNVs in both ordered and disordered regions. Other 12 classes show enrichments only in one protein region type, most times in ordered. ADP-ribosylation, carboxylation and myristoylation are the PTM types with lower number of instances (123, 85 and 78 respectively) that are hit significantly by disease-related or pathogenic DNA variants.

### A global network of PTM types and inherited diseases associations

To extract significant pairwise associations between specific PTM types and particular genetic diseases we performed Fisher’s exact test to check if the proportion of nsSNVs causing a genetic disease and hitting in a PTM type in respect to other sites, is greater than the ratio of nsSNVs causing other genetic diseases affecting the same PTM type, also compared to other sites. As result we found 217 significant associations (FDR<0.05) between 18 PTM types and 150 inheritance diseases (Table S9) that we annotated performing a literature review as: i) known, ii) having partial evidences, or iii) novel. Within partial evidences we comprised several heterogeneous documented examples that although do not report an association between the removal of the PTM type and the disorder, may suggest a link indirectly. They include: i) reports of one variant case; ii) associations of the PTM type with a related or similar disease; and iii) regulatory evidences when the PTM type is involved the deregulation causing the disease. Summing up, our strategy was able to catch 50 known associations, generalize 74 and provide for the first time 93 novel links (Figure 2A) which are unequally distributed over PTM types (Figure 2B). Figure 2C displays all predicted PTM types and disease associations in a network format. Thus, acetylation stands up as the PTM with more associations, 61 in total, 36 of them not previously reported, and only 6 known. Next, ubiquitination and N-linked glycosylation are associated with 29 and 22 pathologies respectively, and other 6 PTM types removal (phosphorylation, methylation, S-glutathionylation, S-nitrosylation, SUMOylation and protein cleavege) are predicted to be the cause of 10 or more inheritance pathologies. Still, even low frequent PTM types such as malonylation or neddylation, with only 45 and 40 modified instances each, show associations. In terms of novelty, besides acetylation, S-glutathionylation, S-nitrosylation, O-GalNAc glycosylation, O-GlcNAc glycosylation, ADP-ribosylation and hydroxylation are the types most underestimated, with ≥50% of no previously described associations although the last ones are linked only to 4, 3 and 1 diseases respectively. Proportionally, phosphorylation is the PTM with more evidences, with only 3 novel associations from the 19 predicted (16%). Interestingly, there are a number of diseases predicted to be associated with several PTM types, with Li-Fraumeni syndrome and Pachyonychia congenita at the top with 5 associations and followed by 7 diseases with 4 associations: Amyotrophic lateral sclerosis, Hemofilia B, Hutchinson-Gildford progeria syndrome, Lissencephaly due to TUBA1A mutation, Lissencephaly type 3, Hyperinsulinism-hyperammonemia syndrome and Dilated cardiomyophathy. Still, we observed a 74% with a unique PTM type involved in their molecular causes.

**Figure 2.**
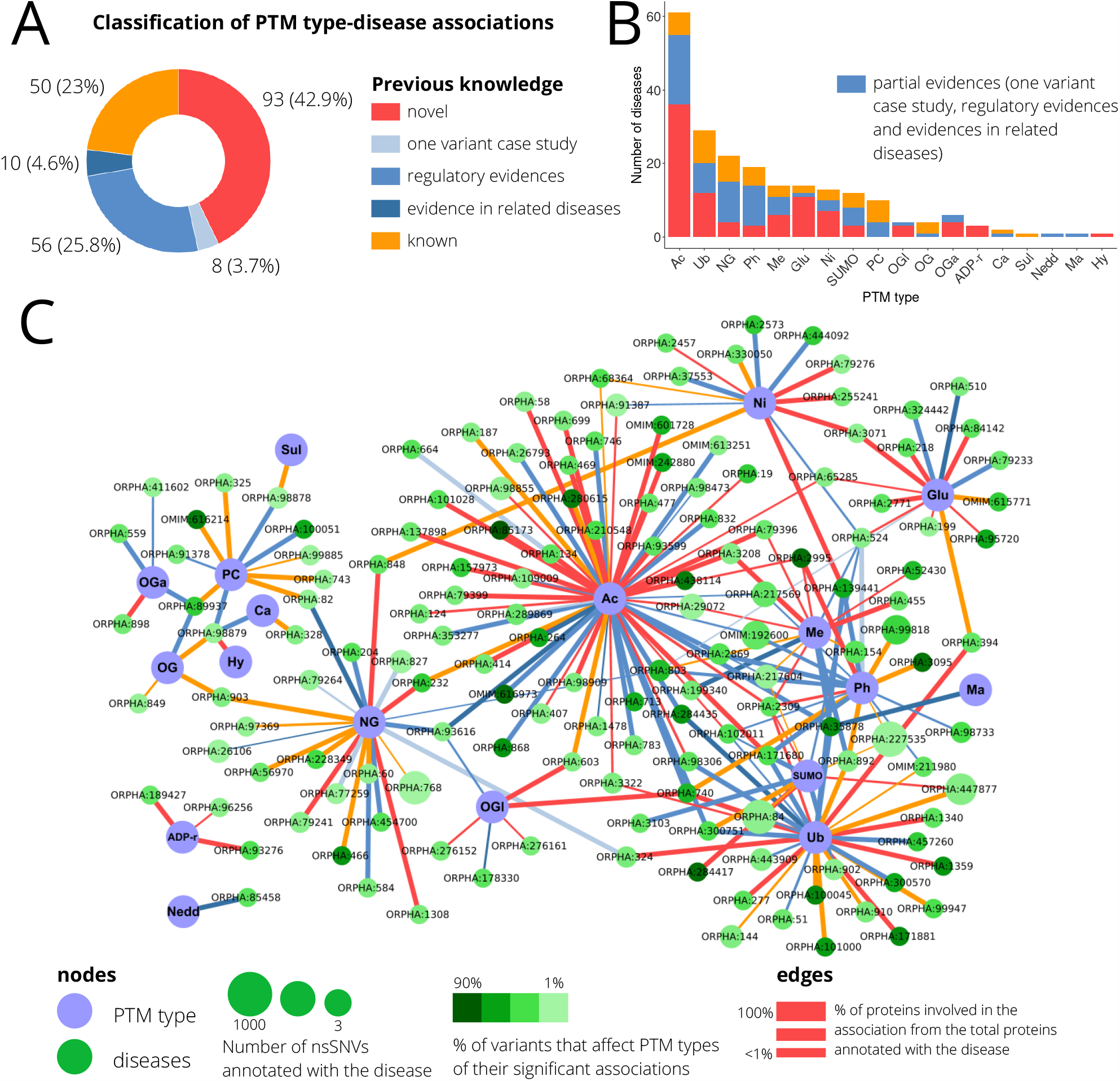
Predicted associations between PTM types and genetic diseases. A. Classification of the associations based on the current knowledge obtained by means of a literature review. B. Per each PTM type, we show the number of diseases to which associations have been predicted, classified using current knowledge. C. Network providing all predicted associations between PTM types and genetic diseases. Two types of nodes are present: PTM types and diseases, and links represent a predicted association that is colored according to its current knowledge status.

We also checked the degree in which the signal of the predictions may come from a saturation of modifications in the proteins. Thus, for each of the significant PTM type and pathology pairwise association, we randomly permuted the location of the modifications, maintaining the ratios of PTMs in ordered and disordered regions and only in modifiable residues by the PTM type. This exercise was performed 100 times, and for each, a Fisher’s exact test was calculated. We call here Occupancy Score (OS) to the number of times an association has been found significant in this process (Figure 3A), being 100 the greater value and meaning that PTMs of a type are affected by variants associated to a particular disease even if the modifications are randomly located; and 0 the smaller, meaning that the actual location of the PTMs is, almost, unique in being affected by the nsSNVs. Thus, 96 of our predicted associations (44.2%) had from very low to medium values of OS, and 121 (55.8%) high or very high OSs (Figure 3B, Figure S2). Interestingly, we observed that predicted novel associations display more robust OSs and lower numbers of PTMs and nsSNVs compared to known and those with partial evidences, p-values = 0.045 and 0.001 respectively (Figure 3C). In addition, OS levels are unequally distributed over associations in terms of the PTM types involved, with phosphorylation showing the greater level of OSs and protein cleavage the smallest, considering PTM types with >=10 associations (Figure S3).

**Figure 3.**
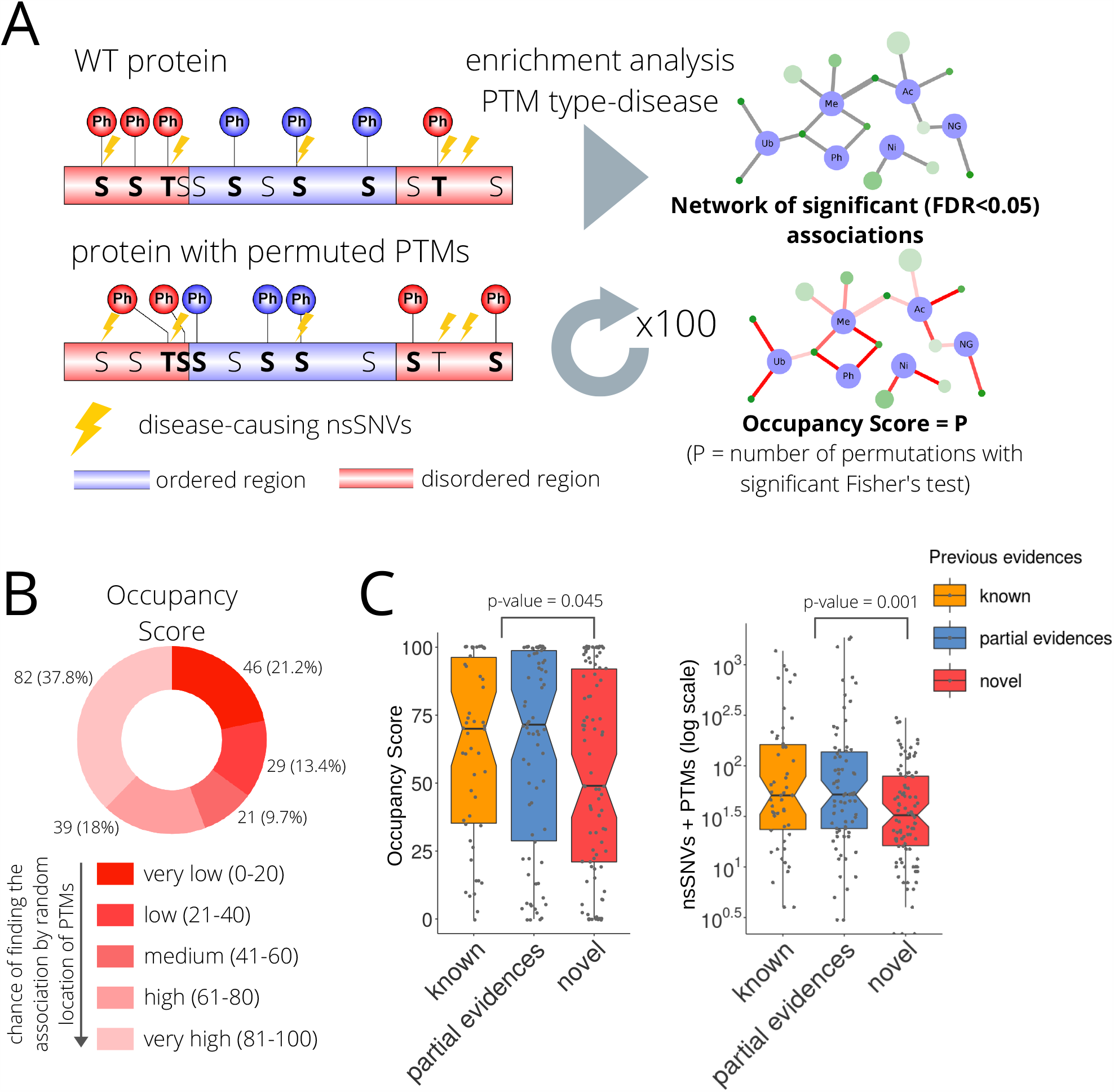
Assessment of PTM type-disease association in term of PTM occupancy. A. Schema for the calculation of the Occupancy Score (OS). B. Classification of the associations based on their OSs. C. Boxplots for: 1) OS distributions in associations annotated as known, having partial evidences, and novel; 2) for the same classification of the predicted associations: distribution of the number of nsSNVs associated to the disease and PTMs of the type being studied coinciding in the proteins annotated as involved in the disease. For the comparisons of these distributions, known and partial evidences where merged and compared to novels using Wilcoxon rank sum tests.

### Associations between the removal of PTM types and disease families and phenotypic features

We wanted to describe patterns of PTM types whose removal affect similar pathologies. Thus, we applied Orphanet disease ontology to classify diseases into families (Table S10) and use them to merge our predicted associations of PTM types-diseases. From these new connections, we calculated Jaccard index for all pairs of PTM types over the disease families that each one is linked to and arranged them into groups of PTMs sharing families (clusters, Figure S4). In Figure 4A we show the PTM types distributed according to their Jaccard index and linked to the disease families. Three main clusters of PTM types are observed. First, a core of 8 PTM types (cluster 1) formed by phosphorylation, ubiquitination, acetylation, SUMOylation, methylation, N-linked glycosylation, S-nitrosylation and S-glutathionylation, that are predicted to be behind the molecular basis of the 71% of the associations in the network (124 links out of a total of 175). This core is predicted to be responsible for the deregulation of important functions producing 88 diseases grouped in 17 families, being the skin and cardiac diseases the largest groups. Second, a cluster of 5 PTM types (cluster 2) formed by O-GalNAc glycosylation, O-linked glycosylation, proteolytic cleavage, O-GlcNAc glycosylation and ADP-ribosylation, that is connected to 11 disease families, shared with cluster 1, being neurologic diseases which groups 72 different pathologies the largest family followed by developmental defect during embryogenesis, neoplastic, endocrine, hematologic and eye diseases sorted by the number of pathologies affected. Cluster 2 also shows an exclusive link between proteolytic cleavage and allergic diseases. The third cluster of PTM types (cluster 3) is integrated by hydroxylation, sulfation and carboxylation, whose removal is predicted to affect only to hematologic diseases. Last neddylation and malonylation affect both to two disease families, neurologic and systemic or rheumatologic, and inborn errors of metabolism and endocrine, respectively.

**Figure 4.**
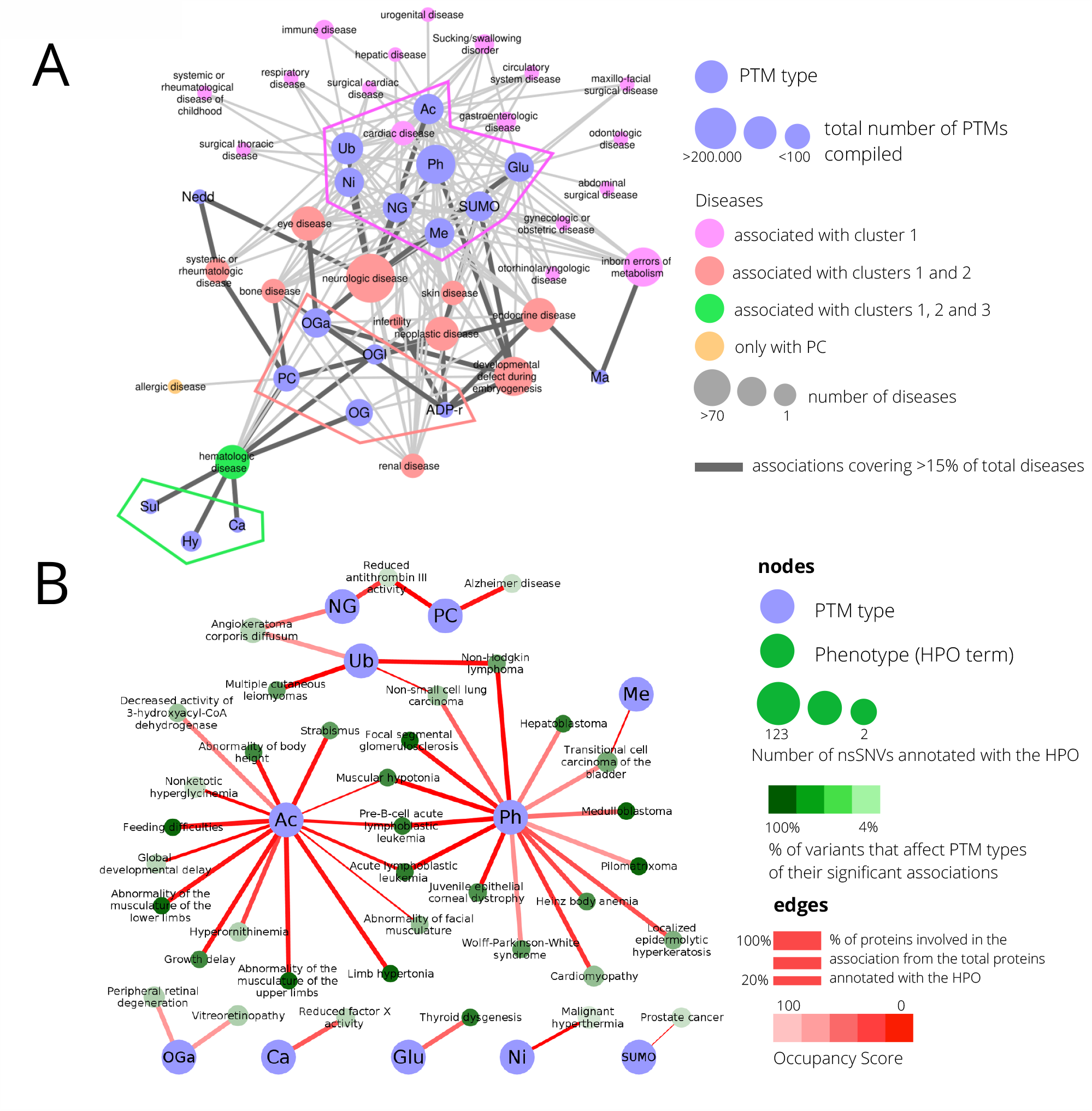
Associations between PTM types and disease families and phenotypes. A. The PTM type-disease predicted associations where merged into PTM type-disease families associations and represented in a network. Nodes are PTM types and disease families, and a edge represent a the number of predicted associations between the PTM type and disease from a particular family. PTM types are distributed according to a jaccard index calculated with the families they share. Connections based on the jaccard index between PTM types are not represented. PTM types are arranged in clusters designed based on the disease families they share and disease families are colored according to their connections to clusters. B. Network providing all predicted associations between PTM types and human phenotypes using Human Ontology Phenotype (HPO) terms. Two types of nodes are present: PTM types and HPO terms, and links represent a predicted association that is colored according to its Occupancy Score.

Next, we extracted associations between PTM types and phenotypic features taken from the Human Phenotype Ontology [53] using same methods (Table S11). Up to 133 significant associations were found between 11 PTM types and 101 HPO terms, 37 parent and 64 child terms (Table S12 and Figure S5). A global network is presented in Figure 4B with only HPO parent terms for clarity. Here the removal of acetylated and phosphorylated sites show the greater number of associations, both with 15 HPOs associated, 3 of them shared (Muscular hypotonia and two leukemia-related phenotypes). In fact the network is quite disconnected from the phenotypes point of view, phosphorylation shares 3 other cancer-related terms, 2 with ubiquitination (Non-Hodgkin lymphoma and Non-small cell lung carcinoma) and one with methylation (Transitional cell carcinoma of the bladder). The link between phosphorylation and cardyopathies comes up again although most of the terms are indeed related to cancer, same happening with ubiquitination. Last, acetylation seems to be crucial to phenotypes related to development and growth.

### Prediction of new potential mutation sites associated to diseases

Delving on our hypothesis, if there is a global association between a PTM type and a genetic disease, other modifications of the same type and proteins would become candidate sites for eventual nsSNVs to produce the same phenotype. As a proof of concept, for each of the predicted associations, we compared the pathogenicity, measured using Polyphen2 [18], that *in silico* randomly generated nsSNVs would cause in: i) sites modified by the same PTM type not affected by a variant related to the disease in the same proteins where the association has been found, and ii) sites with the same PTM type in proteins not involved in any association related to that PTM type, as the base-line pathogenicity of residues with the same PTM type, same functionality. Here, sites refer to nucleotides in codons coding for the modified amino acids. In Figure 5A we show the general schema of this comparison. In total, 37 associations between PTM types and diseases showed a significantly higher pathogenicity in PTMs not affected by a disease variant (Table 1).

**Table 1.**
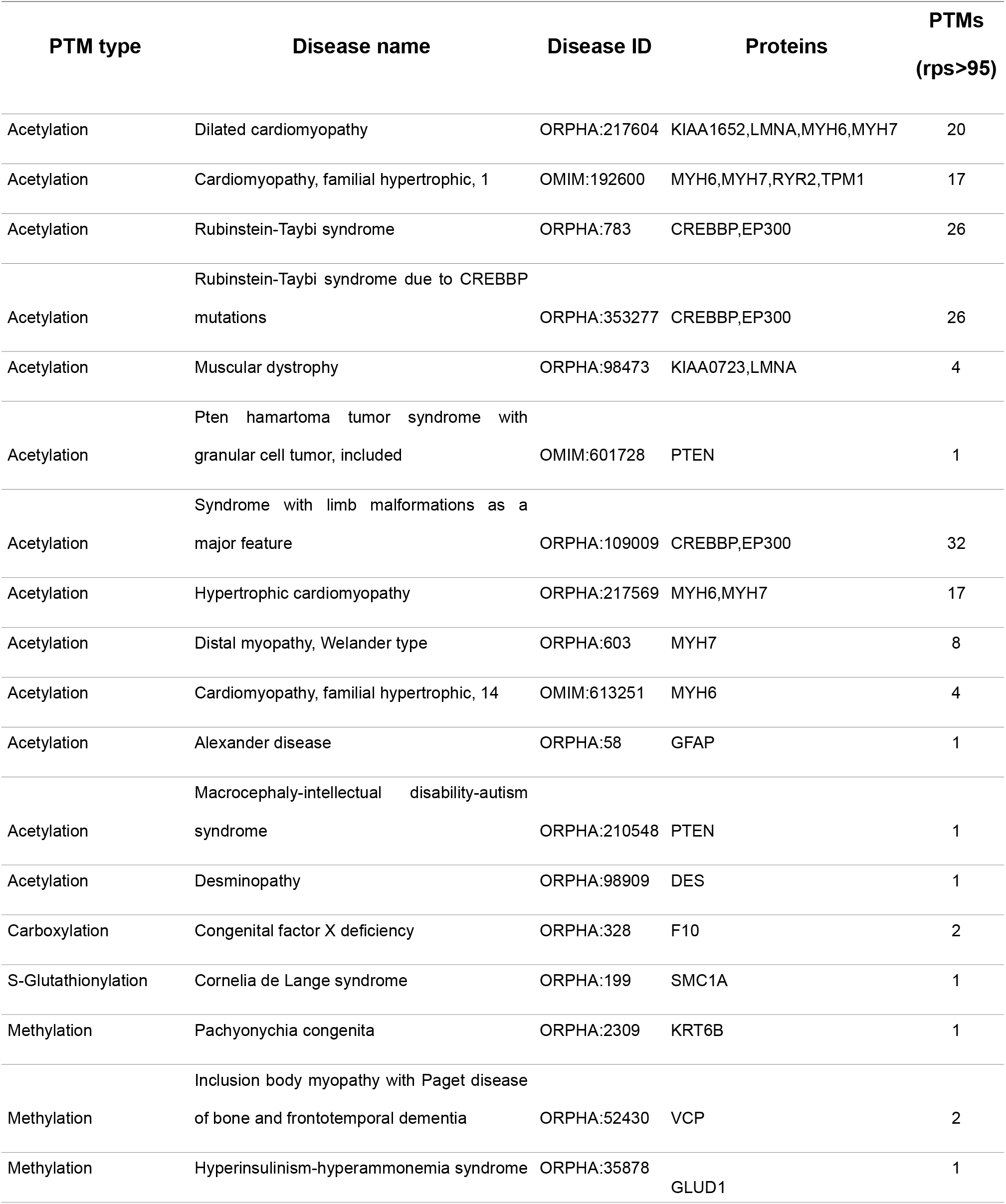
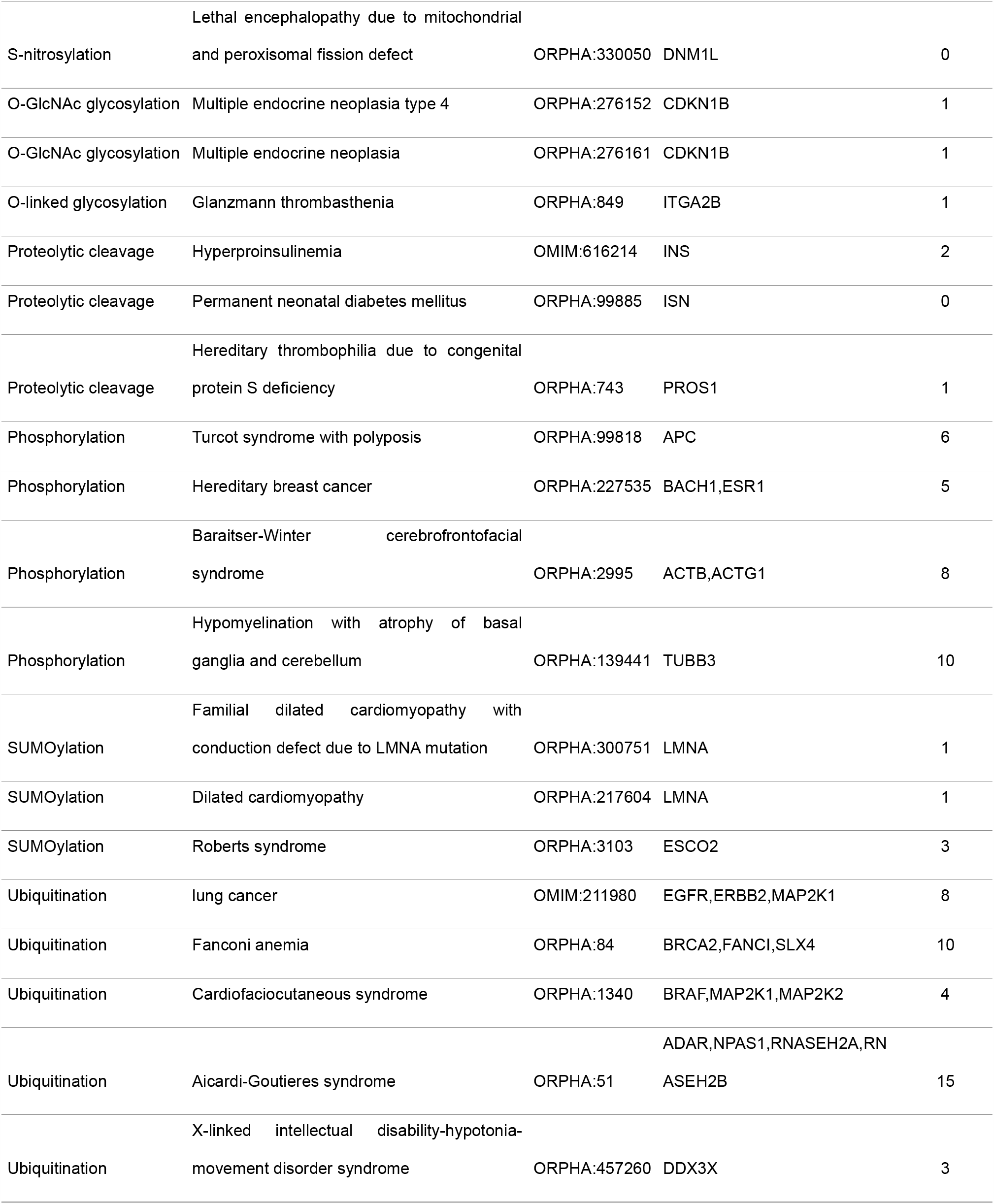
Associations between PTM types and genetic diseases where there is significant higher pathogenicity in the alteration of modified sites of the same type but not affected by the disease variants and other PTM sites. Proteins and sites predicted to be potentially involved in the disease if a genetic alteration occurs are given.

**Figure 5.**
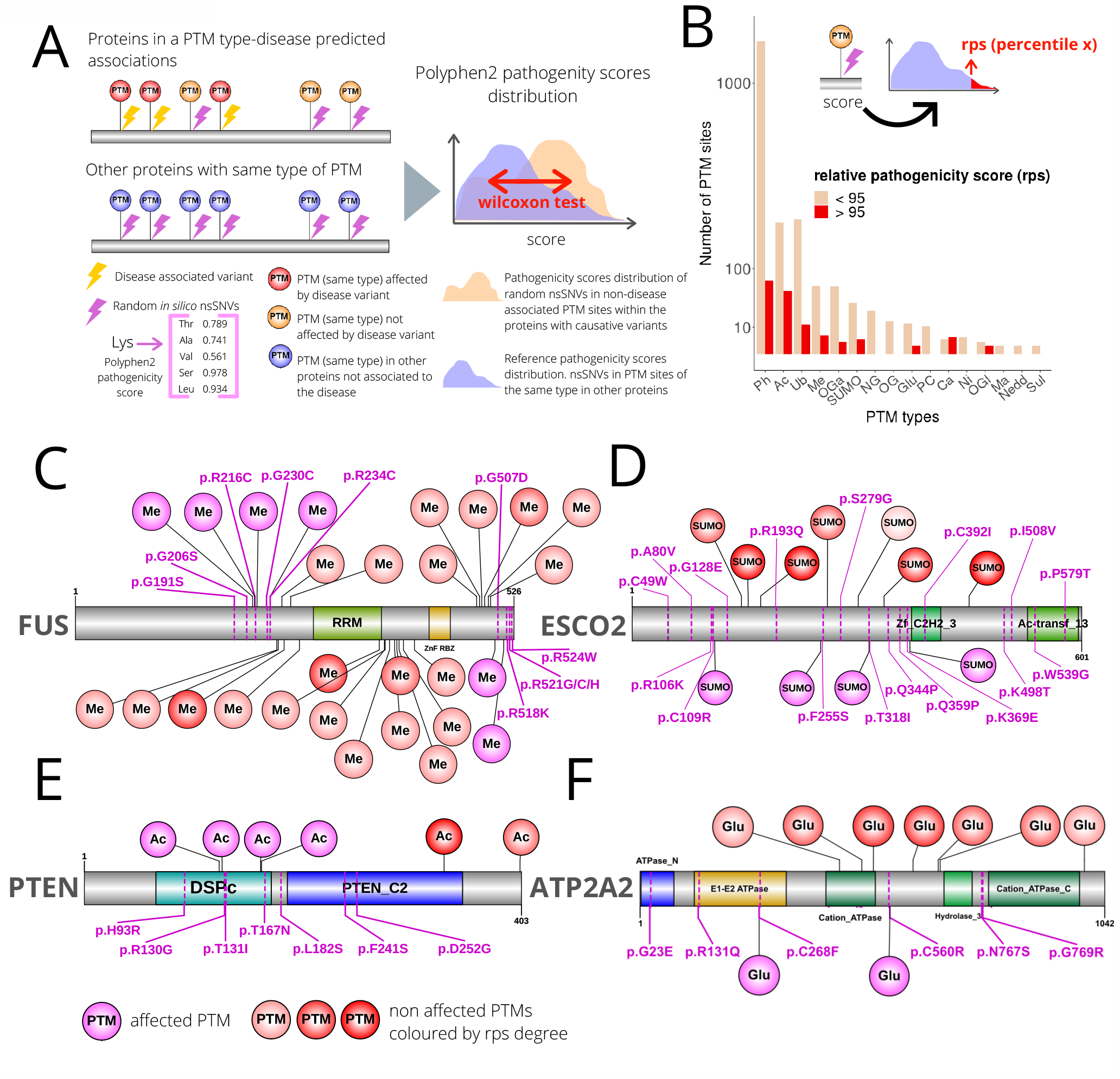
Prediction of pathogenicity for PTM sites in the context of associations between PTM types and genetic diseases. A. General schema of the comparison between: 1) the potential pathogenicity of selected PTM sites by being non affected by disease nsSNVs when the disease and its type are predicted to be associated (colored in bluish), and 2) reference distribution of the potential pathogenicity scores of PTM sites of the type but not linked to diseases (colored in orangey). B. Barplot with the number of PTM sites with pathogenicity predictions per PTM type and schema of the rps calculation. C-F. Four examples of proteins involved in predicted associations between PTM types and genetic diseases.

In order to predict new pathogenic sites associated to a disease, we introduced here the relative pathogenicity score (rps). The rps is calculated in the context of PTM type-disease associations for PTM sites of the same type but not affected by a disease variant, as: the percentile of the pathogenicity estimation of the site in a reference distribution (Figure 5B, Material and Methods), rps>95 is set as significant. Thus, we evaluated the potential pathogenicity of the deletion of 2,205 PTMs sites in 153 proteins, from them 156 sites (7.6%) had a significant rps (Table S13). Figure 5B shows the number of sites and their rps per PTM type. We see that phosphorylation is not surprisingly the type that presents a higher number of sites evaluated, still with a percentage of significant pathogenic sites below the average (5.3%). Although carboxylation and O-GlcNAc glycosylation are the types with a higher percentage of significant pathogenic sites (57,1% and 33.3% respectively), acetylation deserves again a special attention, with a large number of PTM sites evaluated (292) arises as an essential type whose deletion may produce disease-related phenotypes, with a 18.9% of sites with a significant pathogenicity. See Table S14 for all sites and their evaluation.

Last, we discuss a few examples that illustrate some of the findings of this work. Amyotrophic Lateral Sclerosis (ALS, ORPHA:803) is a neurodegenerative disease producing muscular paralysis, the most common genetic disorder due to a degeneration of motor neurons. Several genes have been reported to carry mutations deriving in this phenotype, including FUS, an RNA binding protein with genomic variations leading to the formation of stress granules. Methylation of FUS is required to bind the autophagy receptor p62 which in association with other proteins can control the aberrant accumulation of the granules [54]. Other studies have reported methylation as an important modification type in the accumulation and elimination of these toxic granules [55] supporting our predicted association with ALS (Figure 2C). In a close look, Figure 5C shows FUS with methylation sites affected and non-affected by the ALS mutations, the former with the probability of being more pathogenic than regular methylation sites, measured using rps.

Second example (Figure 5D) shows protein ESCO2, and acetyltransferase with genomic variants causing Roberts syndrome (RBS, ORPHA:3103), an ultra rare disorder yielding limb and facial abnormalities and a slow growth pre and post-natal. In RBS, the dysfunction of ESCO2 produces a lack of acetylation in the cohesin complex that regulates a proper sister chromatid cohesion in the DNA replication process [56]. We predict an association between the lack of SUMO sites and RBS. Figure 5D shows four SUMO sites in ESCO2 affected by RBS variants and seven more that might be candidates for the same type of regulation, three of them with a significant rps. A sister chromatid cohesion impairment has already been observed by a lack of SUMOylation directly in the cohesin complex [57] as well as in a knockout of SUMO protease SENP1 in chicken cell lines [58] suggesting an important role of this PTM type in this process still with no reports linking it to RBS. We annotated the association as with a partial evidence (Figure 2C).

Next we briefly describe two examples of associations with no previous evidences. Starting with the PTM type with more number of associations, acetylation, we show the phosphatase PTEN (Figure 5E) with 7 mutations linked to the Macrocephaly-intellectual disability-autism syndrome (ORPHA:210548), a rare neurological disorder with macrocephaly and facial deformity as notable anatomical features and psychomotor delay, intellectual disability and autism, as main disabilities. PTEN acts at the phosphoinositide 3-kinase pathway and it is described to be intensively regulated by acetylation [59,60]. Figure 5E shows PTEN protein with four acetylation sites affected by variants and K332 predicted to be a functionally associated site to the mentioned phenotype. In a second novel case we focus on S-glutathionylation, the eight most abundant PTM type in our collection, having almost 80% of our predictions not reported in the scientific literature (11 out of 14). One of these new predicted associations is with Darier Disease (DD, ORPHA:218), a rare skin disorder characterized by the formation of keratotic papules during puberty that are frequently subject of infections. Causative mutations have been described only in the gene ATP2A2 that codifies for protein a SERCA Ca(2+)-ATPases, responsible for endoplasmic reticulum (ER) Ca2+ uptake. Malfunctions in this protein produce an alteration of Ca2+ homeostasis in the cytoplasm that alters keratonocytes functions downstream [61]. Figure 5F shows ATP2A2 with DD variants and S-glutathionylation sites mapped, two of these residues are directly removed by the variants and we calculate a proxy score to detect the probability of other glutathionylated residues to be key sites in the disease. Although S-glutathionylation has been described to be involved in the regulation of Ca2+ ER pumps, for instance activating SERCA in arterial relaxation [62] or promoting its optimal oscillation by diamide in bovine aortic endothelial cells [63], to the best of our knowledge this is the first report suggesting an association with the disorder.

## Discussion

There are two motivations in the study of PTMs in the context of genetic diseases, first and most obvious is to contribute to deepen into the genetic basis of these pathologies. Systematic and global approaches are of special interest in rare diseases, where resources hardly cover a small percentage of them due to their individual low incidence but research is encouraged as affecting together around 7% of the population [64]. Here we have studied >3,200 genetic diseases in sync and extracted significant results for 150 with new hypothesis linking the loss of specific functions to diseases and phenotypes as well as describing general patterns of associations. The second motivation comes to add functional information to individual PTMs [3,65,66], frequently extracted from high-throughput experiments and most with succinct annotation [36]. The annotation of PTMs, the study of the types as functional entities, and their interactions, is essential to decode the regulation of proteins and how cell responds to a changing environment, named the PTM code [5]. Although several previous resources have crossmatched genomic variation and PTM data [4,12,13], for the first time we add a global interpretation of PTM types in the context of genetic diseases as well as prediction capacities, even to the less studied PTM types.

In terms of dimensionality, we believe we present herein the largest study crossmatching human genomic variation and protein modifications, comprising 4,464,412 germline nsSNVs (44,969 involved in diseases and 392,185 predicted pathogenic) and 321,643 of 59 types that revealed 1,743,207 PTM-nsSNVs co-occurrences in 16,541 proteins. Of those PTMs and nsSNVs co-occurrences, 15,607 involved diseases and 133,292 pathogenicity. Before us, Reimand and colleagues [10] focused on four PTM types (130,000 sites) reporting 77,829 matches of PTM regions with nsSNVs. Kim et al. [67] extended the number of PTM types to 20, extracting 179,325 PTM-SNVs associations using 517,466 missense and stop gain SNVs and 599,387 PTMs, most of them predicted (404,501) which are generally reported with low accuracy [68]. More recently, the AWESOME database [14] also provides a large dataset with 1,043,668 nsSNVs and experimentally verified and predicted PTMs of six types that sum up 481,557 matches. Other approaches focused as us on diseases associations, but compiling curated associations from the scientific literature, thus PTMD [13] and PhosphoSitePlus [4] for instance report 1,950 and 1,230 PTMs-SNVs associations respectively. The topic is clearly of interest [69].

Our framework produced a total of 217 predicted associations between the loss of a function provided by a PTM type and a genetic mendelian disease. Although other types of genomic variations are reported to affect functions led by PTMs [70], we concentrated in reported causal nsSNVs for each of the disorders. Since the global diagnosis rate of these disorders is about 50%, and the numbers of predicted pathogenic variants exceed substantially the numbers for annotated disease causing variants (Figure 1B), it is feasible to state that the scenario described here is underestimated. The incomplete repertoire of protein modifications is an additional limitation of this type studies [34]. Other sources of implications of PTMs in harmful phenotypes are for instance the possibility of genetic variants to produce new modifiable sites or PTM motifs [11,14], or the alteration of the enzymatic cascades that attach the PTMs, which we have used as confirmatory rule (Figure 2A, 2C). Regarding PTM functionality, all systematic studies compiling their information should aware of the cell state dependence of their occupancy [15].

This wealth of information has been scanned in two different manners. First we report an extensive literature review where we extracted known associations between PTM types and diseases (23% of the total), and evidences not linking directly the PTM type to the disease but showing related functional proofs (34%). Altogether they represents a 57% of a our associations which grant a good endorsement of our approach and still provides new insights for about 167 associations, 93 completely novel. We are aware that with a literature review of such a magnitude, there might be overlooked associations, still we believe they would fall into partial evidences as they are less obvious links. Additionally, we checked if the associations are extracted only due to a high level of PTM occupancy using a permutation approach as others [71]. Our Occupancy Score measures this possibility (Figure 1A) showing up to 44% of the pairwise predictions to have PTMs displayed in proteins with a medium to high chance of being the unique conformation providing the association (Figure 1B). Importantly, PTM occupancy is lower in the novel predicted associations and higher in PTM types like phosphorylation, ubiquitination or SUMOylation which might indicate that the discovery of associations depends at least partially on the level of knowledge accumulated on PTM types and proteins/diseases. Thus, potential associations involving low studied proteins or PTM types with little proteome coverage seem to be more probable to be missed unless systematic studies like this one are performed. Stressing this idea, we found for instance phosphorylation having a great proportion of the predicted associations with some kind of evidences, only three completely novel.

Summing up the findings, at the top of the PTM types we highlight acetylation with not only histone functions [72] predicted to be involved in up to 61 diseases, almost 60% of them no previously reported, linked to several phenotypes, 12 of them exclusively, and with a wide range of OS values. Its implication seems to be relevant in several disease families, with neurologic and inborn errors of metabolism summing 19% and 14% of its associations but also being connected to urogenital, hepatic and immune diseases almost alone. Other PTMs seems to be of special interest, of course ubiquitination due to its role in protein degradation which deregulation is described widely associated to human pathologies [25], but also S-glutathionylation, S-nitrosylation or the different glycosylations, that are normally not included in the systematic studies but with a wealth of predictions here. Displayed in a network format and merging diseases in families we were able to extract general patterns that provides clues for researches to explore PTM types implications in diseases that do not have enough genetic information to be explored. Similarly, a network of links between PTMs and phenotypes is also produced. In this sense, it is expected that disorders with similar phenotypes are caused by similar deregulation processes [73]. Last about network analysis capacities, it is feasible to think that the convergence of several PTM types implicated in similar diseases, might be a signal for their functional crosstalk [3,74].

Finally, if our strategy until this point has been to try to infer general patterns from individual events (PTM-nsSNV co-occurrences) we wanted to give an extra value to the predictions scaling down the functional implications and providing a bunch of PTM sites whose removal by genomic variations are predicted to produce specific disorders. The score we built for this task, the rps, is quite conservative in the sense of making validated pathogenicity scores [18] being referenced to functional sites that are expected to be pathogenic themselves [75]. Besides the four examples discussed in detail, we add as Table S14 all the PTM sites predicted to be functionally linked to the diseases, with detailed Polyphen2 scores and rps values. This, together with all the PTM-nsSNV co-occurrences available as Supplementary Data 1 (>1.7M) represents a valuable and unique source of information to be included in bioinformatics pipelines of DNASeq analysis to annotate genomic variants. Since, the filtering and prioritization of variants is an essential task in genetic diagnosis based on deep sequencing [76], we believe the resources reported in this work may contribute to create new hypothesis on specific variants that could impact in more conclusive diagnosis of patients.

## Material and Methods

### Compilation of nsSNVs and annotation as disease associated and pathogenic

We compiled human nsSNVs from the following databases: dbSNP [77], gnoMAD v2 [44,78], Humsavar (UniProt) [39], ClinVar [45], and DisGeNET [46]. Protein ids were mapped to STRING v10 [79] dictionary and sites were mapped into eggNOG v4 [80] set of human sequences confirming the wild type amino acid.

The disease annotation of nsSNVs were retrieved from Humsavar [39], ClinVar [45] and DisGeNET [46] databases as OMIM [81] and Orphanet [52] diseases. Phenotype information was retrieved from ClinVar as Human Phenotype Ontology terms (HPO) [82]. OMIM and Orphanet ids for the same diseases were merged into Orphanet ids.

To assess the pathogenicity of the nsSNVs, we used pathogenicity predictions from five algorithms: PolyPhen2 [18], SIFT [47], PROVEAN [48], FATHMM [49], and MutationTaster [50], all calculated using the Variant Effect Predictor pipeline [83]. A nsSNV is classified as “pathogenic” if >50% of the predictions are deleterious.

### Compilation of PTMs

Human experimentally verified PTM sites were obtained from: PTMCode v2 [34], dbPTM3 [84], PhosphoSitePlus [85], PhosphoELM [86], PHOSIDA [36], PhosphoGrid [87], OGlycoBase [38], UniProt [39], HPRD [88], RedoxDB [40], dbGSH [41] and dbSNO [42]. Protein ids were mapped to STRING v10 [79] dictionary and PTM sites were mapped into eggNOG v4 [89] set of human sequences as in [34]. PTM types were annotated.

### Co-occurrences between PTMs and nsSNVs

We identified nsSNVs and PTMs matching same protein positions. In addition, we identified the PTM types with enzymatic motifs in the target proteins: phosphorylation, ubiquitination, acetylation, SUMOylation, methylation, N-linked glycosylation, S-glutathionylation, S-nitrosylation, O-GalNAc glycosylation, O-linked glycosylation, O-GlcNAc glycosylation, neddylation, malonylation [90–98]. A consensus of window of ±5 amino acids from the PTM was agreed to consider the nsSNV matching motifs.

### Enrichment of disease associated and pathogenic nsSNVs affecting PTMs in PTM types

We extracted ordered and disordered regions in every protein using MobiDB [99]. PTMs-nsSNVs co-occurrences were assigned to ordered or disordered regions.

We measured if the set of nsSNPs affecting specific PTM types are enriched in disease causing nsSNPs. Thus, we counted: disease-nsSNVs matching PTM sites, disease-nsSNVs not matching PTM sites, non-disease-nsSNVs matching PTM sites and non-disease-nsSNVs not matching PTM sites, applied a Fisher’s Extact test and p-values adjusted by Benjamini-Hochberg False Discovery Rate (FDR). FDR < 0.05 was considered significant. The same analysis was performed for predicted pathogenic nsSNPs.

### Extraction of associations of PTM types and diseases or phenotypes

We measured if there are associations between specific PTM types and specific genetic diseases. For each PTM type and genetic disease in our compilation, we counted: nsSNVs causing the disease affecting the PTM type, nsSNVs causing other diseases affecting the PTM type, nsSNVs causing the disease not affecting the PTM type and nsSNVs causing other diseases not affecting the PTM type. We applied Fisher’s exact test and p-values adjusted by FDR. FDR < 0.05 was considered significant. The same analysis was performed for nsSNPs affecting phenotypes annotated as HPO terms.

### A literature review to classify the predicted PTM type and disease associations based on current knowledge

We performed a literature review to catch the current knowledge for each of the 217 predicted associations between PTM types and genetic diseases. Due to the volume of the combinations (>190K) it was not possible to calculate false negatives. The searches were performed with the two terms (PTM type and disease) in PubMed and Google considering derived and related terms if appropriate. The associations were classified as: i) known, if at least a scientific article describes the implication of the loss of the PTM type in the disease; ii) having partial evidences, or iii) novel, if no information is found linking both terms. Within partial evidences we include: i) reports of one variant case, where a single nsSNV, if more than two are described, is reported to prevent the modification of the residue; ii) associations of the PTM type with a related or similar disease; and iii) regulatory evidences, where at least a scientific work reports that the disease is produced by a deregulation involving the PTM type within the regulatory neighborhood of the proteins with the deletereous genomic variations. PubMed ids are provided for all the cases with some kind of evidence, if multiple are found we provide a selection.

### Occupancy Score calculation

To evaluate whether the predicted associations are due to a high occupancy of PTMs of a type in the proteins with variants annotated with the disease, we implemented the Occupancy Score (OS). OS is calculated for each predicted association as follows: 1) Same number of PTMs of the selected type are randomly located in modifiable residues of the PTM type, maintaining their ratio of ordered and disordered regions; 2) new co-occurrences of PTMs and nsSNVs associated to the disease are extracted; 3) same Fisher’s Exact test described above is performed and p-values adjusted; 4) steps 1, 2 and 3 are performed 100 times; 5) OS is assigned to the association as the number of times the association is found significant. The comparisons between OS distributions were performed by means of Wilcoxon rank sum tests.

The same methodology was applied to calculate OSs for phenotype-PTM type associations.

### Classification of diseases in families

We used Orphanet disease families to classify the 150 disorders that had at least a significant association with a PTM type. OMIM diseases that did not have a correspondence to Orphanet diseases were manually mapped to the families.

### Jaccard index calculation

We merged the associations of PTM types and diseases into supra association between PTM types and disease families. We calculated jaccard index [100] of every pair of the 18 PTM types with at least a significant association. We built a network with only PTM types as nodes and edges annotated with their jaccard index that was uploaded into Cytoscape [101] and Prefuse Force Directed layout applied on the jaccard index. We uploaded into Cytoscape the network between PTM types and disease families maintaining the conformation of PTM types produced by their similarity. Clusters of PTM types were created according to their sharing disease families.

### Pathogenicity comparison of PTMs of a type in proteins causing the disease but not affected by variants and PTMs of the same type in other proteins

For each of the predicted associations between PTM types and genetic diseases we extracted the PTMs of the selected type that are not affected by the nsSNPs causing the disease. For them, we randomly produced five non synonymous amino acid *in silico* changes (mimicking the effect of nsSNPs), measured their pathogenicity using Polyphen2 [18] and calculating their mean. The distribution of these means was compared to a reference distribution composed by the same values for PTMs of the same type in proteins not associated to any disease where the PTM type is predicted to be involved. The comparison was made using Wilcoxon rank sum tests and p-values adjusted by FDR. FDR<0.05 was taken as significant.

### Relative pathogenicity score (rps) calculation

For a PTM site with suspicion to be associated to a disease, a rps can be calculated as: 1) production of five random non synonymous amino acid *in silico* changes; 2) get Polyphen2 score for them; 3) calculate the mean; 4) calculate the percentile of the mean in the reference distribution (see section above). Thus, the rps ranges from 0 to 100. Rps >95 was considered significant.

## Acknowledgments

We wish to thank members of the Bioinformatics Unit and the Genetics Department of the IIS-FJD for their comments and critical review of the work. We also thank IIS-FJD and ISCIII for their support and the RAREGenomics network for providing a forum of discussion.

## Author information

### Contributions

PV collected the data, designed and performed the analyses, interpreted the data and reviewed the manuscript. PM conceived the project, designed and performed the analyses, interpreted the data and wrote the manuscript. The authors read and approved the final manuscript.

## Supplementary Information

### Availability of data and materials

The datasets supporting the conclusions of this article are included within the article and its additional files.

### Additional file 1

**Table S1. Dataset of Post Translational Modifications (PTMs), nsSNVs and genetic diseases**. Total numbers of the datasets compiled for: protein post-translational modification sites in humans, human nsSNVs, genetic diseases from OMIM and Orphanet databases, phenotypes from the Human Phenotype Ontology database, and nsSNVs with disease and phenotype annotations as well as predicted as pathogenic.

**Table S2. Disease-related nsSNVs involved in disease families**. Orphanet disease families classification was adopted.

**Table S3. Dataset of PTMs classified by type**. Number of human experimentally verified PTMs compiled, grouped by PTM types and sorted by abundance.

**Table S4. Mapping of nsSNVs and PTM sites**. Human PTMs and nsSNVs co-occurrences. A co-occurrence is considered when the nsSNV affects the modifiable residue or is located within a ±5 amino acids from the PTM site.

**Table S5. Crossmatching of PTM sites and disease-related nsSNVs**. Number of disease-related nsSNVs and PTM sites matched grouped by PTM type. We include number of proteins and diseases affected. PTMs with no matches are not shown.

**Table S6. Crossmatching of PTM sites and predicted pathogenic nsSNVs**. Number of predicted pathogenic nsSNVs and PTM sites matched grouped by PTM type. We include number of proteins and diseases affected. PTMs with no matches are not shown.

**Table S7. Data for the enrichment of disease-related nsSNVs in PTM types**. Data and results for Figure 1D.

**Table S8. Data for the enrichment of predicted pathogenic nsSNVs in PTM types**. Data and results for Figure 1D.

**Table S9. PTM types and genetic diseases predicted associations**. Together with p-values and FDR, Occupancy scores (OS) are also shown. Next to last columns holds the category related to the previous knowledge of the association, three classes: known, previous evidences, divided into one variant case study (OVCS), evidences in related diseases (ERD) and regulatory evidences (RE), and novel. Last column shows the PubMed Ids of the scientific papers supporting the classification.

**Table S10. Classification of diseases in families according to Orphanet classification**.

**Table S11. Crossmatching of PTM sites and phenotype-related nsSNVs**. Number of phenotype-related nsSNVs (form ClinVar and using HPO terms) and PTM sites matched grouped by PTM type. We include number of proteins and phenotypes affected. PTMs with no matches are not shown.

**Table S12. Data to build a network of PTM types and phenotypes associations**. Together with the p-values and FDRs, Occupancy scores (OS) are also shown.

**Table S13. Prediction of new potential pathogenic PTM sites functionally associated to specific diseases**. For every predicted association between PTM types and genetic diseases, we show the p-values and FDRs for the tests comparing pathogenicity predictions of *in silico* nsSNVs in PTMs of the same type but not affected by a disease-related nsSNV and pathogenicity predictions of *in silico* nsSNVs affecting PTMs of the same type not predicted to be involved in diseases. We add the number of PTM sites predicted to be involved in the disease and those that are not, based on relative pathogenicity scores (rps).

### Additional file 2

**Table S14. Evaluation of the pathogenicity of nsSNVs in PTM sites in the context of associations between PTM types and genetic diseases**. Grouped by PTM type and per each association with a genetic disease, we show the PTM sites (protein and position), the Polyphen2 score as the mean of the scores of 5 random *in silico* nsSNVs, and its relative pathogenicity score (rps) that represents the percentile within the reference distribution of Polyphen2 scores build by 5 random nsSNVs in same type of PTM types outside the proteins involved in the disease.

### Additional file 3

**Figure S1.**
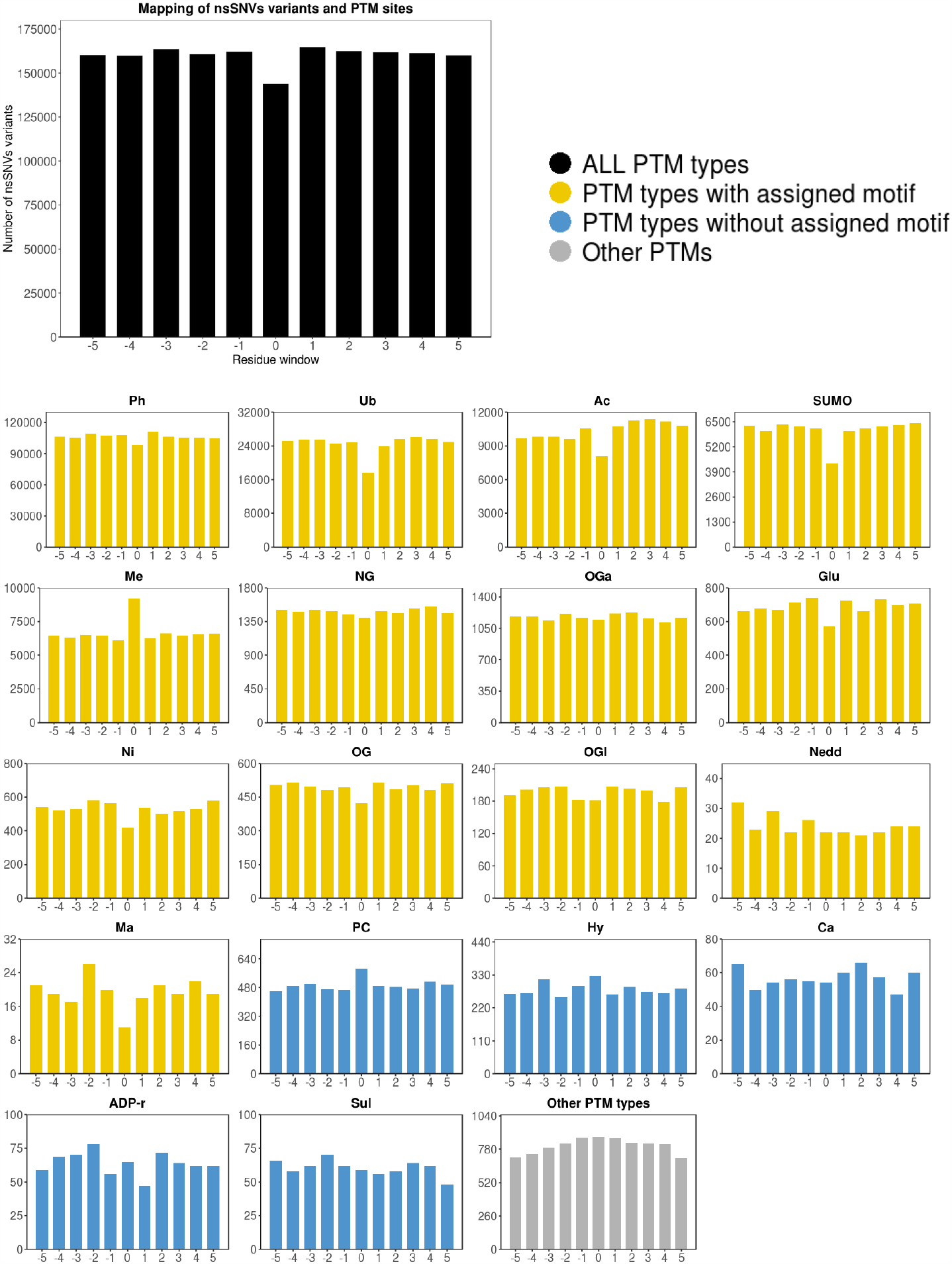
Distribution of nsSNVs in PTM regions classified by PTM type.

### Additional file 4

**Figure S2.**
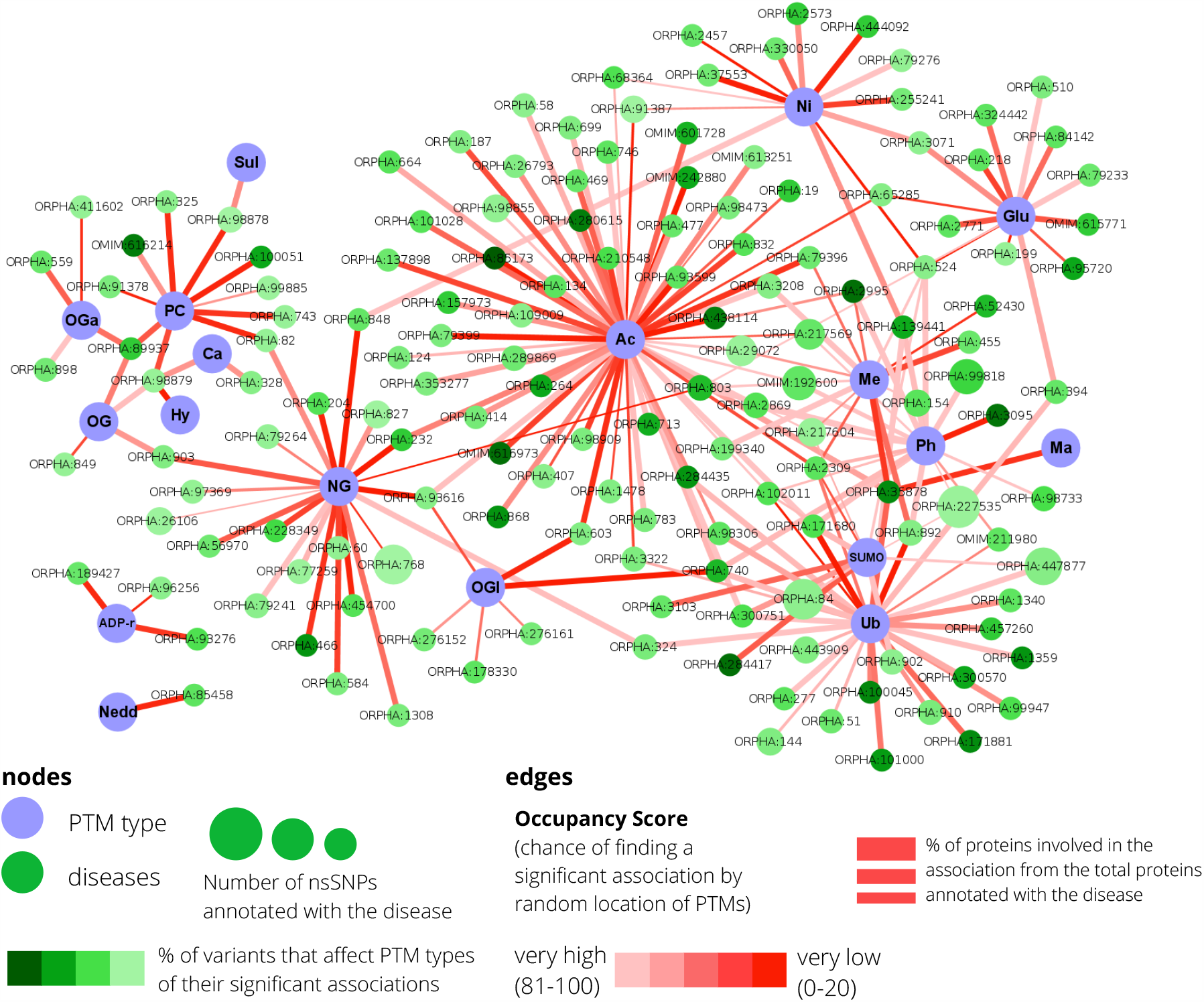
Predicted associations between PTM types and genetic diseases. Network providing all predicted associations between PTM types and genetic diseases. Two types of nodes are present: PTM types and diseases, and links represent a predicted association that is colored according to its Occupancy Score (OS).

### Additional file 5

**Figure S3.**
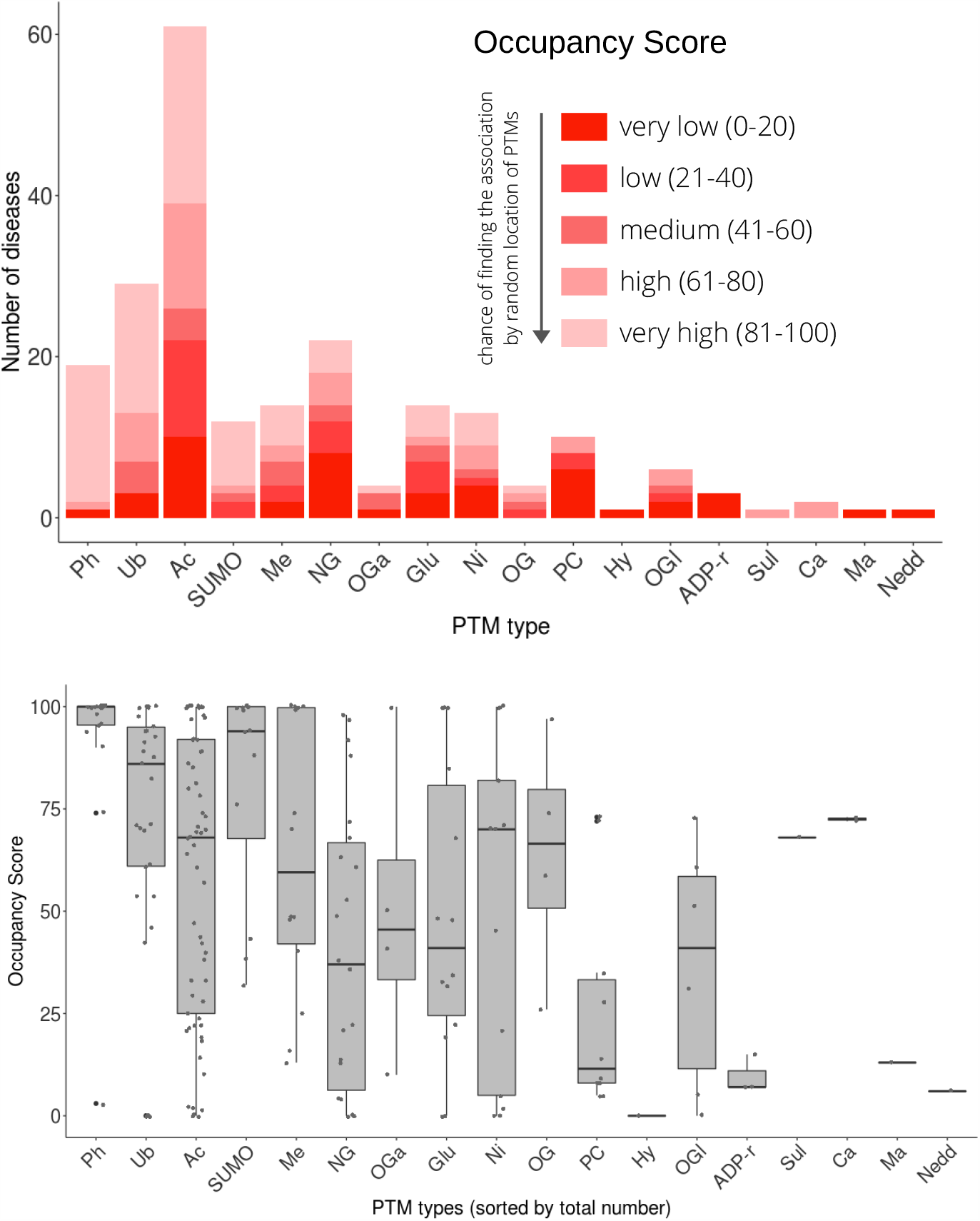
Occupancy scores in associations between PTM types and genetic disease grouped by PTM type.

### Additional file 6

**Figure S4.**
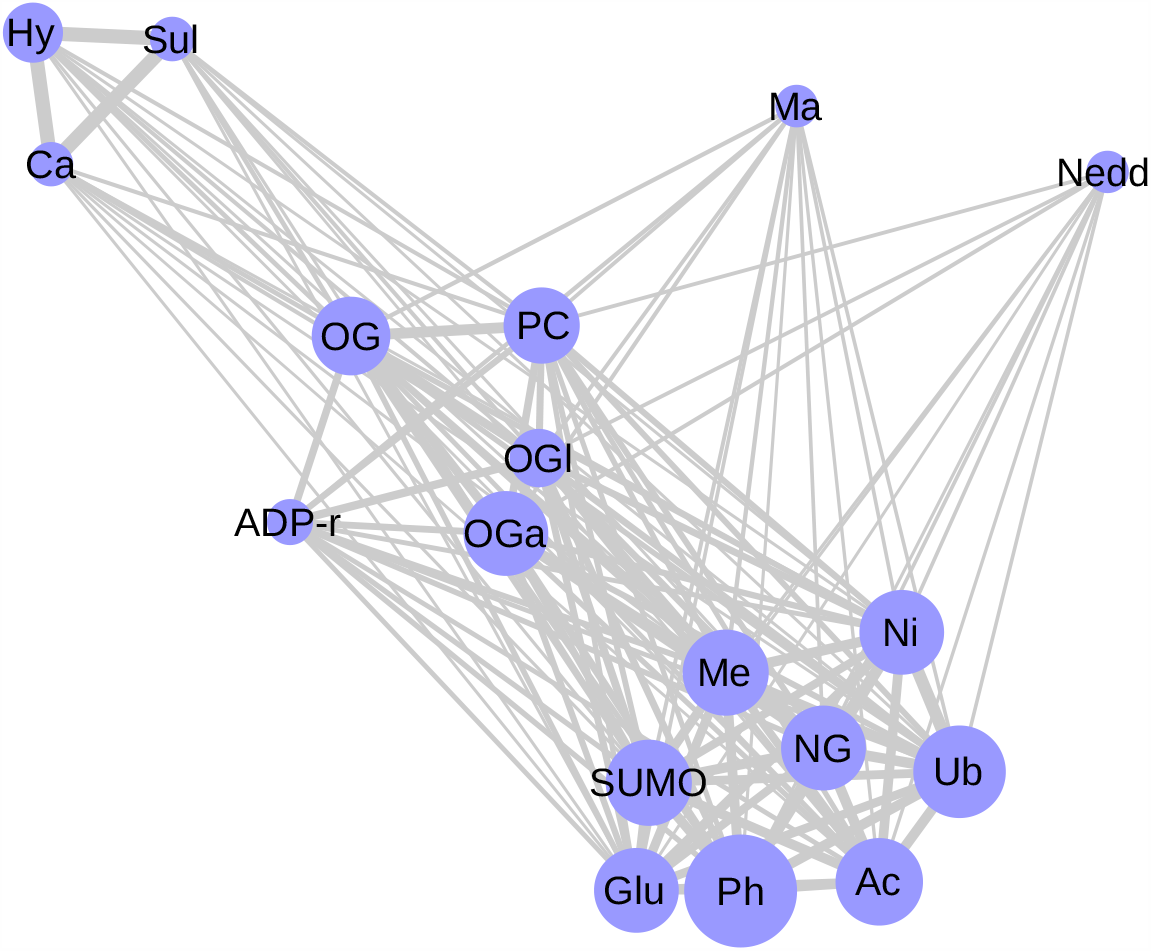
Network of PTM types linked by their jaccard index calculated by their connection to disease families.

### Additional file 7

**Figure S5.**
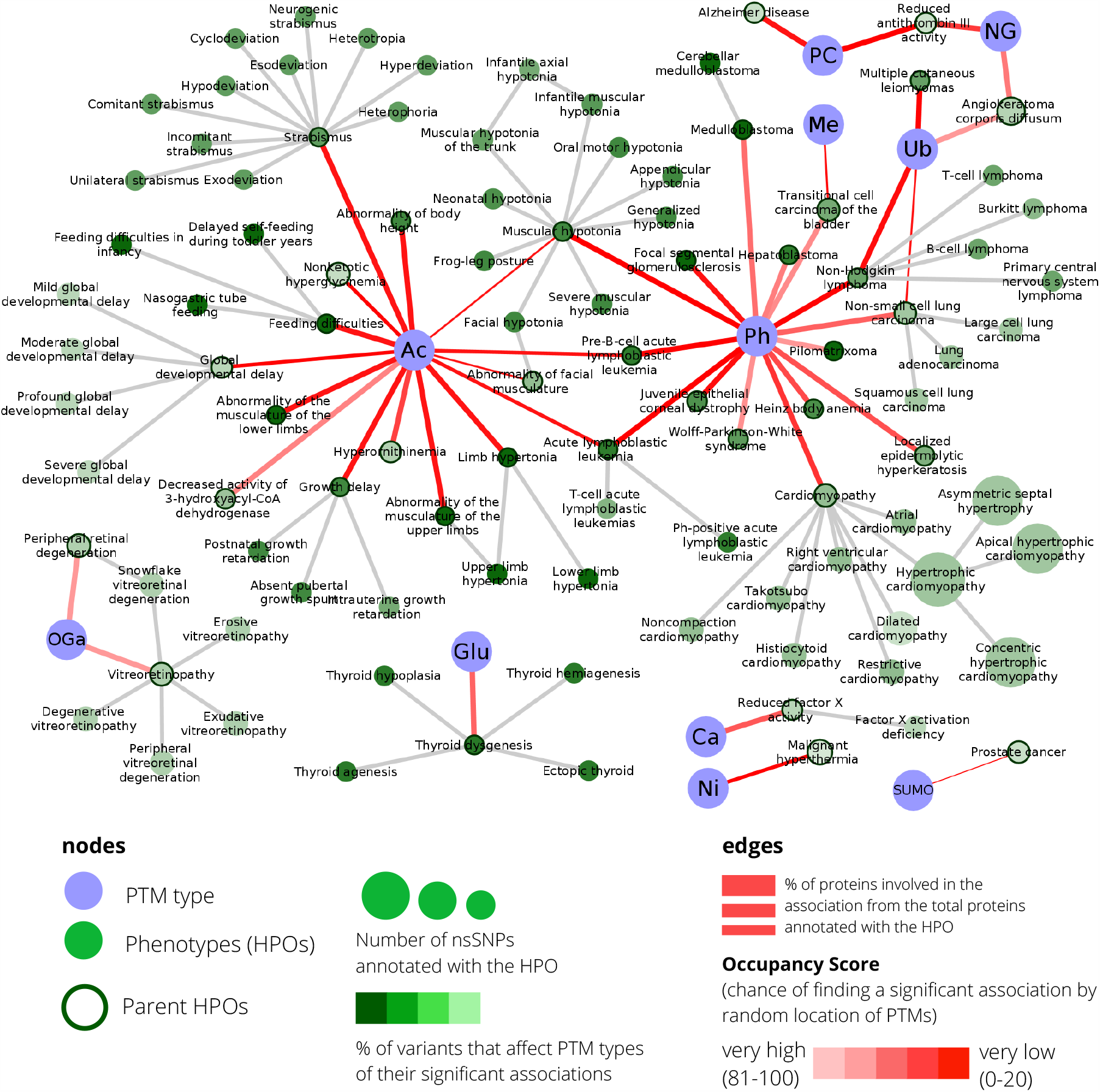
Predicted associations between PTM types and human phenotypes. Network providing all predicted associations between PTM types and human phenotypes using Human Ontology Phenotype (HPO) terms. Two types of nodes are present: PTM types and HPO terms, and links represent a predicted association that is colored according to its Occupancy Score.

### Additional file 8

**Supplementary Data 1**. Human PTMs and nsSNVs co-occurrences within a window of ±5 amino acids from the modifiable residue.

## References

1. Larsen MR, Trelle MB, Thingholm TE, Jensen ON. Analysis of posttranslational modifications of proteins by tandem mass spectrometry. Biotechniques. 2006; 790–8.

2. Buuh ZY, Lyu Z, Wang RE. Interrogating the Roles of Post-Translational Modifications of Non-Histone Proteins. J. Med. Chem. 2018; 3239–52.

3. Minguez P, Parca L, Diella F, Mende D, Kumar R, Helmer-Citterich M, et al. Deciphering a global network of functionally associated post-translational modifications. Mol Syst Biol. 2012; 8:599.

4. Hornbeck P V, Zhang B, Murray B, Kornhauser JM, Latham V, Skrzypek E. PhosphoSitePlus, 2014: mutations, PTMs and recalibrations. Nucleic Acids Res. 2015; 43:D512–20.

5. Hunter T. The age of crosstalk: phosphorylation, ubiquitination, and beyond. Mol Cell. 2007; 28:730–8.

6. De La Fuente L, Arzalluz-Luque Á, Tardáguila M, Del Risco H, Martí C, Tarazona S, et al. TappAS: A comprehensive computational framework for the analysis of the functional impact of differential splicing. Genome Biol. 2020; 21(1):119

7. Radivojac P, Baenziger PH, Kann MG, Mort ME, Hahn MW, Mooney SD. Gain and loss of phosphorylation sites in human cancer. Bioinformatics. 2008; 24:i241–7.

8. Reimand J, Wagih O, Bader GD. The mutational landscape of phosphorylation signaling in cancer. Sci Rep. 2013; 3:2651

9. Narayan S, Bader GD, Reimand J. Frequent mutations in acetylation and ubiquitination sites suggest novel driver mechanisms of cancer. Genome Med. 2016;8(1):55.

10. Reimand J, Wagih O, Bader GD. Evolutionary constraint and disease associations of post-translational modification sites in human genomes. PLoS Genet. 2015; 11:e1004919.

11. Krassowski M, Paczkowska M, Cullion K, Huang T, Dzneladze I, Ouellette BFF, et al. ActiveDriverDB: Human disease mutations and genome variation in post-translational modification sites of proteins. Nucleic Acids Res. 2018; 46:D901–10.

12. Huang KY, Lee TY, Kao HJ, Ma CT, Lee CC, Lin TH, et al. DbPTM in 2019: Exploring disease association and cross-Talk of post-Translational modifications. Nucleic Acids Res. 2019; 47:D298– 308.

13. Xu H, Wang Y, Lin S, Deng W, Peng D, Cui Q, et al. PTMD: A Database of Human Disease-associated Post-translational Modifications. Genomics, Proteomics Bioinforma. 2018;16:244–51.

14. Yang Y, Peng X, Ying P, Tian J, Li J, Ke J, et al. AWESOME: A database of SNPs that affect protein post-translational modifications. Nucleic Acids Res. 2019; 47:D874–80.

15. Wagih O, Galardini M, Busby BP, Memon D, Typas A, Beltrao P. A resource of variant effect predictions of single nucleotide variants in model organisms. Mol Syst Biol. 2018; 14:14:e8430.

16. Peng D, Li H, Hu B, Zhang H, Chen L, Lin S, et al. PTMsnp: A Web Server for the Identification of Driver Mutations That Affect Protein Post-translational Modification. Front Cell Dev Biol. 2020; 8:1330.

17. Amann T, Schmieder V, Faustrup Kildegaard H, Borth N, Andersen MR. Genetic engineering approaches to improve posttranslational modification of biopharmaceuticals in different production platforms. Biotechnol Bioeng. 2019; 116:2778–96.

18. Adzhubei I, Jordan DM, Sunyaev SR. Predicting functional effect of human missense mutations using PolyPhen-2. Curr Protoc Hum Genet. 2013; Chapter 7:Unit7.20

19. Kumar P, Henikoff S, Ng PC. Predicting the effects of coding non-synonymous variants on protein function using the SIFT algorithm. Nat Protoc. 2009; 4:1073–81.

20. Barber KW, Rinehart J. The ABCs of PTMs. Nat Chem Biol. 2018;14:188–92.

21. Perluigi M, Barone E, Di Domenico F, Butterfield DA. Aberrant protein phosphorylation in Alzheimer disease brain disturbs pro-survival and cell death pathways. Biochim. Biophys. Acta -Mol. Basis Dis. 2016 1871–82.

22. Wieland T, Attwood P V. Alterations in reversible protein histidine phosphorylation as intracellular signals in cardiovascular disease. Front Pharmacol. 2015; 6:173.

23. Kondo Y, Shen L, Issa J-PJ. Critical role of histone methylation in tumor suppressor gene silencing in colorectal cancer. Mol Cell Biol. 2003; 23:206–15.

24. Arif M, Senapati P, Shandilya J, Kundu TK. Protein lysine acetylation in cellular function and its role in cancer manifestation. Biochim Biophys Acta. 2010; 1799:702–16.

25. Jiang YH, Beaudet al. Human disorders of ubiquitination and proteasomal degradation. Curr Opin Pediatr. 2004; 16:419–26.

26. Foster MW, Hess DT, Stamler JS. Protein S-nitrosylation in health and disease: a current perspective. Trends Mol Med. 2009; 15:391–404.

27. Dudman NP, Guo XW, Gordon RB, Dawson PA, Wilcken DE. Human homocysteine catabolism: three major pathways and their relevance to development of arterial occlusive disease. J Nutr. 1996; 126:1295S–300S.

28. Brosnan JT, Jacobs RL, Stead LM, Brosnan ME. Methylation demand: a key determinant of homocysteine metabolism. Acta Biochim Pol. 2004; 51:405–13.

29. Ngoh GA, Facundo HT, Zafir A, Jones SP. O-GlcNAc signaling in the cardiovascular system. Circ Res. 2010; 107:171–85.

30. Mattson MP. Acetylation unleashes protein demons of dementia. Neuron. 2010; 67:900–2.

31. Lu Z, Scott I, Webster BR, Sack MN. The emerging characterization of lysine residue deacetylation on the modulation of mitochondrial function and cardiovascular biology. Circ Res. 2009; 105:830–41.

32. Méndez JD, Xie J, Aguilar-Hernández M, Méndez-Valenzuela V. Molecular susceptibility to glycation and its implication in diabetes mellitus and related diseases. Mol Cell Biochem. 2010; 344:185–93.

33. Jaeken J. Congenital disorders of glycosylation. Physician’s Guid to Treat Follow Metab Dis. Springer Berlin Heidelberg. 2006; p. 217–20.

34. Minguez P, Letunic I, Parca L, Garcia-Alonso L, Dopazo J, Huerta-Cepas J, et al. PTMcode v2: a resource for functional associations of post-translational modifications within and between proteins. Nucleic Acids Res. 2015; 43:D494–502.

35. Lu C-T, Huang K-Y, Su M-G, Lee T-Y, Bretaña NA, Chang W-C, et al. dbPTM 3.0: an informative resource for investigating substrate site specificity and functional association of protein post-translational modifications. Nucleic Acids Res. 2013; 41:D295–305.

36. Gnad F, Gunawardena J, Mann M. PHOSIDA 2011: the posttranslational modification database. Nucleic Acids Res. 2011;39:D253–60.

37. Sadowski I, Breitkreutz B-J, Stark C, Su T-C, Dahabieh M, Raithatha S, et al. The PhosphoGRID Saccharomyces cerevisiae protein phosphorylation site database: version 2.0 update. Database (Oxford). 2013; 2013:bat026.

38. Gupta R, Birch H, Rapacki K, Brunak S, Hansen JE. O-GLYCBASE version 4.0: a revised database of O-glycosylated proteins. Nucleic Acids Res. 1999; 27:370–2.

39. Bateman A. UniProt: A worldwide hub of protein knowledge. Nucleic Acids Res. 2019; 47(D1):D506–D515.

40. Sun M -a., Wang Y, Cheng H, Zhang Q, Ge W, Guo D. RedoxDB--a curated database for experimentally verified protein oxidative modification. Bioinformatics. 2012; 28:2551–2.

41. Chen Y-J, Lu C-T, Lee T-Y, Chen Y-J. dbGSH: a database of S-glutathionylation. Bioinformatics. 2014;30:2386–8.

42. Chen Y-J, Lu C-T, Su M-G, Huang K-Y, Ching W-C, Yang H-H, et al. dbSNO 2.0: a resource for exploring structural environment, functional and disease association and regulatory network of protein S-nitrosylation. Nucleic Acids Res. 2015; 43:D503–11.

43. Sayers EW, Barrett T, Benson DA, Bolton E, Bryant SH, Canese K, et al. Database resources of the National Center for Biotechnology Information. Nucleic Acids Res. 2011; 39:D38–51.

44. Koch L. Exploring human genomic diversity with gnomAD. Nat. Rev. Genet. 2020; 21(8):448.

45. Landrum MJ, Lee JM, Riley GR, Jang W, Rubinstein WS, Church DM, et al. ClinVar: public archive of relationships among sequence variation and human phenotype. Nucleic Acids Res. 2014; 42:D980–5.

46. Piñero J, Bravo A, Queralt-Rosinach N, Gutierrez-Sacristan A, Deu-Pons J, Centeno E, et al. DisGeNET: a comprehensive platform integrating information on human disease-associated genes and variants. Nucleic Acids Res. 2017;45:D833–9.

47. Ng PC, Henikoff S. Predicting deleterious amino acid substitutions. Genome Res. 2001; 11(5):863–74

48. Choi Y, Chan AP. PROVEAN web server: A tool to predict the functional effect of amino acid substitutions and indels. Bioinformatics. 2015; 31(16):2745–7.

49. Shihab HA, Gough J, Mort M, Cooper DN, Day INM, Gaunt TR. Ranking non-synonymous single nucleotide polymorphisms based on disease concepts. Hum Genomics. 2014;8(1):11.

50. Schwarz JM, Cooper DN, Schuelke M, Seelow D. Mutationtaster2: Mutation prediction for the deep-sequencing age. Nat. Methods. 2014; 11(4):361–2.

51. Amberger JS, Bocchini CA, Schiettecatte F, Scott AF, Hamosh A. OMIM.org: Online Mendelian Inheritance in Man (OMIM®), an online catalog of human genes and genetic disorders. Nucleic Acids Res. 2015;43:D789–98.

52. Pavan S, Rommel K, Mateo Marquina ME, Höhn S, Lanneau V, Rath A. Clinical Practice Guidelines for Rare Diseases: The Orphanet Database. PLoS One. 2017;12:e0170365.

53. Köhler S, Carmody L, Vasilevsky N, Jacobsen JOB, Danis D, Gourdine J-P, et al. Expansion of the Human Phenotype Ontology (HPO) knowledge base and resources. Nucleic Acids Res. 2019; 47:D1018–27.

54. Chitiprolu M, Jagow C, Tremblay V, Bondy-Chorney E, Paris G, Savard A, et al. A complex of C9ORF72 and p62 uses arginine methylation to eliminate stress granules by autophagy. Nat Commun. 2018; 9(1):2794.

55. Simandi Z, Pajer K, Karolyi K, Sieler T, Jiang LL, Kolostyak Z, et al. Arginine methyltransferase PRMT8 provides cellular stress tolerance in aging motoneurons. J Neurosci. 2018; 38:7683–700.

56. Xu B, Lu S, Gerton JL. Roberts syndrome. Rare Dis. 2014; 2:e27743.

57. Almedawar S, Colomina N, Bermúdez-López M, Pociño-Merino I, Torres-Rosell J. A SUMO-dependent step during establishment of sister chromatid cohesion. Curr Biol. 2012; 22:1576–81.

58. Era S, Abe T, Arakawa H, Kobayashi S, Szakal B, Yoshikawa Y, et al. The SUMO protease SENP1 is required for cohesion maintenance and mitotic arrest following spindle poison treatment. Biochem Biophys Res Commun. 2012; 426:310–6.

59. Meng Z, Jia LF, Gan YH. PTEN activation through K163 acetylation by inhibiting HDAC6 contributes to tumour inhibition. Oncogene. 2016; 35:2333–44.

60. Ikenoue T, Inoki K, Zhao B, Guan K-LL. PTEN acetylation modulates its interaction with PDZ domain. Cancer Res. 2008; 68:6908–12.

61. Savignac M, Edir A, Simon M, Hovnanian A. Darier disease: A disease model of impaired calcium homeostasis in the skin. Biochim. Biophys. Acta - Mol. Cell Res. 2011; p. 1111–7.

62. Adachi T, Weisbrod RM, Pimentel DR, Ying J, Sharov VS, Schöneich C, et al. S-glutathiolation by peroxynitrite activates SERCA during arterial relaxation by nitric oxide. Nat Med. 2004; 10:1200–7.

63. Lock JT, Sinkins WG, Schilling WP. Effect of protein S-glutathionylation on Ca2+ homeostasis in cultured aortic endothelial cells. Am J Physiol. 2011; 300:H493.

64. Dharssi S, Wong-Rieger D, Harold M, Terry S. Review of 11 national policies for rare diseases in the context of key patient needs. Orphanet J. Rare Dis. 2017;12(1):63.

65. Beltrao P, Bork P, Krogan NJ, van Noort V. Evolution and functional cross-talk of protein post-translational modifications. Mol Syst Biol. 2013; 9:714.

66. Naegle KM, Gymrek M, Joughin BA, Wagner JP, Welsch RE, Yaffe MB, et al. PTMScout, a Web resource for analysis of high throughput post-translational proteomics studies. Mol Cell Proteomics. 2010; 9:2558–70.

67. Kim Y, Kang C, Min B, Yi GS. Detection and analysis of disease-associated single nucleotide polymorphism influencing post-translational modification. BMC Med Genomics. 2015; 8:S7.

68. Piovesan D, Hatos A, Minervini G, Quaglia F, Monzon AM, Tosatto SCE. Assessing predictors for new post translational modification sites: A case study on hydroxylation. Iakoucheva LM. PLOS Comput Biol. 2020; 16:e1007967.

69. Pascovici D, Wu JX, McKay MJ, Joseph C, Noor Z, Kamath K, et al. Clinically relevant post-translational modification analyses—maturing workflows and bioinformatics tools. Int. J. Mol. Sci. 2019; 20(1):16.

70. Gonçalves E, Fragoulis A, Garcia-Alonso L, Cramer T, Saez-Rodriguez J, Beltrao P. Widespread Post-transcriptional Attenuation of Genomic Copy-Number Variation in Cancer. Cell Syst. 2017; 5:386-398.e4.

71. Woodsmith J, Kamburov A, Stelzl U. Dual coordination of post translational modifications in human protein networks. Russell RB, editor. PLoS Comput Biol. 2013; 9:e1002933.

72. Narita T, Weinert BT, Choudhary C. Functions and mechanisms of non-histone protein acetylation. Nat. Rev. Mol. Cell Biol. 2019; p.156–74.

73. Xue H, Peng J, Shang X. Predicting disease-related phenotypes using an integrated phenotype similarity measurement based on HPO. BMC Syst Biol. 2019; 13:34.

74. Beltrao P, Albanèse V, Kenner LR, Swaney DL, Burlingame A, Villén J, et al. Systematic Functional Prioritization of Protein Posttranslational Modifications. Cell. 2012; 150:413–25.

75. Holehouse AS, Naegle KM. Reproducible Analysis of Post-Translational Modifications in Proteomes—Application to Human Mutations. Lisacek F. PLoS One. 2015; 10:e0144692.

76. Roy S, Coldren C, Karunamurthy A, Kip NS, Klee EW, Lincoln SE, et al. Standards and Guidelines for Validating Next-Generation Sequencing Bioinformatics Pipelines: A Joint Recommendation of the Association for Molecular Pathology and the College of American Pathologists. J. Mol. Diagnostics. 2018; p. 4–27.

77. Sherry ST, Ward MH, Kholodov M, Baker J, Phan L, Smigielski EM, et al. dbSNP: the NCBI database of genetic variation. Nucleic Acids Res. 2001;29:308–11.

78. Collins R, Brand H, Karczewski K, Zhao X, Alföldi J, Francioli L, et al. An open resource of structural variation for medical and population genetics. bioRxiv. 2019;

79. Szklarczyk D, Franceschini A, Wyder S, Forslund K, Heller D, Huerta-Cepas J, et al. STRING v10: protein-protein interaction networks, integrated over the tree of life. Nucleic Acids Res. 2015; 43:D447–52.

80. Huerta-Cepas J, Szklarczyk D, Forslund K, Cook H, Heller D, Walter MC, et al. EGGNOG 4.5: A hierarchical orthology framework with improved functional annotations for eukaryotic, prokaryotic and viral sequences. Nucleic Acids Res. 2016; 44:D286–93.

81. Hamosh A, Scott AF, Amberger J, Bocchini C, Valle D, McKusick VA. Online Mendelian Inheritance in Man (OMIM), a knowledgebase of human genes and genetic disorders. Nucleic Acids Res.2002; 30:52–5.

82. Köhler S, Vasilevsky NA, Engelstad M, Foster E, McMurry J, Aymé S, et al. The human phenotype ontology in 2017. Nucleic Acids Res. 2017; 45:D865–76.

83. McLaren W, Gil L, Hunt SE, Riat HS, Ritchie GRS, Thormann A, et al. The Ensembl Variant Effect Predictor. Genome Biol. 2016; 17:122.

84. Huang K-Y, Su M-G, Kao H-J, Hsieh Y-C, Jhong J-H, Cheng K-H, et al. dbPTM 2016: 10-year anniversary of a resource for post-translational modification of proteins. Nucleic Acids Res. 2016; 44:D435–46.

85. Hornbeck P V., Kornhauser JM, Tkachev S, Zhang B, Skrzypek E, Murray B, et al. PhosphoSitePlus: a comprehensive resource for investigating the structure and function of experimentally determined post-translational modifications in man and mouse. Nucleic Acids Res. 2012; 40:D261–70.

86. Dinkel H, Chica C, Via A, Gould CM, Jensen LJ, Gibson TJ, et al. Phospho.ELM: a database of phosphorylation sites--update 2011. Nucleic Acids Res. 2011; 39:D261–7.

87. Stark C, Su TC, Breitkreutz A, Lourenco P, Dahabieh M, Breitkreutz BJ, et al. PhosphoGRID: a database of experimentally verified in vivo protein phosphorylation sites from the budding yeast Saccharomyces cerevisiae. Database (Oxford). 2010; 2010:bap026.

88. Keshava Prasad TS, Goel R, Kandasamy K, Keerthikumar S, Kumar S, Mathivanan S, et al. Human Protein Reference Database -2009 update. Nucleic Acids Res. 2009; 37(Database issue):D767–72

89. Huerta-Cepas J, Szklarczyk D, Forslund K, Cook H, Heller D, Walter MC, et al. EGGNOG 4.5: A hierarchical orthology framework with improved functional annotations for eukaryotic, prokaryotic and viral sequences. Nucleic Acids Res. 2016; 44:D286–93.

90. Su M-G, Weng JT-Y, Hsu JB-K, Huang K-Y, Chi Y-H, Lee T-Y. Investigation and identification of functional post-translational modification sites associated with drug binding and protein-protein interactions. BMC Syst Biol. 2017; 11:132.

91. Rust HL, Thompson PR. Kinase consensus sequences: A breeding ground for crosstalk. ACS Chem. Biol. 2011; 6(9):881–92.

92. Hamby SE, Hirst JD. Prediction of glycosylation sites using random forests. BMC Bioinformatics. 2008; 9:500.

93. Lee TY, Chen YJ, Lu TC, Huang H Da, Chen YJ. Snosite: Exploiting maximal dependence decomposition to identify cysteine S-Nitrosylation with substrate site specificity. PLoS One. 2011;6(7):e21849.

94. Marino SM, Gladyshev VN. Structural Analysis of Cysteine S-Nitrosylation: A Modified Acid-Based Motif and the Emerging Role of Trans-Nitrosylation. J Mol Biol. 2010; 395(4):844–59.

95. Lee TY, Chang CW, Lu CT, Cheng TH, Chang TH. Identification and characterization of lysine-methylated sites on histones and non-histone proteins. Comput Biol Chem. 2014; 50:11–8.

96. Lu CT, Lee TY, Chen YJ, Chen YJ. An Intelligent System for Identifying Acetylated Lysine on Histones and Nonhistone Proteins. Biomed Res Int. 2014; 2014:528650.

97. Aicart-Ramos C, Valero RA, Rodriguez-Crespo I. Protein palmitoylation and subcellular trafficking. Biochim. Biophys. Acta. 2011; 1808(12):2981–94.

98. Sigrist CJA, Cerutti L, Hulo N, Gattiker A, Falquet L, Pagni M, et al. PROSITE: a documented database using patterns and profiles as motif descriptors. Brief Bioinform. 2002; 3(3):265–74.

99. Necci M, Piovesan D, Dosztanyi Z, Tosatto SCE. MobiDB-lite: Fast and highly specific consensus prediction of intrinsic disorder in proteins. Bioinformatics. 2017; 33(9):1402–1404.

100. Bass JIF, Diallo A, Nelson J, Soto JM, Myers CL, Walhout AJM. Using networks to measure similarity between genes: Association index selection. Nat. Methods. 2013; 10(12):1169–76.

101. Shannon P, Markiel A, Ozier O, Baliga NS, Wang JT, Ramage D, et al. Cytoscape: a software environment for integrated models of biomolecular interaction networks. Genome Res. 2003; 13:2498–504.

