## Supplementary Tables for "A global map of the impact of deletion of Post-Translational Modification sites in genetic diseases"

**Table S1. Dataset of Post Translational Modifications (PTMs), nsSNVs and genetic diseases.** Total numbers of the datasets compiled for: protein post-translational modification sites in humans, human nsSNVs, genetic diseases from OMIM and Orphanet databases, phenotypes from the Human Phenotype Ontology database, and nsSNVs with disease and phenotype annotations as well as predicted as pathogenic.

| Collection | Total number |
|---|---|
| Post Translational Modification sites | 321,643 |
| nsSNVs coding variants | 4,464,412 |
| Diseases (ORPHANET and OMIM) | 12,388 |
| Phenotypes | 13,156 |
| nsSNVs coding variants associated to diseases | 44,969 |
| nsSNVs coding variants associated to phenotypes | 4,550 |
| Pathogenic nsSNVs coding variants | 392,185 |
| Diseases with proteins with PTMs and variants | 3,228 |
| Phenotypes with proteins with PTMs and variants | 1,909 |

**Table S2. Disease-related nsSNVs involved in disease families.** Orphanet disease families classification was adopted.

| Disease family | nsSNVs coding variants |
|---|---|
| Rare neurologic disease | 16,342 |
| Developmental defect during embryogenesis | 13,743 |
| Rare eye disease | 10,026 |
| Rare inborn errors of metabolism | 9,949 |
| Rare neoplastic disease | 7,729 |
| Rare skin disease | 6,831 |
| Rare renal disease | 6,422 |
| Rare bone disease | 5,524 |
| Rare cardiac disease | 5,093 |
| Rare endocrine disease | 4,553 |
| Rare gastroenterologic disease | 3,814 |

| | |
|---|---|
| Rare hematologic disease | 3,714 |
| Sucking/swallowing disorder | 3,358 |
| Rare gynecologic or obstetric disease | 2,376 |
| Rare infertility | 2,294 |
| Rare otorhinolaryngologic disease | 2,276 |
| Rare immune disease | 1,949 |
| Rare circulatory system disease | 1,829 |
| Rare respiratory disease | 1,817 |
| Rare systemic or rheumatologic disease | 1,674 |
| Rare hepatic disease | 1,612 |
| Rare surgical cardiac disease | 957 |
| Rare abdominal surgical disease | 957 |
| Rare urogenital disease | 742 |
| Rare odontologic disease | 734 |
| Rare maxillo-facial surgical disease | 718 |
| Rare surgical thoracic disease | 695 |
| Rare systemic or rheumatological disease of childhood | 435 |
| Rare infectious disease | 23 |
| Rare allergic disease | 21 |
| Unclassified disease | 932 |

**Table S3. Dataset of PTMs classified by type.** Number of human experimentally verified PTMs compiled, grouped by PTM types and sorted by abundance.

| PTM type | PTM Abbreviations | Number of sites | Number of proteins | Average PTM sites per protein |
|---|---|---|---|---|
| phosphorylation | Ph | 205,634 | 15,946 | 12.9 |
| ubiquitination | Ub | 55,279 | 9,391 | 5.9 |
| acetylation | Ac | 22,245 | 7,116 | 3.1 |
| methylation | Me | 14,370 | 4,786 | 2.7 |
| SUMOylation | SUMO | 13,488 | 3,816 | 3.5 |
| N-linked glycosylation | NG | 2,725 | 1,034 | 2.6 |
| O-GalNAc glycosylation | OGa | 1,922 | 428 | 4.5 |
| S-glutathionylation | Glu | 1,619 | 902 | 1.8 |
| S-nitrosylation | Ni | 1,184 | 684 | 1.7 |
| O-linked glycosylation | OG | 863 | 286 | 3.0 |
| proteolytic cleavage | PC | 808 | 344 | 2.3 |
| hydroxylation | Hy | 443 | 42 | 10.5 |
| O-GlcNAc glycosylation | OGl | 404 | 146 | 2.8 |
| caspase cleavage aspartic acid | CCAsp | 323 | 279 | 1.2 |
| citrullination | Ci | 178 | 64 | 2.8 |
| palmitoylation | Pal | 156 | 82 | 1.98 |
| intra-polypeptide disulfide bond formation | DisB | 131 | 60 | 2.2 |

| | | | | |
|---|---|---|---|---|
| ADP-ribosylation | ADP-r | 123 | 62 | 2 |
| sulfation | Sul | 90 | 34 | 2.62 |
| carboxylation | Ca | 85 | 12 | 7.1 |
| myristoylation | My | 78 | 76 | 1.1 |
| nitration | Nit | 73 | 54 | 1.3 |
| prenylation | Pre | 69 | 58 | 1.2 |
| pyrrolidone carboxylic acid formation | PyCar | 62 | 61 | 1.0 |
| glycation | Glu | 49 | 14 | 3.5 |
| protein-pyridoxal-5-phosphate linkage | P5Pl | 49 | 48 | 1.0 |
| amidation | Ami | 46 | 31 | 1.5 |
| oxidation | Oxi | 46 | 43 | 1.1 |
| malonylation | Ma | 45 | 33 | 1.4 |
| C-linked glycosylation | CG | 42 | 11 | 3.8 |
| neddylation | Nedd | 40 | 8 | 5.0 |
| glycophosphatidilinositol attachment | GPIa | 37 | 20 | 1.8 |
| isopeptide bond | IsoB | 25 | 17 | 1.5 |
| allysine | Ally | 20 | 17 | 1.2 |
| carbamidation | Cb | 20 | 17 | 1.2 |
| L-3-oxoalanine | L3O | 16 | 16 | 1.0 |
| binding to a nucleotide sequence | Nuc | 10 | 7 | 1.4 |
| glutamine deamidation | GluDea | 10 | 7 | 1.4 |
| topaquinone formation | Topa | 8 | 8 | 1.0 |
| iodination | Io | 7 | 1 | 7.0 |
| N6-retinylidene-L-lysine | N6RL | 7 | 7 | 1.0 |
| alkylation | Alky | 6 | 3 | 2.0 |
| transglutamination | Tglu | 6 | 3 | 2.0 |
| O-phosphopantetheine-L-serine | OPLS | 5 | 5 | 1.0 |
| L-glutamyl 5-glycerylphosphorylethanolamine | LG5GP | 4 | 2 | 2.0 |
| protein-FAD linkage | FADl | 4 | 4 | 1.0 |
| farnesylation | Far | 3 | 3 | 1.0 |
| glycosylation | Glyco | 3 | 3 | 1.0 |
| unidentified N-terminal blocking modifications | UntBM | 3 | 3 | 1.0 |
| aspartic acid conversion into alanine | Asp2Ala | 1 | 1 | 1.0 |
| dehydroalanine | DHAla | 1 | 1 | 1.0 |
| dipyrrolylmethanemethyl-L-cysteine | DipMetLCys | 1 | 1 | 1.0 |
| glycoprotein | GlyP | 1 | 1 | 1.0 |
| N6-1-carboxyethyl-L-lysine | N61CLys | 1 | 1 | 1.0 |
| N6-carboxylysine | N6CaL | 1 | 1 | 1.0 |
| N-pyruvic acid 2-iminyl-L-cysteine | Np2ICys | 1 | 1 | 1.0 |
| serine conversion into pyruvic acid | Ser2Pyr | 1 | 1 | 1.0 |
| S-linked glycosylation | SG | 1 | 1 | 1.0 |
| thioether bond | TB | 1 | 1 | 1.0 |

**Table S4. Mapping of nsSNVs and PTM sites.** Human PTMs and nsSNVs co-occurrences. A co-occurrence is considered when the nsSNV affects the modifiable residue or is located within a ±5 amino acids from the PTM site.

| PTM types | nsSNVs disrupting PTM sites | Affected PTM sites | Proteins involved | Co-ocurrences between nsSNVs and PTM sites |
|---|---|---|---|---|
| phosphorylation | 792,384 | 198,970 | 15,728 | 1,167,626 |
| ubiquitination | 240,961 | 52,925 | 9,257 | 269,550 |
| acetylation | 94,839 | 21,369 | 6,929 | 112,983 |
| methylation | 63,282 | 12,579 | 4,691 | 73,637 |
| SUMOylation | 62,176 | 12,911 | 3,742 | 66,663 |
| N-linked glycosylation | 15,671 | 2,682 | 1,023 | 16,310 |
| glutathionylation | 7,465 | 1,540 | 876 | 7,553 |
| S-nitrosylation | 5,567 | 1,119 | 653 | 5,822 |
| O-GalNAc glycosylation | 7,636 | 1,906 | 424 | 12,914 |
| O-linked glycosylation | 4,085 | 844 | 282 | 5,410 |
| O-GlcNAc glycosylation | 1,677 | 384 | 139 | 2,281 |
| proteolytic cleavage | 586 | 401 | 213 | 586 |
| hydroxylation | 328 | 242 | 28 | 328 |
| malonylation | 209 | 44 | 33 | 213 |
| carboxylation | 54 | 41 | 10 | 54 |
| citrullination | 195 | 105 | 47 | 195 |
| neddylation | 187 | 40 | 8 | 267 |
| caspase cleavage aspartic acid | 179 | 141 | 133 | 179 |
| palmitoylation | 83 | 61 | 41 | 83 |
| intra-polypeptide disulfide bond formation | 71 | 61 | 41 | 71 |
| ADP-ribosylation | 65 | 52 | 36 | 72 |
| sulfation | 59 | 35 | 21 | 59 |
| glycation | 35 | 14 | 8 | 35 |
| pyrrolidone carboxylic acid formation | 33 | 24 | 24 | 33 |
| nitration | 31 | 24 | 22 | 31 |
| oxidation | 29 | 22 | 22 | 29 |
| myristoylation | 27 | 22 | 22 | 27 |
| amidation | 24 | 21 | 16 | 24 |
| isopeptide bond | 19 | 13 | 8 | 19 |
| prenylation | 18 | 15 | 14 | 18 |
| allysine | 17 | 9 | 8 | 17 |
| protein-pyridoxal-5-phosphate linkage | 16 | 12 | 12 | 16 |
| carbamidation | 13 | 8 | 8 | 13 |
| C-linked glycosylation | 12 | 10 | 7 | 12 |
| binding to a nucleotide sequence | 12 | 5 | 3 | 12 |
| glycophosphatidilinositol | 10 | 8 | 7 | 10 |

| | | | | |
|---|---|---|---|---|
| attachment | | | | |
| glutamine deamidation | 10 | 5 | 5 | 10 |
| L-3-oxoalanine | 8 | 5 | 5 | 8 |
| protein-FAD linkage | 6 | 3 | 3 | 6 |
| N6-retinylidene-L-lysine | 4 | 3 | 3 | 4 |
| alkylation | 4 | 3 | 2 | 4 |
| iodination | 4 | 2 | 1 | 4 |
| transglutamination | 4 | 3 | 1 | 4 |
| unidentified N-terminal blocking modifications | 3 | 2 | 2 | 3 |
| serine conversion into pyruvic acid | 3 | 1 | 1 | 3 |
| glycosylation | 2 | 2 | 2 | 2 |
| topaquinone formation | 1 | 1 | 1 | 1 |
| O-phosphopantetheine-L-serine | 1 | 1 | 1 | 1 |
| dehydroalanine | 1 | 1 | 1 | 1 |
| N6-carboxylysine | 1 | 1 | 1 | 1 |
| dipyrrolylmethanemethyl-L-cysteine | 1 | 1 | 1 | 1 |
| thioether bond | 1 | 1 | 1 | 1 |
| N-pyruvic acid 2-iminyl-L-cysteine | 1 | 1 | 1 | 1 |
| L-glutamyl 5-glycerylphosphorylethanolamine | 0 | 0 | 0 | 0 |
| farnesylation | 0 | 0 | 0 | 0 |
| aspartic acid conversion into alanine | 0 | 0 | 0 | 0 |
| glycoprotein | 0 | 0 | 0 | 0 |
| N6-1-carboxyethyl-L-lysine | 0 | 0 | 0 | 0 |
| S-linked glycosylation | 0 | 0 | 0 | 0 |
| **ALL PTM TYPES** | **1,093,134** | **308,695** | **16,541** | **1,743,207** |

**Table S5. Crossmatching of PTM sites and disease-related nsSNVs**. Number of disease-related nsSNVs and PTM sites matched grouped by PTM type. We include number of proteins and diseases affected. PTMs with no matches are not shown.

| PTM type | Disease-related nsSNVs disrupting PTMs | Affected PTMs | Co-occurrences of disease-related nsSNVs and PTMs | Proteins | Diseases |
|---|---|---|---|---|---|
| phosphorylation | 6,685 | 6,056 | 9,715 | 1,438 | 1,509 |
| ubiquitination | 2,227 | 1,611 | 2,434 | 703 | 739 |
| acetylation | 1,175 | 858 | 1,379 | 410 | 499 |
| methylation | 630 | 422 | 691 | 270 | 320 |
| SUMOylation | 546 | 359 | 583 | 190 | 232 |

| | | | | | |
|---|---|---|---|---|---|
| N-linked glycosylation | 279 | 178 | 290 | 111 | 150 |
| S-glutathionylation | 137 | 91 | 141 | 68 | 89 |
| S-nitrosylation | 96 | 61 | 101 | 44 | 66 |
| O-linked glycosylation | 65 | 55 | 78 | 32 | 52 |
| O-GalNAc glycosylation | 53 | 56 | 73 | 24 | 29 |
| proteolytic cleavage | 45 | 27 | 45 | 17 | 24 |
| O-GlcNAc glycosylation | 31 | 19 | 33 | 13 | 25 |
| carboxylation | 12 | 9 | 12 | 6 | 6 |
| malonylation | 6 | 1 | 6 | 1 | 1 |
| neddylation | 4 | 7 | 7 | 2 | 4 |
| ADP-ribosylation | 4 | 3 | 4 | 3 | 7 |
| hydroxylation | 3 | 2 | 3 | 2 | 3 |
| palmitoylation | 2 | 2 | 2 | 2 | 2 |
| intra-polypeptide disulfide bond formation | 2 | 2 | 2 | 2 | 2 |
| sulfation | 2 | 2 | 2 | 1 | 1 |
| citrullination | 1 | 1 | 1 | 1 | 1 |
| glycation | 1 | 1 | 1 | 1 | 1 |
| protein-pyridoxal-5-phosphate linkage | 1 | 1 | 1 | 1 | 1 |
| N6-retinylidene-L-lysine | 1 | 1 | 1 | 1 | 1 |
| protein-FAD linkage | 1 | 1 | 1 | 1 | 1 |
| alkylation | 1 | 1 | 1 | 1 | 1 |
| **ALL PTM TYPES** | **9,684** | **9,827** | **15,607** | **1,723** | **1,790** |

**Table S6. Crossmatching of PTM sites and predicted pathogenic nsSNVs**. Number of predicted pathogenic nsSNVs and PTM sites matched grouped by PTM type. We include number of proteins and diseases affected. PTMs with no matches are not shown.

| PTM types | Pathogenic nsSNVs disrupting PTMs | Affected PTMs | Co-occurrences of pathogenic nsSNVs and PTMs | Proteins |
|---|---|---|---|---|
| phosphorylation | 60,060 | 62,530 | 88,137 | 11,203 |
| ubiquitination | 19,263 | 15,702 | 21,501 | 5,628 |
| acetylation | 7,567 | 6,298 | 8,902 | 3,176 |
| methylation | 4,964 | 4,013 | 5,783 | 2,096 |
| SUMOylation | 4,621 | 3,629 | 4,948 | 1,837 |
| N-linked glycosylation | 1,279 | 947 | 1,335 | 545 |
| S-glutathionylation | 697 | 475 | 702 | 360 |
| S-nitrosylation | 538 | 371 | 555 | 274 |
| O-linked glycosylation | 351 | 300 | 449 | 139 |
| O-GalNAc glycosylation | 334 | 373 | 470 | 171 |
| O-GlcNAc glycosylation | 156 | 139 | 212 | 69 |

| | | | | |
|---|---|---|---|---|
| proteolytic cleavage | 86 | 68 | 86 | 49 |
| hydroxylation | 43 | 39 | 43 | 15 |
| neddylation | 36 | 24 | 51 | 6 |
| malonylation | 27 | 18 | 27 | 17 |
| carboxylation | 15 | 14 | 15 | 6 |
| ADP-ribosylation | 13 | 8 | 13 | 7 |
| palmitoylation | 9 | 8 | 9 | 8 |
| myristoylation | 6 | 5 | 6 | 5 |
| caspase cleavage aspartic acid | 6 | 6 | 6 | 6 |
| intra-polypeptide disulfide bond formation | 6 | 6 | 6 | 5 |
| citrullination | 7 | 7 | 7 | 6 |
| sulfation | 5 | 5 | 5 | 3 |
| oxidation | 4 | 4 | 4 | 4 |
| amidation | 4 | 4 | 4 | 4 |
| protein-pyridoxal-5-phosphate linkage | 4 | 4 | 4 | 4 |
| glutamine deamidation | 4 | 2 | 4 | 2 |
| binding to a nucleotide sequence | 2 | 2 | 2 | 1 |
| glycation | 1 | 1 | 1 | 1 |
| C-linked glycosylation | 1 | 1 | 1 | 1 |
| prenylation | 1 | 1 | 1 | 1 |
| protein-FAD linkage | 1 | 1 | 1 | 1 |
| N6-retinylidene-L-lysine | 1 | 1 | 1 | 1 |
| glycosylation | 1 | 1 | 1 | 1 |
| **ALL PTM TYPES** | **84,387** | **95,007** | **133,292** | **16,528** |

**Table S7. Data for the enrichment of disease-related nsSNVs in PTM types**. Data and results for Figure 1D.

| PTM type | Protein region | Disease-nsSNVs matching PTMs | Disease-nsSNVs not matching PTMs | Non-disease-nsSNVs matching PTMs | Non-disease-nsSNVs not matching PTMs | p-value | FDR |
|---|---|---|---|---|---|---|---|
| Ph | Disordered | 1,260 | 2,512 | 237,465 | 48,366 | 2.73e-01 | 5.92e-01 |
| Ac | Disordered | 104 | 1,546 | 19,488 | 235,799 | 9.83e-01 | 1.00e+00 |
| Ub | Disordered | 99 | 1,529 | 19,617 | 253,226 | 9.65e-01 | 1.00e+00 |
| Me | Disordered | 95 | 1,127 | 20,558 | 193,650 | 9.88e-01 | 1.00e+00 |
| SUMO | Disordered | 44 | 570 | 10,648 | 114,035 | 9.04e-01 | 1.00e+00 |
| OG | Disordered | 17 | 203 | 1,282 | 21,062 | 1.34e-01 | 5.31e-01 |
| OGa | Disordered | 14 | 305 | 2,880 | 21,665 | 1.00e+00 | 1.00e+00 |

| | | | | | | | |
|---|---|---|---|---|---|---|---|
| OGl | Disordered | 11 | 35 | 480 | 9,563 | 9.17e-06 | 1.19e-04 |
| NG | Disordered | 7 | 222 | 406 | 19,129 | 2.04e-01 | 5.31e-01 |
| PC | Disordered | 5 | 239 | 101 | 10,665 | 8.64e-02 | 5.31e-01 |
| Glu | Disordered | 2 | 48 | 227 | 13,764 | 1.96e-01 | 5.31e-01 |
| Hy | Disordered | 1 | 399 | 307 | 6,025 | 1.00e+00 | 1.00e+00 |
| Nedd | Disordered | 1 | 1 | 49 | 79 | 6.23e-01 | 1.00e+00 |
| Ph | Ordered | 5,425 | 32,654 | 548,234 | 3,520,500 | 6.71e-06 | 3.49e-05 |
| Ub | Ordered | 2,128 | 23,793 | 219,117 | 2,381,860 | 8.95e-01 | 9.46e-01 |
| Ac | Ordered | 1,071 | 19,752 | 74,176 | 2,021,551 | 8.30e-32 | 2.16e-30 |
| Me | Ordered | 535 | 14,280 | 42,094 | 1,412,366 | 3.28e-07 | 2.84e-06 |
| SUMO | Ordered | 502 | 8,959 | 50,982 | 983,403 | 4.93e-02 | 1.43e-01 |
| NG | Ordered | 272 | 6,443 | 14986 | 345,553 | 6.75e-01 | 7.98e-01 |
| Glu | Ordered | 135 | 2,618 | 7,101 | 210,771 | 3.99e-06 | 2.60e-05 |
| Ni | Ordered | 96 | 2,640 | 5,365 | 158,453 | 2.62e-01 | 5.24e-01 |
| OG | Ordered | 48 | 1,613 | 2,738 | 99,698 | 3.14e-01 | 5.44e-01 |
| PC | Ordered | 40 | 2,799 | 440 | 107,492 | 1.21e-10 | 1.57e-09 |
| OGa | Ordered | 39 | 1,651 | 4,703 | 158,383 | 9.35e-01 | 9.46e-01 |
| OGl | Ordered | 20 | 940 | 1,166 | 57,422 | 4.51e-01 | 6.89e-01 |
| Ca | Ordered | 12 | 329 | 42 | 3,638 | 1.49e-03 | 6.44e-03 |
| Ma | Ordered | 6 | 63 | 203 | 5,438 | 3.96e-02 | 1.29e-01 |
| ADP-r | Ordered | 4 | 145 | 37 | 7,983 | 6.38e-03 | 2.37e-02 |
| Nedd | Ordered | 3 | 85 | 134 | 1,804 | 9.46e-01 | 9.46e-01 |
| Hy | Ordered | 2 | 531 | 18 | 14,405 | 1.58e-01 | 3.59e-01 |
| Pa | Ordered | 2 | 590 | 76 | 21,328 | 6.25e-01 | 7.98e-01 |
| DisB | Ordered | 2 | 267 | 69 | 10,532 | 5.28e-01 | 7.22e-01 |
| Sul | Ordered | 2 | 477 | 53 | 11,150 | 6.66e-01 | 7.98e-01 |
| Ci | Ordered | 1 | 104 | 26 | 8,300 | 2.87e-01 | 5.34e-01 |
| Gly | Ordered | 1 | 160 | 34 | 2,418 | 8.94e-01 | 9.46e-01 |
| P5Pl | Ordered | 1 | 416 | 15 | 14,015 | 3.74e-01 | 6.08e-01 |
| FADl | Ordered | 1 | 134 | 5 | 1,083 | 5.05e-01 | 7.22e-01 |
| N6RL | Ordered | 1 | 51 | 3 | 1,121 | 1.66e-01 | 3.59e-01 |
| Alky | Ordered | 1 | 42 | 3 | 2,038 | 8.01e-02 | 2.08e-01 |

**Table S8. Data for the enrichment of predicted pathogenic nsSNVs in PTM types**. Data and results for Figure 1D.

| **PTM type** | **Protein region** | **Pathogenic nsSNVs matching PTMs** | **Pathogenic nsSNVs not matching PTMs** | **Non-pathogenic nsSNVs matching PTMs** | **Non-pathogenic nsSNVs not matching PTMs** | **p-value** | **FDR** |
|---|---|---|---|---|---|---|---|
| Ph | Disordered | 13,946 | 24,590 | 224,779 | 461,588 | 4.48e-44 | 8.45e-43 |
| Me | Disordered | 1,289 | 10,079 | 19,364 | 184,698 | 1.06e-10 | 1.06e-09 |

| | | | | | | | |
|---|---|---|---|---|---|---|---|
| Ac | Disordered | 1,180 | 13,751 | 18,412 | 223,594 | 9.69e-02 | 2.77e-01 |
| Ub | Disordered | 1,074 | 12,640 | 18,642 | 242,115 | 1.51e-03 | 7.56e-03 |
| SUMO | Disordered | 527 | 5,442 | 10,165 | 109,163 | 2.07e-01 | 3.77e-01 |
| OG | Disordered | 122 | 1,344 | 1,177 | 19,921 | 2.22e-05 | 1.48e-04 |
| OGa | Disordered | 97 | 1,435 | 2,797 | 20,535 | 1.00e+00 | 1.00e+00 |
| OGl | Disordered | 38 | 617 | 453 | 8,981 | 1.46e-01 | 3.61e-01 |
| Hy | Disordered | 38 | 1,033 | 270 | 5,391 | 9.70e-01 | 1.00e+00 |
| NG | Disordered | 16 | 1,564 | 397 | 17,787 | 1.00e+00 | 1.00e+00 |
| PC | Disordered | 12 | 1,056 | 94 | 9,848 | 3.31e-01 | 5.13e-01 |
| Glu | Disordered | 10 | 715 | 219 | 13,097 | 7.51e-01 | 1.00e+00 |
| Nedd | Disordered | 7 | 15 | 43 | 65 | 8.27e-01 | 1.00e+00 |
| Ni | Disordered | 4 | 400 | 102 | 8,480 | 7.08e-01 | 1.00e+00 |
| Ci | Disordered | 4 | 45 | 163 | 1,002 | 9.24e-01 | 1.00e+00 |
| CCAsp | Disordered | 3 | 1,021 | 86 | 16,037 | 9.07e-01 | 1.00e+00 |
| My | Disordered | 3 | 66 | 8 | 1,621 | 8.38e-03 | 3.53e-02 |
| Pa | Disordered | 1 | 22 | 4 | 531 | 1.90e-01 | 3.77e-01 |
| Ami | Disordered | 1 | 20 | 2 | 638 | 9.24e-02 | 2.77e-01 |
| Pre | Disordered | 1 | 30 | 1 | 334 | 1.62e-01 | 3.61e-01 |
| Ph | Ordered | 46,114 | 287,066 | 507,545 | 3,266,088 | 1.41e-10 | 4.67e-09 |
| Ub | Ordered | 18,189 | 188,998 | 203,056 | 2,216,655 | 7.07e-10 | 1.17e-08 |
| Ac | Ordered | 6,387 | 168,008 | 68,860 | 1,873,295 | 6.07e-03 | 2.50e-02 |
| SUMO | Ordered | 4,094 | 77,502 | 47,390 | 914,860 | 1.23e-01 | 2.79e-01 |
| Me | Ordered | 3,675 | 116,073 | 38,954 | 1,310,573 | 1.79e-04 | 1.18e-03 |
| NG | Ordered | 1,263 | 32,830 | 13,995 | 319,166 | 1.00e+00 | 1.00e+00 |
| Glu | Ordered | 687 | 17,842 | 6,549 | 195,547 | 4.19e-04 | 2.30e-03 |
| Ni | Ordered | 534 | 14,341 | 4,927 | 146,852 | 9.47e-03 | 3.47e-02 |
| OGa | Ordered | 237 | 13,018 | 4,505 | 147,016 | 1.00e+00 | 1.00e+00 |
| OG | Ordered | 229 | 9,276 | 2,557 | 92,035 | 9.59e-01 | 1.00e+00 |
| OGl | Ordered | 118 | 5,445 | 1,068 | 52,917 | 2.47e-01 | 4.41e-01 |
| Nedd | Ordered | 29 | 374 | 108 | 1,515 | 3.84e-01 | 5.51e-01 |
| Ma | Ordered | 27 | 553 | 182 | 4,948 | 1.12e-01 | 2.79e-01 |
| Ca | Ordered | 15 | 447 | 39 | 3,520 | 7.83e-04 | 3.69e-03 |
| ADP-r | Ordered | 13 | 755 | 28 | 7,373 | 5.90e-05 | 4.87e-04 |
| Pa | Ordered | 8 | 2,325 | 70 | 19,593 | 5.94e-01 | 7.85e-01 |
| DisB | Ordered | 6 | 1,232 | 65 | 9,567 | 8.34e-01 | 9.83e-01 |
| Hy | Ordered | 5 | 1,568 | 15 | 13,368 | 5.18e-02 | 1.42e-01 |
| Sul | Ordered | 5 | 1,174 | 50 | 10,453 | 6.64e-01 | 8.42e-01 |
| Oxi | Ordered | 4 | 938 | 25 | 8,836 | 3.02e-01 | 4.75e-01 |
| P5Pl | Ordered | 4 | 1,165 | 12 | 13,266 | 3.53e-02 | 1.06e-01 |
| GluDea | Ordered | 4 | 108 | 6 | 798 | 2.48e-02 | 8.17e-02 |
| Ci | Ordered | 3 | 581 | 24 | 7,283 | 2.87e-01 | 4.73e-01 |
| CCAsp | Ordered | 3 | 8,166 | 87 | 89,085 | 9.84e-01 | 1.00e+00 |

| | | | | | | | |
|---|---|---|---|---|---|---|---|
| My | Ordered | 3 | 1,369 | 13 | 16,029 | 1.27e-01 | 2.79e-01 |
| Ami | Ordered | 3 | 132 | 18 | 1,915 | 1.53e-01 | 3.16e-01 |
| Nuc | Ordered | 2 | 60 | 10 | 692 | 2.54e-01 | 4.41e-01 |
| Gly | Ordered | 1 | 281 | 34 | 2,297 | 9.82e-01 | 1.00e+00 |
| CG | Ordered | 1 | 419 | 11 | 3,306 | 7.61e-01 | 9.31e-01 |
| FADI | Ordered | 1 | 101 | 5 | 1,116 | 4.08e-01 | 5.60e-01 |
| N6RL | Ordered | 1 | 106 | 3 | 1,066 | 3.18e-01 | 4.76e-01 |
| Glyco | Ordered | 1 | 135 | 1 | 1,063 | 2.14e-01 | 4.15e-01 |

**Table S9. PTM types and genetic diseases predicted associations.** Together with p-values and FDR, Occupancy scores (OS) are also shown. Next to last columns holds the category related to the previous knowledge of the association, three classes: known, previous evidences, divided into one variant case study (OVCS), evidences in related diseases (ERD) and regulatory evidences (RE), and novel. Last column shows the PubMed Ids of the scientific papers supporting the classification.

| PTM type | Disease ID | nsSNVs involved in association | Total nsSNVs associated to disease | p-value | FDR | OS | OS degree | Previous knowledge category | PMID |
|---|---|---|---|---|---|---|---|---|---|
| Ac | ORPHA:289869 | 35 | 138 | 2.30e-24 | 7.34e-21 | 100 | VERY HIGH | OVCS | 19318352 |
| Ac | ORPHA:2869 | 35 | 142 | 6.54e-24 | 1.04e-20 | 100 | VERY HIGH | novel | - |
| Ac | ORPHA:3208 | 33 | 130 | 4.58e-23 | 4.87e-20 | 100 | VERY HIGH | novel | - |
| Ac | ORPHA:217604 | 39 | 287 | 8.90e-17 | 7.10e-14 | 100 | VERY HIGH | RE | 28542596 26101264 |
| Ac | ORPHA:29072 | 45 | 403 | 7.54e-16 | 4.81e-13 | 100 | VERY HIGH | novel | - |
| Ac | ORPHA:35878 | 8 | 10 | 1.06e-11 | 5.63e-09 | 80 | HIGH | RE | 20934340 30683850 |
| Ac | OMIM:192600 | 46 | 568 | 3.69e-11 | 1.53e-08 | 100 | VERY HIGH | RE | 28542596 26101264 |
| Ac | ORPHA:171680 | 12 | 34 | 3.84e-11 | 1.53e-08 | 100 | VERY HIGH | novel | - |
| Ac | ORPHA:85173 | 7 | 8 | 7.29e-11 | 2.58e-08 | 0 | VERY LOW | novel | - |
| Ac | ORPHA:102011 | 12 | 36 | 8.34e-11 | 2.66e-08 | 100 | VERY HIGH | novel | - |
| Ac | ORPHA:2309 | 13 | 47 | 1.93e-10 | 5.58e-08 | 28 | LOW | novel | - |
| Ac | ORPHA:740 | 9 | 18 | 2.56e-10 | 6.80e-08 | 92 | VERY HIGH | RE | 23217256 |
| Ac | ORPHA:746 | 11 | 33 | 5.15e-10 | 1.26e-07 | 100 | VERY HIGH | known | 27457618 |
| Ac | ORPHA:713 | 7 | 12 | 6.57e-09 | 1.50e-06 | 100 | VERY HIGH | RE | 22225877 |
| Ac | ORPHA:664 | 9 | 25 | 9.09e-09 | 1.93e-06 | 85 | VERY HIGH | OVCS | 19318352 |
| Ac | ORPHA:783 | 13 | 66 | 1.75e-08 | 3.49e-06 | 97 | VERY HIGH | novel | - |
| Ac | ORPHA:134 | 8 | 24 | 1.24e-07 | 2.20e-05 | 86 | VERY HIGH | novel | - |
| Ac | ORPHA:300751 | 8 | 24 | 1.24e-07 | 2.20e-05 | 100 | VERY HIGH | RE | 22645641 26101264 30131726 |

| | | | | | | | | | |
|---|---|---|---|---|---|---|---|---|---|
| Ac | ORPHA:98855 | 24 | 261 | 1.73e-07 | 2.90e-05 | 97 | VERY HIGH | novel | - |
| Ac | ORPHA:3322 | 15 | 116 | 4.92e-07 | 7.85e-05 | 25 | LOW | novel | - |
| Ac | ORPHA:353277 | 11 | 63 | 7.90e-07 | 1.20e-04 | 81 | VERY HIGH | novel | - |
| Ac | ORPHA:414 | 9 | 41 | 1.06e-06 | 1.54e-04 | 74 | HIGH | novel | - |
| Ac | ORPHA:284435 | 5 | 9 | 1.53e-06 | 2.12e-04 | 78 | HIGH | RE | 22225877 |
| Ac | ORPHA:1478 | 11 | 68 | 1.74e-06 | 2.32e-04 | 22 | LOW | ERD | 29769443 |
| Ac | OMIM:616973 | 4 | 5 | 2.45e-06 | 3.12e-04 | 21 | LOW | novel | - |
| Ac | ORPHA:157973 | 5 | 10 | 2.99e-06 | 3.67e-04 | 73 | HIGH | novel | - |
| Ac | ORPHA:699 | 10 | 59 | 3.32e-06 | 3.92e-04 | 92 | VERY HIGH | novel | - |
| Ac | ORPHA:98473 | 9 | 47 | 3.58e-06 | 4.08e-04 | 57 | MEDIUM | RE | 26302492 |
| Ac | ORPHA:868 | 4 | 6 | 7.19e-06 | 7.91e-04 | 68 | HIGH | RE | 21879450<br>19410549 |
| Ac | ORPHA:2995 | 7 | 30 | 1.11e-05 | 1.18e-03 | 69 | HIGH | novel | - |
| Ac | OMIM:601728 | 4 | 7 | 1.64e-05 | 1.69e-03 | 42 | MEDIUM | novel | - |
| Ac | ORPHA:93616 | 5 | 14 | 2.17e-05 | 2.17e-03 | 98 | VERY HIGH | ERD | 4531009<br>10974199 |
| Ac | ORPHA:109009 | 11 | 88 | 2.24e-05 | 2.17e-03 | 70 | HIGH | RE | 25911675 |
| Ac | ORPHA:407 | 11 | 93 | 3.79e-05 | 3.55e-03 | 33 | LOW | novel | - |
| Ac | ORPHA:469 | 5 | 17 | 6.29e-05 | 5.71e-03 | 40 | LOW | novel | - |
| Ac | ORPHA:79396 | 6 | 27 | 6.45e-05 | 5.71e-03 | 14 | VERY LOW | novel | - |
| Ac | ORPHA:232 | 3 | 4 | 7.38e-05 | 5.92e-03 | 61 | HIGH | known | 4531009 |
| Ac | ORPHA:280615 | 3 | 4 | 7.38e-05 | 5.92e-03 | 47 | MEDIUM | known | 4531009 |
| Ac | ORPHA:438114 | 3 | 4 | 7.38e-05 | 5.92e-03 | 0 | VERY LOW | novel | - |
| Ac | ORPHA:217569 | 25 | 396 | 7.42e-05 | 5.92e-03 | 33 | LOW | RE | 28542596<br>26101264 |
| Ac | ORPHA:137898 | 5 | 19 | 1.13e-04 | 8.59e-03 | 24 | LOW | novel | - |
| Ac | ORPHA:603 | 5 | 19 | 1.13e-04 | 8.59e-03 | 10 | VERY LOW | known | 30611609 |
| Ac | ORPHA:101028 | 4 | 11 | 1.42e-04 | 1.03e-02 | 44 | MEDIUM | novel | - |
| Ac | ORPHA:832 | 4 | 11 | 1.42e-04 | 1.03e-02 | 22 | LOW | RE | 24725594 |
| Ac | ORPHA:124 | 6 | 31 | 1.46e-04 | 1.04e-02 | 92 | VERY HIGH | novel | - |
| Ac | OMIM:242880 | 3 | 5 | 1.81e-04 | 1.20e-02 | 0 | VERY LOW | novel | - |
| Ac | ORPHA:264 | 3 | 5 | 1.81e-04 | 1.20e-02 | 69 | HIGH | ERD | 26302492 |
| Ac | ORPHA:68364 | 3 | 5 | 1.81e-04 | 1.20e-02 | 89 | VERY HIGH | known | 4531009 |
| Ac | ORPHA:26793 | 7 | 46 | 2.01e-04 | 1.31e-02 | 68 | HIGH | RE | 20934340 |
| Ac | ORPHA:93599 | 4 | 12 | 2.09e-04 | 1.33e-02 | 1 | VERY LOW | novel | - |
| Ac | ORPHA:91387 | 17 | 236 | 2.21e-04 | 1.39e-02 | 2 | VERY LOW | novel | - |
| Ac | OMIM:613251 | 8 | 62 | 2.33e-04 | 1.43e-02 | 64 | HIGH | RE | 28542596,<br>26101264 |
| Ac | ORPHA:65285 | 6 | 34 | 2.50e-04 | 1.50e-02 | 21 | LOW | novel | - |
| Ac | ORPHA:58 | 9 | 80 | 2.78e-04 | 1.64e-02 | 89 | VERY HIGH | novel | - |
| Ac | ORPHA:19 | 3 | 6 | 3.55e-04 | 2.06e-02 | 38 | LOW | novel | - |
| Ac | ORPHA:477 | 4 | 14 | 4.04e-04 | 2.30e-02 | 71 | HIGH | novel | - |
| Ac | ORPHA:98306 | 6 | 39 | 5.41e-04 | 3.01e-02 | 66 | HIGH | novel | - |
| Ac | ORPHA:79399 | 5 | 26 | 5.48e-04 | 3.01e-02 | 2 | VERY LOW | novel | - |
| Ac | ORPHA:210548 | 3 | 7 | 6.08e-04 | 3.29e-02 | 18 | VERY LOW | novel | - |
| Ac | ORPHA:187 | 6 | 40 | 6.23e-04 | 3.31e-02 | 29 | LOW | known | 14633929 |
| Ac | ORPHA:98909 | 5 | 27 | 6.58e-04 | 3.44e-02 | 19 | VERY LOW | novel | - |
| ADP-r | ORPHA:93276 | 2 | 5 | 6.16e-08 | 1.96e-04 | 15 | VERY LOW | novel | - |
| ADP-r | ORPHA:189427 | 2 | 7 | 1.29e-07 | 2.06e-04 | 7 | VERY LOW | novel | - |
| ADP-r | ORPHA:96256 | 2 | 17 | 8.38e-07 | 8.91e-04 | 7 | VERY LOW | novel | - |

| | | | | | | | | | |
|---|---|---|---|---|---|---|---|---|---|
| Ca | ORPHA:328 | 4 | 28 | 6.39e-11 | 2.04e-07 | 72 | HIGH | known | 19141158 |
| Ca | ORPHA:98879 | 3 | 63 | 6.05e-07 | 9.64e-04 | 73 | HIGH | RE | 19141158<br>22872812 |
| Glu | ORPHA:803 | 12 | 149 | 5.42e-14 | 1.73e-10 | 100 | VERY HIGH | OVCS | 24253732 |
| Glu | ORPHA:95720 | 6 | 10 | 1.67e-13 | 2.66e-10 | 32 | LOW | novel | - |
| Glu | ORPHA:524 | 7 | 69 | 2.19e-09 | 2.14e-06 | 0 | VERY LOW | novel | - |
| Glu | ORPHA:394 | 7 | 71 | 2.68e-09 | 2.14e-06 | 85 | VERY HIGH | known | 29101223 |
| Glu | ORPHA:65285 | 5 | 34 | 6.94e-08 | 4.43e-05 | 19 | VERY LOW | novel | - |
| Glu | ORPHA:79233 | 4 | 15 | 1.18e-07 | 6.28e-05 | 100 | VERY HIGH | novel | - |
| Glu | ORPHA:510 | 3 | 16 | 1.59e-05 | 7.25e-03 | 100 | VERY HIGH | novel | - |
| Glu | ORPHA:199 | 4 | 54 | 2.49e-05 | 9.94e-03 | 0 | VERY LOW | novel | - |
| Glu | OMIM:615771 | 2 | 4 | 5.72e-05 | 2.02e-02 | 34 | LOW | known | 30658464 |
| Glu | ORPHA:3071 | 3 | 29 | 1.01e-04 | 3.22e-02 | 68 | HIGH | novel | - |
| Glu | ORPHA:218 | 2 | 6 | 1.42e-04 | 3.24e-02 | 22 | LOW | novel | - |
| Glu | ORPHA:2771 | 2 | 6 | 1.42e-04 | 3.24e-02 | 33 | LOW | novel | - |
| Glu | ORPHA:324442 | 2 | 6 | 1.42e-04 | 3.24e-02 | 48 | MEDIUM | novel | - |
| Glu | ORPHA:84142 | 2 | 6 | 1.42e-04 | 3.24e-02 | 48 | MEDIUM | novel | - |
| Hy | ORPHA:98879 | 2 | 63 | 6.01e-06 | 1.92e-02 | 0 | VERY LOW | novel | - |
| Ma | ORPHA:35878 | 6 | 10 | 2.05e-23 | 6.54e-20 | 13 | VERY LOW | ERD | 25418362 |
| Me | ORPHA:803 | 21 | 149 | 4.49e-15 | 1.43e-11 | 100 | VERY HIGH | known | 29243911<br>26895297 |
| Me | ORPHA:3208 | 19 | 130 | 4.59e-14 | 7.33e-11 | 99 | VERY HIGH | novel | - |
| Me | ORPHA:2309 | 11 | 47 | 5.04e-11 | 5.36e-08 | 40 | LOW | novel | - |
| Me | ORPHA:2995 | 9 | 30 | 2.55e-10 | 2.03e-07 | 49 | MEDIUM | novel | - |
| Me | ORPHA:455 | 9 | 33 | 6.62e-10 | 4.22e-07 | 25 | LOW | novel | - |
| Me | ORPHA:52430 | 6 | 11 | 3.60e-09 | 1.81e-06 | 13 | VERY LOW | ERD | 29243911<br>26895297 |
| Me | ORPHA:524 | 11 | 69 | 3.97e-09 | 1.81e-06 | 100 | VERY HIGH | RE | 24395704 |
| Me | ORPHA:199340 | 9 | 41 | 5.43e-09 | 2.17e-06 | 100 | VERY HIGH | ERD | 29163212 |
| Me | ORPHA:892 | 14 | 158 | 7.38e-08 | 2.59e-05 | 48 | MEDIUM | RE | 30943211<br>27577262 |
| Me | ORPHA:79396 | 7 | 27 | 8.11e-08 | 2.59e-05 | 49 | MEDIUM | novel | - |
| Me | ORPHA:154 | 16 | 217 | 1.15e-07 | 3.34e-05 | 100 | VERY HIGH | known | 27600370 |
| Me | ORPHA:35878 | 5 | 10 | 1.39e-07 | 3.69e-05 | 16 | VERY LOW | RE | 23281078<br>17971302 |
| Me | ORPHA:217604 | 18 | 287 | 2.17e-07 | 5.32e-05 | 70 | HIGH | known | 27600370 |
| Me | ORPHA:29072 | 21 | 403 | 4.78e-07 | 1.09e-04 | 74 | HIGH | novel | - |
| Nedd | ORPHA:85458 | 2 | 6 | 9.24e-08 | 2.95e-04 | 6 | VERY LOW | ERD | 22805479 |
| Ni | ORPHA:139441 | 7 | 32 | 5.94e-13 | 1.90e-09 | 71 | HIGH | novel | - |
| Ni | ORPHA:524 | 7 | 69 | 1.78e-10 | 2.85e-07 | 0 | VERY LOW | novel | - |
| Ni | ORPHA:37553 | 4 | 17 | 4.90e-08 | 5.21e-05 | 2 | VERY LOW | RE | 11533140 |
| Ni | ORPHA:848 | 3 | 10 | 1.18e-06 | 9.44e-04 | 100 | VERY HIGH | known | 18841126<br>18658051<br>27066490 |
| Ni | ORPHA:255241 | 3 | 14 | 3.57e-06 | 2.28e-03 | 21 | LOW | novel | - |
| Ni | ORPHA:91387 | 6 | 236 | 1.36e-05 | 7.24e-03 | 70 | HIGH | RE | 31694393 |
| Ni | ORPHA:2573 | 2 | 4 | 2.80e-05 | 1.12e-02 | 70 | HIGH | novel | - |
| Ni | ORPHA:444092 | 2 | 4 | 2.80e-05 | 1.12e-02 | 5 | VERY LOW | RE | 7508323<br>22746191 |
| Ni | ORPHA:3071 | 3 | 29 | 3.50e-05 | 1.24e-02 | 82 | VERY HIGH | novel | - |
| Ni | ORPHA:68364 | 2 | 5 | 4.66e-05 | 1.49e-02 | 100 | VERY HIGH | known | 18841126<br>18658051 |

| | | | | | | | | | |
|---|---|---|---|---|---|---|---|---|---|
| | | | | | | | | | 27066490 |
| Ni | ORPHA:79276 | 3 | 35 | 6.21e-05 | 1.80e-02 | 100 | VERY HIGH | novel | - |
| Ni | ORPHA:2457 | 2 | 7 | 9.77e-05 | 2.60e-02 | 0 | VERY LOW | novel | - |
| Ni | ORPHA:330050 | 2 | 8 | 1.30e-04 | 3.19e-02 | 45 | MEDIUM | known | 25110733 |
| NG | ORPHA:79241 | 14 | 116 | 2.68e-14 | 8.54e-11 | 98 | VERY HIGH | novel | - |
| NG | ORPHA:324 | 11 | 133 | 1.01e-09 | 1.61e-06 | 97 | VERY HIGH | OVCS | 9620884 |
| NG | ORPHA:228349 | 5 | 11 | 4.37e-09 | 4.64e-06 | 22 | LOW | known | 20680390<br>10330339 |
| NG | ORPHA:82 | 7 | 40 | 5.84.e-09 | 4.66e-06 | 68 | HIGH | ERD | Hudson et al 2017 |
| NG | ORPHA:56970 | 5 | 20 | 1.40e-07 | 8.92e-05 | 38 | LOW | known | 30135544<br>30886856<br>10557270 |
| NG | ORPHA:79264 | 8 | 100 | 2.60e-07 | 1.38e-04 | 61 | HIGH | OVCS | 11462245<br>11136716<br>14736728 |
| NG | ORPHA:584 | 5 | 27 | 7.02e-07 | 3.20e-04 | 21 | LOW | RE | 23959878<br>30042467 |
| NG | ORPHA:26106 | 11 | 260 | 9.82e-07 | 3.53e-04 | 88 | VERY HIGH | ERD | 28052004 |
| NG | ORPHA:232 | 3 | 4 | 9.96e-07 | 3.53e-04 | 0 | VERY LOW | novel | - |
| NG | ORPHA:93616 | 4 | 14 | 1.49e-06 | 4.75e-04 | 0 | VERY LOW | RE | 30111543 |
| NG | ORPHA:803 | 8 | 149 | 5.33e-06 | 1.54e-03 | 0 | VERY LOW | RE | 22521585 |
| NG | ORPHA:903 | 5 | 42 | 6.85e-06 | 1.72e-03 | 36 | LOW | known | 20418283 |
| NG | ORPHA:1308 | 4 | 20 | 7.00e-06 | 1.72e-03 | 49 | MEDIUM | novel | - |
| NG | ORPHA:204 | 3 | 8 | 1.37e-05 | 2.91e-03 | 4 | VERY LOW | RE | 19461968 |
| NG | ORPHA:454700 | 3 | 8 | 1.37e-05 | 2.91e-03 | 4 | VERY LOW | RE | 19461968 |
| NG | ORPHA:60 | 5 | 55 | 2.62e-05 | 5.22e-03 | 53 | MEDIUM | known | 24892502 |
| NG | ORPHA:848 | 3 | 10 | 2.90e-05 | 5.45e-03 | 0 | VERY LOW | novel | - |
| NG | ORPHA:827 | 9 | 256 | 4.12e-05 | 7.30e-03 | 72 | HIGH | OVCS | 21721517 |
| NG | ORPHA:97369 | 5 | 62 | 4.70e-05 | 7.89e-03 | 63 | HIGH | known | 17042482<br>29146600<br>21695262<br>24163131 |
| NG | ORPHA:77259 | 6 | 112 | 8.27e-05 | 1.32e-02 | 92 | VERY HIGH | OVCS | 24020503<br>27023912 |
| NG | ORPHA:466 | 2 | 3 | 1.19e-04 | 1.81e-02 | 14 | VERY LOW | known | 30135544 |
| NG | ORPHA:768 | 15 | 841 | 3.43e-04 | 4.98e-02 | 13 | VERY LOW | known | 9925876 |
| OGa | ORPHA:898 | 14 | 60 | 3.31e-29 | 1.05e-25 | 100 | VERY HIGH | novel | - |
| OGa | ORPHA:89937 | 3 | 4 | 6.54e-09 | 1.04e-05 | 50 | MEDIUM | RE | 25051439 |
| OGa | ORPHA:559 | 3 | 11 | 2.68e-07 | 2.85e-04 | 41 | MEDIUM | novel | - |
| OGa | ORPHA:411602 | 3 | 38 | 1.34e-05 | 1.07e-02 | 10 | VERY LOW | novel | - |
| OGl | ORPHA:276152 | 8 | 36 | 6.61e-19 | 2.11e-15 | 73 | HIGH | novel | - |
| OGl | ORPHA:276161 | 3 | 14 | 1.14e-07 | 1.81e-04 | 61 | HIGH | novel | - |
| OGl | ORPHA:603 | 3 | 19 | 3.02e-07 | 3.21e-04 | 5 | VERY LOW | novel | - |
| OGl | ORPHA:178330 | 2 | 6 | 7.15e-06 | 5.70e-03 | 51 | MEDIUM | ERD | 30858582<br>30237983 |
| OGl | ORPHA:93616 | 2 | 14 | 4.32e-05 | 2.76e-02 | 31 | LOW | RE | 30858582<br>30237983 |
| OGl | ORPHA:740 | 2 | 18 | 7.26e-05 | 3.86e-02 | 0 | VERY LOW | novel | - |
| OG | ORPHA:903 | 7 | 42 | 2.79e-13 | 8.91e-10 | 74 | HIGH | known | 22616016 |
| OG | ORPHA:89937 | 3 | 4 | 1.22e-08 | 1.94e-05 | 59 | MEDIUM | RE | 25051439 |
| OG | ORPHA:98879 | 5 | 63 | 3.90e-08 | 4.15e-05 | 97 | VERY HIGH | known | 32378017 |
| OG | ORPHA:849 | 4 | 56 | 1.49e-06 | 1.14e-03 | 26 | LOW | known | 22988088 |

| | | | | | | | | | |
|---|---|---|---|---|---|---|---|---|---|
| PC | OMIM:616214 | 6 | 8 | 2.22e-17 | 7.09e-14 | 72 | HIGH | known | 8138054 |
| PC | ORPHA:91378 | 4 | 21 | 5.57e-09 | 8.89e-06 | 5 | VERY LOW | RE | 25538858 |
| PC | ORPHA:100051 | 3 | 5 | 9.90e-09 | 1.05e-05 | 5 | VERY LOW | RE | 25538858 |
| PC | ORPHA:98879 | 4 | 63 | 5.38e-07 | 4.29e-04 | 28 | LOW | RE | 22872812 |
| PC | ORPHA:99885 | 4 | 91 | 2.36e-06 | 1.51e-03 | 73 | HIGH | known | 26600265 |
| PC | ORPHA:89937 | 2 | 4 | 6.09e-06 | 3.24e-03 | 35 | LOW | known | 12130585 |
| PC | ORPHA:82 | 3 | 40 | 9.54e-06 | 4.35e-03 | 8 | VERY LOW | known | 16359509<br>27248165 |
| PC | ORPHA:743 | 3 | 46 | 1.46e-05 | 5.82e-03 | 14 | VERY LOW | known | 16359509<br>27248165 |
| PC | ORPHA:98878 | 4 | 164 | 2.43e-05 | 8.62e-03 | 8 | VERY LOW | RE | 18492805<br>16839343 |
| PC | ORPHA:325 | 2 | 17 | 1.37e-04 | 4.37e-02 | 9 | VERY LOW | known | 30333979 |
| Ph | ORPHA:99818 | 287 | 611 | 1.72e-78 | 5.47e-75 | 100 | VERY HIGH | known | 29713058<br>10759189 |
| Ph | ORPHA:803 | 81 | 149 | 1.33e-28 | 2.12e-25 | 100 | VERY HIGH | RE | 28148298 |
| Ph | ORPHA:171680 | 31 | 34 | 1.37e-22 | 1.39e-19 | 98 | VERY HIGH | RE | 29057214 |
| Ph | ORPHA:102011 | 32 | 36 | 1.74e-22 | 1.39e-19 | 95 | VERY HIGH | RE | 29057214 |
| Ph | OMIM:192600 | 176 | 568 | 8.64e-22 | 5.51e-19 | 100 | VERY HIGH | RE | 26124132<br>25202278<br>27683560 |
| Ph | ORPHA:892 | 73 | 158 | 1.91e-20 | 1.01e-17 | 100 | VERY HIGH | known | 26973240 |
| Ph | ORPHA:227535 | 244 | 961 | 5.47e-17 | 2.49e-14 | 100 | VERY HIGH | known | 10373534<br>19016568<br>32286328 |
| Ph | ORPHA:2309 | 32 | 47 | 3.91e-16 | 1.56e-13 | 90 | VERY HIGH | novel | - |
| Ph | ORPHA:2995 | 24 | 30 | 4.79e-15 | 1.70e-12 | 100 | VERY HIGH | novel | - |
| Ph | ORPHA:98733 | 55 | 127 | 2.74e-14 | 8.73e-12 | 100 | VERY HIGH | RE | 20052757<br>18378677 |
| Ph | ORPHA:217569 | 117 | 396 | 2.19e-13 | 6.35e-11 | 100 | VERY HIGH | RE | 26124132<br>25202278<br>27683560 |
| Ph | ORPHA:154 | 76 | 217 | 3.42e-13 | 9.08e-11 | 100 | VERY HIGH | RE | 26124132<br>25202278<br>27683560 |
| Ph | OMIM:211980 | 37 | 75 | 3.64e-12 | 8.93e-10 | 74 | HIGH | RE | 26918193 |
| Ph | ORPHA:2869 | 55 | 142 | 6.53e-12 | 1.49e-09 | 100 | VERY HIGH | RE | 29399144<br>9837816<br>9425897 |
| Ph | ORPHA:524 | 33 | 69 | 1.51e-10 | 3.22e-08 | 100 | VERY HIGH | OVCS | 24173284 |
| Ph | ORPHA:139441 | 20 | 32 | 1.39e-09 | 2.77e-07 | 96 | VERY HIGH | novel | - |
| Ph | ORPHA:217604 | 82 | 287 | 4.41e-09 | 8.28e-07 | 100 | VERY HIGH | RE | 26124132<br>25202278<br>27683560 |
| Ph | ORPHA:3095 | 14 | 18 | 5.53e-09 | 9.28e-07 | 3 | VERY LOW | known | 25165434<br>22302819<br>31717404 |
| Ph | ORPHA:740 | 14 | 18 | 5.53e-09 | 9.28e-07 | 94 | VERY HIGH | known | 32208162<br>24741066 |
| SUMO | ORPHA:227535 | 52 | 961 | 8.26e-19 | 2.64e-15 | 100 | VERY HIGH | known | 22697792 |
| SUMO | ORPHA:171680 | 10 | 34 | 7.79e-12 | 1.24e-08 | 94 | VERY HIGH | RE | 23034179 |
| SUMO | ORPHA:102011 | 10 | 36 | 1.48e-11 | 1.57e-08 | 94 | VERY HIGH | RE | 23034179 |
| SUMO | ORPHA:902 | 15 | 135 | 1.59e-10 | 1.27e-07 | 99 | VERY HIGH | RE | 10806190<br>12771022 |
| SUMO | ORPHA:447877 | 32 | 733 | 1.14e-09 | 7.24e-07 | 100 | VERY HIGH | novel | - |
| SUMO | ORPHA:2309 | 9 | 47 | 5.71e-09 | 3.04e-06 | 32 | LOW | novel | - |
| SUMO | ORPHA:84 | 34 | 882 | 7.83e-09 | 3.57e-06 | 100 | VERY HIGH | known | 27148358 |

| | | | | | | | | | |
|---|---|---|---|---|---|---|---|---|---|
| | | | | | | | | | 21896657 |
| SUMO | ORPHA:300751 | 7 | 24 | 1.23e-08 | 4.92e-06 | 76 | HIGH | known | 18606848 |
| SUMO | ORPHA:217604 | 18 | 287 | 2.59e-08 | 9.17e-06 | 88 | VERY HIGH | known | 18606848 |
| SUMO | ORPHA:284417 | 4 | 5 | 1.15e-07 | 3.66e-05 | 38 | LOW | novel | - |
| SUMO | ORPHA:524 | 9 | 69 | 1.87e-07 | 5.41e-05 | 100 | VERY HIGH | RE | 21183956 |
| SUMO | ORPHA:3103 | 5 | 17 | 1.56e-06 | 4.14e-04 | 43 | MEDIUM | RE | 25054091 |
| Sul | ORPHA:98878 | 2 | 164 | 1.37e-05 | 4.38e-02 | 68 | HIGH | known | 8051097 |
| Ub | ORPHA:447877 | 107 | 733 | 5.40e-23 | 1.72e-19 | 100 | VERY HIGH | known | 15355978 |
| Ub | ORPHA:139441 | 19 | 32 | 3.90e-17 | 6.23e-14 | 100 | VERY HIGH | novel | - |
| Ub | OMIM:211980 | 24 | 75 | 1.40e-13 | 1.49e-10 | 61 | HIGH | known | 25820571 |
| Ub | ORPHA:84 | 97 | 882 | 6.81e-13 | 5.43e-10 | 100 | VERY HIGH | known | 21896657 |
| Ub | ORPHA:1340 | 21 | 64 | 2.67e-12 | 1.70e-09 | 70 | HIGH | novel | - |
| Ub | ORPHA:144 | 42 | 245 | 3.74e-12 | 1.99e-09 | 93 | VERY HIGH | known | 31697235<br>23248292<br>8797773<br>11920458<br>9322509 |
| Ub | ORPHA:394 | 21 | 71 | 2.53e-11 | 1.15e-08 | 94 | VERY HIGH | novel | - |
| Ub | ORPHA:51 | 22 | 85 | 1.54e-10 | 6.15e-08 | 91 | VERY HIGH | RE | 23979357 |
| Ub | ORPHA:443909 | 36 | 217 | 3.24e-10 | 1.14e-07 | 89 | VERY HIGH | RE | 23248292 |
| Ub | ORPHA:171680 | 14 | 34 | 3.58e-10 | 1.14e-07 | 0 | VERY LOW | RE | 24702957 |
| Ub | ORPHA:324 | 27 | 133 | 5.36e-10 | 1.51e-07 | 100 | VERY HIGH | novel | - |
| Ub | ORPHA:2309 | 16 | 47 | 5.68e-10 | 1.51e-07 | 54 | MEDIUM | novel | - |
| Ub | ORPHA:102011 | 14 | 36 | 8.86e-10 | 2.18e-07 | 0 | VERY LOW | RE | 24702957 |
| Ub | ORPHA:740 | 10 | 18 | 3.17e-09 | 7.22e-07 | 95 | VERY HIGH | novel | - |
| Ub | ORPHA:910 | 24 | 119 | 5.50e-09 | 1.17e-06 | 71 | HIGH | known | 16473935<br>25118285 |
| Ub | ORPHA:99947 | 12 | 29 | 6.16e-09 | 1.23e-06 | 70 | HIGH | known | 28335037<br>23727017 |
| Ub | ORPHA:277 | 13 | 37 | 1.52e-08 | 2.84e-06 | 100 | VERY HIGH | novel | - |
| Ub | ORPHA:98306 | 13 | 39 | 3.14e-08 | 5.57e-06 | 82 | VERY HIGH | novel | - |
| Ub | ORPHA:171881 | 7 | 10 | 8.67e-08 | 1.38e-05 | 46 | MEDIUM | novel | - |
| Ub | ORPHA:35878 | 7 | 10 | 8.67e-08 | 1.38e-05 | 42 | MEDIUM | RE | 24130338<br>16777975 |
| Ub | ORPHA:300751 | 10 | 24 | 1.07e-07 | 1.63e-05 | 100 | VERY HIGH | RE | 12601813<br>31861129 |
| Ub | ORPHA:101000 | 7 | 11 | 2.28e-07 | 3.30e-05 | 54 | MEDIUM | known | 19580544 |
| Ub | ORPHA:3322 | 21 | 116 | 3.34e-07 | 4.63e-05 | 88 | VERY HIGH | novel | - |
| Ub | ORPHA:1359 | 6 | 9 | 1.21e-06 | 1.54e-04 | 98 | VERY HIGH | novel | - |
| Ub | ORPHA:284435 | 6 | 9 | 1.21e-06 | 1.54e-04 | 94 | VERY HIGH | novel | - |
| Ub | ORPHA:892 | 24 | 158 | 1.42e-06 | 1.74e-04 | 0 | VERY LOW | known | 10973499<br>22649785 |
| Ub | ORPHA:457260 | 7 | 14 | 2.07e-06 | 2.45e-04 | 71 | HIGH | RE | 26748699 |
| Ub | ORPHA:300570 | 6 | 10 | 2.89e-06 | 3.29e-04 | 86 | VERY HIGH | RE | 24702957 |
| Ub | ORPHA:100045 | 5 | 7 | 6.28e-06 | 6.88e-04 | 61 | HIGH | known | 28335037<br>23727017 |

### Table S10. Classification of diseases in families according to Orphanet classification.

| Disease | Family/ies |
|---|---|
| Fatal familial insomnia | neurologic disease |
| Ornithine transcarbamylase deficiency | neurologic disease, inborn errors of metabolism |
| Acquired Creutzfeldt-Jakob disease | neurologic disease |

| | |
|---|---|
| Epidermolysis bullosa simplex, Dowling-Meara type | skin disease |
| Muscular dystrophy | neurologic disease |
| Cortical dysplasia, complex, with other brain malformations 6 | neurologic disease, developmental defect during embryogenesis |
| Autosomal recessive spastic paraplegia type 20 | neurologic disease |
| Acute intermittent porphyria | neurologic disease, renal disease, inborn errors of metabolism, skin disease |
| Superficial epidermolytic ichthyosis | skin disease |
| Heinz body anemia | hematologic disease |
| Hereditary cerebral hemorrhage with amyloidosis | neurologic disease, systemic or rheumatologic disease |
| Cap myopathy | neurologic disease |
| Muscular dystrophy, Selcen type | neurologic disease |
| Severe combined immunodeficiency due to adenosine deaminase deficiency | immune disease, inborn errors of metabolism |
| Hereditary diffuse gastric cancer | gastroenterologic disease, neoplastic disease |
| Triose phosphate-isomerase deficiency | neurologic disease, hematologic disease, inborn errors of metabolism |
| Polyostotic fibrous dysplasia | bone disease, developmental defect during embryogenesis |
| Hemoglobinopathy Toms River | hematologic disease |
| Polymerase proofreading-related adenomatous polyposis | neoplastic disease, gastroenterologic disease |
| Cardiomyopathy, familial hypertrophic, 14 | cardiac disease |
| Atypical Rett syndrome | neurologic disease |
| Leukoencephalopathy with brain stem and spinal cord involvement-high lactate syndrome | neurologic disease, inborn errors of metabolism |
| Inclusion body myopathy with Paget disease of bone and frontotemporal dementia | bone disease, developmental defect during embryogenesis, neurologic disease |
| Generalized epidermolysis bullosa simplex, non-Dowling-Meara type | skin disease |
| Marinesco-Sjögren syndrome | eye disease, developmental defect during embryogenesis, neurologic disease |
| Hyperproinsulinemia | endocrine disease |
| Mucopolysaccharidosis type 7 | Sucking/swallowing disorder, developmental defect during embryogenesis, inborn errors of metabolism, eye disease, bone disease |
| Autosomal dominant intermediate Charcot-Marie-Tooth disease type C | neurologic disease |
| C syndrome | neurologic disease, developmental defect during embryogenesis, bone disease |
| Stargardt disease | eye disease |
| Hypomyelination with atrophy of basal ganglia and cerebellum | neurologic disease |
| Familial partial lipodystrophy | endocrine disease, skin disease |
| Sporadic Creutzfeldt-Jakob disease | neurologic disease |
| Hereditary thrombophilia due to congenital antithrombin deficiency | hematologic disease, systemic or rheumatologic disease, bone disease |
| Glanzmann thrombasthenia | hematologic disease |
| CLN2 disease | inborn errors of metabolism, neurologic disease |
| Lynch syndrome | neoplastic disease, gastroenterologic disease |
| Aicardi-Goutières syndrome | Sucking/swallowing disorder, neurologic disease, systemic or rheumatological disease of childhood, systemic or rheumatologic disease, immune disease |
| Turcot syndrome with polyposis | neurologic disease, gastroenterologic disease, neoplastic disease |
| Immunoerythromyeloid hypoplasia | immune disease |
| Biotinidase deficiency | neurologic disease, inborn errors of metabolism, skin disease |
| Costello syndrome | surgical cardiac disease, sucking/swallowing disorder, neurologic disease, skin disease, neoplastic disease, developmental defect during embryogenesis, cardiac disease |
| Glycogen storage disease due to lactate dehydrogenase H-subunit deficiency | inborn errors of metabolism |
| Renal tubular dysgenesis of genetic origin | renal disease, developmental defect during embryogenesis |
| Fanconi anemia | renal disease, skin disease, bone disease, developmental defect during embryogenesis, hematologic disease, neoplastic disease |
| Darier disease | skin disease |
| Cortical dysgenesis with pontocerebellar hypoplasia due to TUBB3 mutation | developmental defect during embryogenesis, neurologic disease |
| Lesch-Nyhan syndrome | hematologic disease, inborn errors of metabolism, neurologic disease, renal disease |

| | |
|---|---|
| Hyperinsulinism-hyperammonemia syndrome | inborn errors of metabolism, endocrine disease |
| Hereditary thrombophilia due to congenital protein S deficiency | bone disease, systemic or rheumatologic disease, hematologic disease |
| Isaac syndrome | neurologic disease |
| RARS-related autosomal recessive hypomyelinating leukodystrophy | neurologic disease |
| Permanent neonatal diabetes mellitus | endocrine disease |
| Hypertrophic cardiomyopathy | cardiac disease |
| Macrocephaly-intellectual disability-autism syndrome | developmental defect during embryogenesis, neurologic disease |
| Thyroid hypoplasia | endocrine disease |
| Hereditary angioedema | systemic or rheumatologic disease, allergic disease |
| Peutz-Jeghers syndrome | gastroenterologic disease, developmental defect during embryogenesis, eye disease, neoplastic disease, skin disease |
| Hereditary fructose intolerance | inborn errors of metabolism, renal disease, hepatic disease, gastroenterologic disease |
| Cardiomyopathy, familial hypertrophic, 1 | cardiac disease |
| X-linked intellectual disability-hypotonia-movement disorder syndrome | neurologic disease |
| Lethal encephalopathy due to mitochondrial and peroxisomal fission defect | neurologic disease, inborn errors of metabolism |
| Cushing syndrome due to macronodular adrenal hyperplasia | Infertility, endocrine disease |
| Moyamoya disease | neurologic disease |
| Glycogen storage disease due to phosphoglycerate kinase 1 deficiency | neurologic disease, inborn errors of metabolism, hematologic disease |
| Hypoxanthine guanine phosphoribosyltransferase partial deficiency | inborn errors of metabolism, hematologic disease, renal disease, neurologic disease |
| Hereditary angioedema type 2 | systemic or rheumatologic disease, allergic disease |
| Distal myopathy, Welander type | neurologic disease |
| Gaucher disease type 1 | cardiac disease, bone disease, neurologic disease, Sucking/swallowing disorder, inborn errors of metabolism, eye disease, respiratory disease, systemic or rheumatologic disease |
| Carney complex | neoplastic disease, eye disease, cardiac disease, skin disease, endocrine disease |
| Leigh syndrome with leukodystrophy | inborn errors of metabolism, eye disease, neurologic disease, Sucking/swallowing disorder |
| Hereditary breast cancer | gynecologic or obstetric disease, neoplastic disease |
| Disorder of ornithine metabolism | inborn errors of metabolism |
| Glycine encephalopathy | neurologic disease, inborn errors of metabolism |
| Beta-ketothiolase deficiency | inborn errors of metabolism |
| Hereditary pheochromocytoma-paraganglioma | neoplastic disease, renal disease, circulatory system disease, endocrine disease |
| Autosomal dominant hypophosphatemic rickets | endocrine disease, renal disease, bone disease, developmental defect during embryogenesis |
| Noonan syndrome and Noonan-related syndrome | developmental defect during embryogenesis, surgical cardiac disease |
| Dilated cardiomyopathy | cardiac disease |
| Congenital factor II deficiency | hematologic disease |
| Hemophilia B | hematologic disease |
| Cardiofaciocutaneous syndrome | developmental defect during embryogenesis, cardiac disease, skin disease, neurologic disease, surgical cardiac disease |
| Pearson syndrome | hematologic disease, inborn errors of metabolism, immune disease, gastroenterologic disease, Sucking/swallowing disorder, endocrine disease |
| Mandibuloacral dysplasia | bone disease, skin disease, developmental defect during embryogenesis, endocrine disease |
| Classic homocystinuria | neurologic disease, developmental defect during embryogenesis, inborn errors of metabolism, eye disease |
| Congenital factor X deficiency | hematologic disease |
| Baraitser-Winter cerebrofrontofacial syndrome | developmental defect during embryogenesis, eye disease, neurologic disease |
| Autoimmune interstitial lung disease-arthritis syndrome | respiratory disease, systemic or rheumatologic disease, renal disease, systemic or rheumatological disease of childhood |
| Cornelia de Lange syndrome | Sucking/swallowing disorder, neurologic disease, abdominal surgical disease, developmental defect during embryogenesis, surgical thoracic disease, eye disease, maxillo-facial surgical disease, otorhinolaryngologic disease, bone disease |

| | |
|---|---|
| Fabry disease | eye disease, inborn errors of metabolism, developmental defect during embryogenesis, Sucking/swallowing disorder, neurologic disease, skin disease, cardiac disease, renal disease |
| Rubinstein-Taybi syndrome due to CREBBP mutations | developmental defect during embryogenesis, eye disease, neoplastic disease, neurologic disease, endocrine disease, bone disease, renal disease |
| Succinyl-CoA:3-ketoacid CoA transferase deficiency | inborn errors of metabolism |
| Hereditary late-onset Parkinson disease | neurologic disease |
| Juvenile neuronal ceroid lipofuscinosis | neurologic disease, inborn errors of metabolism |
| Alexander disease | eye disease, neurologic disease |
| IMAGe syndrome | bone disease, urogenital disease, developmental defect during embryogenesis, endocrine disease |
| Multiple endocrine neoplasia type 4 | endocrine disease, neoplastic disease |
| Primary hyperoxaluria type 2 | renal disease, inborn errors of metabolism |
| Lung cancer | respiratory disease, neoplastic disease |
| Blackfan-Diamond anemia | hematologic disease, developmental defect during embryogenesis, inborn errors of metabolism, neoplastic disease, maxillo-facial surgical disease, otorhinolaryngologic disease |
| Mitochondrial trifunctional protein deficiency | neurologic disease, inborn errors of metabolism, cardiac disease |
| Pten hamartoma tumor syndrome with granular cell tumor, included | neoplastic disease |
| Hereditary nonpolyposis colon cancer | gastroenterologic disease, neoplastic disease |
| Xeroderma pigmentosum | skin disease, neoplastic disease, developmental defect during embryogenesis, neurologic disease |
| Syndrome with limb malformations as a major feature | developmental defect during embryogenesis |
| Von Hippel-Lindau disease | developmental defect during embryogenesis, neoplastic disease, eye disease, neurologic disease, endocrine disease, renal disease, circulatory system disease |
| Familial thoracic aortic aneurysm and aortic dissection | systemic or rheumatologic disease, surgical thoracic disease, circulatory system disease |
| Familial dilated cardiomyopathy with conduction defect due to LMNA mutation | cardiac disease |
| Hoyeraal-Hreidarsson syndrome | hematologic disease, developmental defect during embryogenesis, immune disease, neurologic disease |
| Hemophilia A | hematologic disease |
| Lhermitte-Duclos disease | neoplastic disease, developmental defect during embryogenesis, neurologic disease |
| Isolated succinate-CoQ reductase deficiency | Sucking/swallowing disorder, neurologic disease, inborn errors of metabolism |
| Autosomal dominant Charcot-Marie-Tooth disease type 2A2 | neurologic disease |
| Pachyonychia congenita | developmental defect during embryogenesis, skin disease |
| Autosomal recessive axonal neuropathy with neuromyotonia | neurologic disease |
| Familial long QT syndrome | cardiac disease |
| Werner syndrome | skin disease, developmental defect during embryogenesis, neoplastic disease, eye disease |
| Congenital muscular dystrophy due to LMNA mutation | neurologic disease |
| Bruck syndrome | odontologic disease, bone disease, developmental defect during embryogenesis |
| Alpha-1-antitrypsin deficiency | respiratory disease, inborn errors of metabolism, renal disease, hepatic disease |
| Cardiodysrhythmic potassium-sensitive periodic paralysis | cardiac disease, neurologic disease |
| Lissencephaly type 3 | neurologic disease, developmental defect during embryogenesis |
| Somatotropic adenoma | endocrine disease, neoplastic disease |
| Li-Fraumeni syndrome | neoplastic disease, endocrine disease, neurologic disease |
| KID syndrome | developmental defect during embryogenesis, eye disease, skin disease, otorhinolaryngologic disease |
| Sickle cell anemia | neurologic disease, endocrine disease, systemic or rheumatologic disease, hematologic disease, renal disease, infertility, gynecologic or obstetric disease, bone disease |
| Autosomal dominant limb-girdle muscular dystrophy type 1B | cardiac disease, neurologic disease |
| Von Willebrand disease | hematologic disease |
| Wagner disease | eye disease |
| Hutchinson-Gilford progeria syndrome | skin disease, bone disease, developmental defect during embryogenesis |
| Human prion disease | neurologic disease |
| mental retardation, autosomal dominant 42 | neurologic disease |

| | |
|---|---|
| Rubinstein-Taybi syndrome | bone disease, renal disease, neoplastic disease, eye disease, developmental defect during embryogenesis, neurologic disease, endocrine disease |
| Lissencephaly due to TUBA1A mutation | developmental defect during embryogenesis, neurologic disease |
| Interatrial communication | developmental defect during embryogenesis, surgical cardiac disease |
| Amyotrophic lateral sclerosis | neurologic disease |
| Desminopathy | cardiac disease, neurologic disease |
| Transaldolase deficiency | inborn errors of metabolism |
| Familial isolated dilated cardiomyopathy | cardiac disease |
| Autosomal recessive Emery-Dreifuss muscular dystrophy | cardiac disease, neurologic disease |
| Roberts syndrome | eye disease, developmental defect during embryogenesis, bone disease, otorhinolaryngologic disease, maxillo-facial surgical disease |
| 2-hydroxyglutaric aciduria | inborn errors of metabolism, neurologic disease |
| Multiple endocrine neoplasia | endocrine disease, neoplastic disease |
| Gyrate atrophy of choroid and retina | eye disease, inborn errors of metabolism, neurologic disease |
| Hemoglobinopathy | hematologic disease |
| Hemoglobin H disease | hematologic disease, endocrine disease |
| Phosphoserine aminotransferase deficiency | inborn errors of metabolism, neurologic disease |
| Beta-thalassemia | renal disease, endocrine disease, hematologic disease |
| Citrullinemia | inborn errors of metabolism |
| Very long chain acyl-CoA dehydrogenase deficiency | cardiac disease, inborn errors of metabolism, neurologic disease |

**Table S11. Crossmatching of PTM sites and phenotype-related nsSNVs**. Number of phenotype-related nsSNVs (form ClinVar and using HPO terms) and PTM sites matched grouped by PTM type. We include number of proteins and phenotypes affected. PTMs with no matches are not shown.

| PTM types | Phenotype-related nsSNVs disrupting PTMs | Affected PTM sites | Proteins | Phenotypes involved | Co-occurrences between Phenotype-related nsSNVs and affected PTMs |
|---|---|---|---|---|---|
| phosphorylation | 1,107 | 977 | 351 | 1,112 | 1,612 |
| ubiquitination | 309 | 209 | 122 | 457 | 354 |
| acetylation | 187 | 160 | 87 | 359 | 218 |
| methylation | 68 | 60 | 40 | 158 | 83 |
| N-linked glycosylation | 67 | 31 | 21 | 70 | 68 |
| SUMOylation | 50 | 39 | 24 | 105 | 57 |
| O-linked glycosylation | 28 | 26 | 15 | 150 | 35 |
| O-GalNAc glycosylation | 26 | 39 | 7 | 16 | 44 |
| nitrosylation | 17 | 14 | 10 | 26 | 17 |
| S-glutathionylation | 15 | 9 | 7 | 42 | 15 |
| O-GlcNAc glycosylation | 13 | 10 | 9 | 114 | 13 |
| proteolytic cleavage | 10 | 7 | 6 | 9 | 10 |
| carboxylation | 6 | 4 | 2 | 3 | 6 |
| neddylation | 3 | 6 | 2 | 6 | 6 |
| glycation | 1 | 1 | 1 | 3 | 1 |
| glutamine deamidation | 1 | 1 | 1 | 1 | 1 |
| protein-pyridoxal-5-phosphate linkage | 1 | 1 | 1 | 62 | 1 |

| ALL PTM TYPES | 1,542 | 1,594 | 1,723 | 1,364 | 2,541 |
|---|---|---|---|---|---|

**Table S12. Data to build a network of PTM types and phenotypes associations.** Together with the p-values and FDRs, Occupancy scores (OS) are also shown.

| PTM type | HPO term | Parent or child term | nsSNVs involved in association | nsSNVs associated to phenotype | p-value | FDR | OS | OS degree |
|---|---|---|---|---|---|---|---|---|
| Ac | HP:0001644 | Child | 21 | 188 | 8.45e-08 | 1.61e-04 | 100 | VERY HIGH |
| Ac | HP:0012026 | Parent | 8 | 37 | 8.62e-06 | 5.85e-03 | 48 | MEDIUM |
| Ac | HP:0100950 | Parent | 6 | 19 | 1.15e-05 | 5.85e-03 | 87 | VERY HIGH |
| Ac | HP:0008872 | Child | 3 | 3 | 2.50e-05 | 5.85e-03 | 21 | LOW |
| Ac | HP:0011968 | Parent | 3 | 3 | 2.50e-05 | 5.85e-03 | 21 | LOW |
| Ac | HP:0012381 | Child | 3 | 3 | 2.50e-05 | 5.85e-03 | 21 | LOW |
| Ac | HP:0040288 | Child | 3 | 3 | 2.50e-05 | 5.85e-03 | 21 | LOW |
| Ac | HP:0000297 | Child | 4 | 8 | 4.59e-05 | 5.85e-03 | 21 | LOW |
| Ac | HP:0001252 | Parent | 4 | 8 | 4.59e-05 | 5.85e-03 | 21 | LOW |
| Ac | HP:0001290 | Child | 4 | 8 | 4.59e-05 | 5.85e-03 | 21 | LOW |
| Ac | HP:0006829 | Child | 4 | 8 | 4.59e-05 | 5.85e-03 | 21 | LOW |
| Ac | HP:0009062 | Child | 4 | 8 | 4.59e-05 | e-03 | 21 | LOW |
| Ac | HP:0012389 | Child | 4 | 8 | 4.59e-05 | e-03 | 21 | LOW |
| Ac | HP:0030190 | Child | 4 | 8 | 4.59e-05 | e-03 | 21 | LOW |
| Ac | HP:0031139 | Child | 4 | 8 | 4.59e-05 | e-03 | 21 | LOW |
| Ac | HP:0000002 | Parent | 3 | 4 | 9.78e-05 | 9.83e-03 | 21 | LOW |
| Ac | HP:0001510 | Parent | 3 | 4 | 9.78e-05 | 9.83e-03 | 21 | LOW |
| Ac | HP:0008897 | Child | 3 | 4 | 9.78e-05 | 9.83e-03 | 21 | LOW |
| Ac | HP:0031087 | Child | 3 | 4 | 9.78e-05 | 9.83e-03 | 21 | LOW |
| Ac | HP:0001670 | Child | 25 | 399 | 1.94e-04 | 1.23e-02 | 16 | VERY LOW |
| Ac | HP:0005157 | Child | 25 | 399 | 1.94e-04 | 1.23e-02 | 16 | VERY LOW |
| Ac | HP:0031992 | Child | 25 | 399 | 1.94e-04 | 1.23e-02 | 16 | VERY LOW |
| Ac | HP:0000301 | Parent | 4 | 11 | 2.02E-04 | 1.23e-02 | 21 | LOW |
| Ac | HP:0008947 | Child | 4 | 11 | 2.02E-04 | 1.23e-02 | 21 | LOW |
| Ac | HP:0001639 | Child | 27 | 452 | 2.30E-04 | 1.23e-02 | 45 | MEDIUM |
| Ac | HP:0000486 | Parent | 3 | 5 | 2.30E-04 | 1.23e-02 | 21 | LOW |
| Ac | HP:0010877 | Child | 3 | 5 | 2.30E-04 | 1.23e-02 | 21 | LOW |
| Ac | HP:0020045 | Child | 3 | 5 | 2.30E-04 | 1.23e-02 | 21 | LOW |
| Ac | HP:0020049 | Child | 3 | 5 | 2.30E-04 | 1.23e-02 | 21 | LOW |
| Ac | HP:0025068 | Child | 3 | 5 | 2.30E-04 | 1.23e-02 | 21 | LOW |
| Ac | HP:0025069 | Child | 3 | 5 | 2.30E-04 | 1.23e-02 | 21 | LOW |
| Ac | HP:0025587 | Child | 3 | 5 | 2.30E-04 | 1.23e-02 | 21 | LOW |
| Ac | HP:0025588 | Child | 3 | 5 | 2.30E-04 | 1.23e-02 | 21 | LOW |
| Ac | HP:0025589 | Child | 3 | 5 | 2.30E-04 | 1.23e-02 | 21 | LOW |
| Ac | HP:0031775 | Child | 3 | 5 | 2.30E-04 | 1.23e-02 | 21 | LOW |
| Ac | HP:0032011 | Child | 3 | 5 | 2.30E-04 | 1.23e-02 | 21 | LOW |
| Ac | HP:0032012 | Child | 3 | 5 | 2.30E-04 | 1.23e-02 | 21 | LOW |
| Ac | HP:0008936 | Child | 4 | 12 | 2.96E-04 | 1.49e-02 | 21 | LOW |
| Ac | HP:0001263 | Parent | 6 | 33 | 3.36e-04 | 1.52e-02 | 12 | VERY LOW |

| | | | | | | | | |
|---|---|---|---|---|---|---|---|---|
| Ac | HP:0011342 | Child | 6 | 33 | 3.36e-04 | 1.52e-02 | 12 | VERY LOW |
| Ac | HP:0011343 | Child | 6 | 33 | 3.36e-04 | 1.52e-02 | 12 | VERY LOW |
| Ac | HP:0012736 | Child | 6 | 33 | 3.36e-04 | 1.52e-02 | 12 | VERY LOW |
| Ac | HP:0008288 | Parent | 8 | 62 | 4.06e-04 | 1.80e-02 | 10 | VERY LOW |
| Ac | HP:0001319 | Child | 4 | 13 | 4.18e-04 | 1.81e-02 | 21 | LOW |
| Ac | HP:0001511 | Child | 3 | 6 | 4.68e-04 | 1.86e-02 | 21 | LOW |
| Ac | HP:0004812 | Parent | 3 | 6 | 4.68e-04 | 1.86e-02 | 21 | LOW |
| Ac | HP:0004848 | Child | 3 | 6 | 4.68e-04 | 1.86e-02 | 21 | LOW |
| Ac | HP:0006721 | Parent | 3 | 6 | 4.68e-04 | 1.86e-02 | 21 | LOW |
| Ac | HP:0011344 | Child | 6 | 37 | 6.39e-04 | 2.49e-02 | 9 | VERY LOW |
| Ac | HP:0006727 | Child | 3 | 7 | 8.01e-04 | 2.99e-02 | 21 | LOW |
| Ac | HP:0001437 | Parent | 2 | 2 | 8.61e-04 | 2.99e-02 | 21 | LOW |
| Ac | HP:0001446 | Parent | 2 | 2 | 8.61e-04 | 2.99e-02 | 21 | LOW |
| Ac | HP:0002509 | Parent | 2 | 2 | 8.61e-04 | 2.99e-02 | 21 | LOW |
| Ac | HP:0006895 | Child | 2 | 2 | 8.61e-04 | 2.99e-02 | 21 | LOW |
| Ac | HP:0200049 | Child | 2 | 2 | 8.61e-04 | 2.99e-02 | 21 | LOW |
| Ca | HP:0008321 | Parent | 4 | 22 | 2.04e-09 | 1.96e-06 | 62 | HIGH |
| Ca | HP:0008354 | Child | 4 | 22 | 2.04e-09 | 1.96e-06 | 62 | HIGH |
| Glu | HP:0005990 | Child | 6 | 7 | 4.78e-17 | 1.83e-14 | 69 | HIGH |
| Glu | HP:0008188 | Parent | 6 | 7 | 4.78e-17 | 1.83e-14 | 69 | HIGH |
| Glu | HP:0008191 | Child | 6 | 7 | 4.78e-17 | 1.83e-14 | 69 | HIGH |
| Glu | HP:0011780 | Child | 6 | 7 | 4.78e-17 | 1.83e-14 | 69 | HIGH |
| Glu | HP:0100028 | Child | 6 | 7 | 4.78e-17 | 1.83e-14 | 69 | HIGH |
| Me | HP:0006740 | Parent | 6 | 54 | 9.16e-06 | 1.74e-02 | 16 | VERY LOW |
| Ni | HP:0002047 | Parent | 4 | 85 | 7.50e-06 | 1.43e-02 | 0 | VERY LOW |
| NG | HP:0001071 | Parent | 11 | 123 | 6.08e-10 | 1.16e-02 | 81 | VERY HIGH |
| NG | HP:0001976 | Parent | 5 | 32 | 2.99e-06 | 2.86e-03 | 44 | MEDIUM |
| OGa | HP:0007773 | Parent | 14 | 60 | 1.11e-25 | 4.25e-23 | 100 | VERY HIGH |
| OGa | HP:0007964 | Child | 14 | 60 | 1.11e-25 | 4.25e-23 | 100 | VERY HIGH |
| OGa | HP:0030490 | Child | 14 | 60 | 1.11e-25 | 4.25e-23 | 100 | VERY HIGH |
| OGa | HP:0030673 | Child | 14 | 60 | 1.11e-25 | 4.25e-23 | 100 | VERY HIGH |
| OGa | HP:0200071 | Child | 14 | 60 | 1.11e-25 | 4.25e-23 | 100 | VERY HIGH |
| OGa | HP:0011533 | Child | 14 | 61 | 1.45e-25 | 4.60e-23 | 100 | VERY HIGH |
| OGa | HP:0007769 | Parent | 14 | 62 | 1.87e-25 | 5.09e-23 | 100 | VERY HIGH |
| PC | HP:0002511 | Parent | 3 | 28 | 2.43e-05 | 3.50e-02 | 21 | LOW |
| PC | HP:0001976 | Parent | 3 | 32 | 3.67e-05 | 3.50e-02 | 5 | VERY LOW |
| Ph | HP:0001639 | Child | 137 | 452 | 2.10e-05 | 3.5e-02 | 100 | VERY HIGH |
| Ph | HP:0030078 | Child | 32 | 48 | 2.41e-15 | 2.30e-12 | 69 | HIGH |
| Ph | HP:0030358 | Parent | 30 | 45 | 1.85e-14 | 8,85e-12 | 67 | HIGH |
| Ph | HP:0030360 | Child | 30 | 45 | 1.85e-14 | 8,85e-12 | 67 | HIGH |
| Ph | HP:0030359 | Child | 30 | 46 | 4.54e-14 | 1.73e-11 | 67 | HIGH |
| Ph | HP:0001670 | Child | 118 | 399 | 2.81e-13 | 6.70e-11 | 100 | VERY HIGH |
| Ph | HP:0005157 | Child | 118 | 399 | 2.81e-13 | 6.70e-11 | 100 | VERY HIGH |
| Ph | HP:0031992 | Child | 118 | 399 | 2.81e-13 | 6.70e-11 | 100 | VERY HIGH |
| Ph | HP:0001644 | Child | 62 | 188 | 2.11e-09 | 4.49e-07 | 100 | VERY HIGH |
| Ph | HP:0006740 | Parent | 27 | 54 | 4.27e-09 | 8.15e-07 | 92 | VERY HIGH |
| Ph | HP:0030434 | Parent | 10 | 10 | 9.47e-09 | 1.64e-06 | 99 | VERY HIGH |
| Ph | HP:0007755 | Parent | 13 | 19 | 3.85e-07 | 6.13e-05 | 17 | VERY LOW |
| Ph | HP:0001723 | Child | 24 | 64 | 1.92e-05 | 2.82e-03 | 50 | MEDIUM |

| | | | | | | | | |
|---|---|---|---|---|---|---|---|---|
| Ph | HP:0001319 | Child | 9 | 13 | 2.35e-05 | 3.2e-03 | 9 | VERY LOW |
| Ph | HP:0002885 | Parent | 5 | 5 | 9.87e-05 | 1.26e-02 | 75 | HIGH |
| Ph | HP:0030080 | Child | 9 | 15 | 1.22e-04 | 1.45e-02 | 13 | VERY LOW |
| Ph | HP:0001638 | Parent | 21 | 59 | 1.53e-04 | 1.46e-02 | 40 | LOW |
| Ph | HP:0011663 | Child | 21 | 59 | 1.53e-04 | 1.46e-02 | 40 | LOW |
| Ph | HP:0011665 | Child | 21 | 59 | 1.53e-04 | 1.46e-02 | 40 | LOW |
| Ph | HP:0200127 | Child | 21 | 59 | 1.53e-04 | 1.46e-02 | 40 | LOW |
| Ph | HP:0005511 | Parent | 7 | 10 | 1.87e-04 | 1.66e-02 | 53 | MEDIUM |
| Ph | HP:0005152 | Child | 21 | 60 | 2.00e-04 | 1.66e-02 | 38 | LOW |
| Ph | HP:0012817 | Child | 21 | 60 | 2.00e-04 | 1.66e-02 | 38 | LOW |
| Ph | HP:0001716 | Parent | 8 | 13 | 2.34e-04 | 1.86e-02 | 95 | VERY HIGH |
| Ph | HP:0000297 | Child | 6 | 8 | 3.26e-4 | 2.01e-02 | 8 | VERY LOW |
| Ph | HP:0001252 | Parent | 6 | 8 | 3.26e-4 | 2.01e-02 | 8 | VERY LOW |
| Ph | HP:0001290 | Child | 6 | 8 | 3.26e-4 | 2.01e-02 | 8 | VERY LOW |
| Ph | HP:0006829 | Child | 6 | 8 | 3.26e-4 | 2.01e-02 | 8 | VERY LOW |
| Ph | HP:0012389 | Child | 6 | 8 | 3.26e-4 | 2.01e-02 | 8 | VERY LOW |
| Ph | HP:0030190 | Child | 6 | 8 | 3.26e-4 | 2.01e-02 | 8 | VERY LOW |
| Ph | HP:0031139 | Child | 6 | 8 | 3.26e-4 | 2.01e-02 | 8 | VERY LOW |
| Ph | HP:0008947 | Child | 7 | 11 | 4.48e-04 | 2.38e-02 | 5 | VERY LOW |
| Ph | HP:0012190 | Child | 7 | 11 | 4.48e-04 | 2.38e-02 | 12 | VERY LOW |
| Ph | HP:0012191 | Child | 7 | 11 | 4.48e-04 | 2.38e-02 | 12 | VERY LOW |
| Ph | HP:0012539 | Parent | 7 | 11 | 4.48e-04 | 2.38e-02 | 12 | VERY LOW |
| Ph | HP:0030069 | Child | 7 | 11 | 4.48e-04 | 2.38e-02 | 12 | VERY LOW |
| Ph | HP:0002884 | Parent | 5 | 6 | 5.15e-04 | 2.46e-02 | 84 | VERY HIGH |
| Ph | HP:0004812 | Parent | 5 | 6 | 5.15e-04 | 2.46e-02 | 2 | VERY LOW |
| Ph | HP:0004848 | Child | 5 | 6 | 5.15e-04 | 2.46e-02 | 2 | VERY LOW |
| Ph | HP:0006721 | Parent | 5 | 6 | 5.15e-04 | 2.46e-02 | 2 | VERY LOW |
| Ph | HP:0000097 | Parent | 4 | 4 | 6.26e-04 | 2.85e-02 | 20 | VERY LOW |
| Ph | HP:0007129 | Child | 4 | 4 | 6.26e-04 | 2.85e-02 | 38 | LOW |
| Ph | HP:0008936 | Child | 7 | 12 | 9.23e-04 | 4.12e-02 | 0 | VERY LOW |
| Ph | HP:0007559 | Parent | 10 | 22 | 9.83e-04 | 4.27e-02 | 57 | MEDIUM |
| SUMO | HP:0001644 | Child | 9 | 188 | 4.73e-06 | 9.03e-02 | 76 | HIGH |
| SUMO | HP:0012125 | Parent | 5 | 46 | 1.62e-05 | 1.55e-02 | 2 | VERY LOW |
| Ub | HP:0001071 | Parent | 25 | 123 | 1.65e-11 | 3.16e-08 | 100 | VERY HIGH |
| Ub | HP:0030078 | Child | 15 | 48 | 3.85e-10 | 3.67e-07 | 25 | LOW |
| Ub | HP:0030358 | Parent | 13 | 45 | 1.76e-08 | 8.40e-06 | 21 | LOW |
| Ub | HP:0030360 | Child | 13 | 45 | 1.76e-08 | 8.40e-06 | 21 | LOW |
| Ub | HP:0030359 | Child | 13 | 46 | 2.36e-08 | 9.02e-06 | 21 | LOW |
| Ub | HP:0012190 | Child | 6 | 11 | 1.90e-06 | 4.03e-04 | 17 | VERY LOW |
| Ub | HP:0012191 | Child | 6 | 11 | 1.90e-06 | 4.03e-04 | 17 | VERY LOW |
| Ub | HP:0012539 | Parent | 6 | 11 | 1.90e-06 | 4.03e-04 | 17 | VERY LOW |
| Ub | HP:0030069 | Child | 6 | 11 | 1.90e-06 | 4.03e-04 | 17 | VERY LOW |
| Ub | HP:0030080 | Child | 6 | 15 | 1.79e-05 | 3.42e-03 | 17 | VERY LOW |
| Ub | HP:0007437 | Parent | 4 | 7 | 9.34e-05 | 1.62e-02 | 11 | VERY LOW |
| Ub | HP:0001644 | Child | 19 | 188 | 2.63e-04 | 4.19e-02 | 98 | VERY HIGH |

**Table S13. Prediction of new potential pathogenic PTM sites functionally associated to specific diseases.** For every predicted association between PTM types and genetic diseases, we show the p-values and FDRs for the tests comparing pathogenicity predictions of *in silico* nsSNVs in PTMs of the same type but not affected by a disease-related nsSNV and pathogenicity predictions of *in silico* nsSNVs affecting PTMs of the same type not predicted to be involved in diseases. We add the number of PTM sites predicted to be involved in the disease and those that are not, based on relative pathogenicity scores (rps).

| PTM Type | Disease ID | p-value | FDR | PTM sites rps < 95 | PTM sites rps > 95 |
|---|---|---|---|---|---|
| Ac | ORPHA:289869 | 1.00e+01 | 1.00e+01 | 3 | 0 |
| Ac | ORPHA:2869 | NA | NA | NA | NA |
| Ac | ORPHA:3208 | NA | NA | NA | NA |
| Ac | ORPHA:217604 | 9.02e-08 | 1.65e-06 | 39 | 17 |
| Ac | ORPHA:29072 | 4.40e-01 | 8.45e-01 | 5 | 0 |
| Ac | ORPHA:35878 | 9.20e-01 | 1.00e+01 | 17 | 1 |
| Ac | OMIM:192600 | 1.96e-07 | 3.18e-06 | 38 | 14 |
| Ac | ORPHA:171680 | 9.45e-01 | 1.00e+01 | 4 | 0 |
| Ac | ORPHA:85173 | 4.74e-01 | 8.87e-01 | 1 | 0 |
| Ac | ORPHA:102011 | 9.45e-01 | 1.00e+01 | 4 | 0 |
| Ac | ORPHA:2309 | 9.93e-01 | 1.00e+01 | 2 | 0 |
| Ac | ORPHA:740 | 7.54e-01 | 1.00e+01 | 8 | 0 |
| Ac | ORPHA:746 | 1.00e+01 | 1.00e+01 | 27 | 0 |
| Ac | ORPHA:713 | NA | NA | NA | NA |
| Ac | ORPHA:664 | 1.00e+01 | 1.00e+01 | 3 | 0 |
| Ac | ORPHA:783 | 1.36e-28 | 6.65e-27 | 38 | 24 |
| Ac | ORPHA:134 | 5.39e-01 | 9.84e-01 | 7 | 1 |
| Ac | ORPHA:300751 | 3.96e-01 | 7.92e-01 | 7 | 0 |
| Ac | ORPHA:98855 | 3.96e-01 | 7.92e-01 | 7 | 0 |
| Ac | ORPHA:3322 | NA | NA | NA | NA |
| Ac | ORPHA:353277 | 1.36e-28 | 6.65e-27 | 38 | 24 |
| Ac | ORPHA:414 | 1.00e+01 | 1.00e+01 | 3 | 0 |
| Ac | ORPHA:284435 | 2.07e-01 | 4.88e-01 | 3 | 0 |
| Ac | ORPHA:1478 | 9.73e-02 | 2.58e-01 | 5 | 2 |
| Ac | OMIM:616973 | 7.59e-01 | 1.00e+01 | 1 | 0 |
| Ac | ORPHA:157973 | 7.54e-01 | 1.00e+01 | 8 | 0 |
| Ac | ORPHA:699 | 1.00e+01 | 1.00e+01 | 12 | 0 |
| Ac | ORPHA:98473 | 7.45e-03 | 3.30e-02 | 26 | 0 |
| Ac | ORPHA:868 | NA | NA | NA | NA |
| Ac | ORPHA:2995 | 1.00e+01 | 1.00e+01 | 2 | 0 |
| Ac | OMIM:601728 | 1.04e-02 | 4.23e-02 | 1 | 1 |
| Ac | ORPHA:93616 | 2.52e-02 | 5.75e-02 | 3 | 0 |

| | | | | | |
|---|---|---|---|---|---|
| Ac | ORPHA:109009 | 1.65e-37 | 2.42e-35 | 45 | 29 |
| Ac | ORPHA:407 | 9.28e-01 | 1.00e+01 | 2 | 0 |
| Ac | ORPHA:469 | NA | NA | NA | NA |
| Ac | ORPHA:79396 | 9.64e-01 | 1.00e+01 | 1 | 0 |
| Ac | ORPHA:232 | NA | NA | NA | NA |
| Ac | ORPHA:280615 | NA | NA | NA | NA |
| Ac | ORPHA:438114 | 2.35e-01 | 5.44e-01 | 1 | 2 |
| Ac | ORPHA:217569 | 9.89e-14 | 3.61e-12 | 34 | 17 |
| Ac | ORPHA:137898 | NA | NA | NA | NA |
| Ac | ORPHA:603 | 8.70e-10 | 2.12e-08 | 12 | 8 |
| Ac | ORPHA:101028 | 1.00e+01 | 1.00e+01 | 4 | 0 |
| Ac | ORPHA:832 | 1.00e+01 | 1.00e+01 | 6 | 0 |
| Ac | ORPHA:124 | NA | NA | NA | NA |
| Ac | OMIM:242880 | NA | NA | NA | NA |
| Ac | ORPHA:264 | 3.96e-01 | 7.92e-01 | 7 | 0 |
| Ac | ORPHA:68364 | 4.57e-01 | 8.67e-01 | 1 | 0 |
| Ac | ORPHA:26793 | 1.00e+01 | 1.00e+01 | 12 | 0 |
| Ac | ORPHA:93599 | NA | NA | NA | NA |
| Ac | ORPHA:91387 | 9.67e-01 | 1 | 1 | 0 |
| Ac | OMIM:613251 | 3.80e-06 | 3.47e-05 | 6 | 4 |
| Ac | ORPHA:65285 | 6.22e-01 | 1.00e+01 | 1 | 0 |
| Ac | ORPHA:58 | 1.28e-05 | 1.04e-04 | 1 | 0 |
| Ac | ORPHA:19 | 2.99e-02 | 9.49e-01 | 3 | 1 |
| Ac | ORPHA:477 | NA | NA | NA | NA |
| Ac | ORPHA:98306 | 5.99e-01 | 1.00e+01 | 12 | 2 |
| Ac | ORPHA:79399 | 9.64e-01 | 1.00e+01 | 1 | 0 |
| Ac | ORPHA:210548 | 1.04e-02 | 4.23e-02 | 1 | 1 |
| Ac | ORPHA:187 | 1.00e+01 | 1.00e+01 | 2 | 0 |
| Ac | ORPHA:98909 | 1.70e-03 | 9.18e-03 | 0 | 1 |
| ADP-r | ORPHA:93276 | NA | NA | NA | NA |
| ADP-r | ORPHA:189427 | NA | NA | NA | NA |
| ADP-r | ORPHA:96256 | NA | NA | NA | NA |
| Ca | ORPHA:328 | 3.62e-04 | 2.52e-03 | 1 | 1 |
| Ca | ORPHA:98879 | 6.37e-01 | 1.00e+01 | 2 | 3 |
| Glu | ORPHA:803 | 7.66e-01 | 1.00e+01 | 4 | 0 |
| Glu | ORPHA:95720 | NA | NA | NA | NA |
| Glu | ORPHA:524 | NA | NA | NA | NA |
| Glu | ORPHA:394 | NA | NA | NA | NA |
| Glu | ORPHA:65285 | NA | NA | NA | NA |
| Glu | ORPHA:79233 | 6.38e-01 | 1.00e+01 | 1 | 0 |
| Glu | ORPHA:510 | 6.38e-01 | 1.00e+01 | 1 | 0 |
| Glu | ORPHA:199 | 1.24e-03 | 7.56e-03 | 1 | 1 |
| Glu | OMIM:615771 | NA | NA | NA | NA |
| Glu | ORPHA:3071 | NA | NA | NA | NA |
| Glu | ORPHA:218 | 2.37e-02 | 7.87e-02 | 7 | 0 |

| | | | | | |
|---|---|---|---|---|---|
| Glu | ORPHA:2771 | NA | NA | NA | NA |
| Glu | ORPHA:324442 | NA | NA | NA | NA |
| Glu | ORPHA:84142 | NA | NA | NA | NA |
| Hy | ORPHA:98879 | NA | NA | NA | NA |
| Ma | ORPHA:35878 | 1.00e+01 | 1.00e+01 | 1 | 0 |
| Me | ORPHA:803 | 1.00e+01 | 1.00e+01 | 27 | 2 |
| Me | ORPHA:3208 | NA | NA | NA | NA |
| Me | ORPHA:2309 | 1.39e-03 | 7.78e-03 | 6 | 2 |
| Me | ORPHA:2995 | 8.08e-01 | 1.00e+01 | 1 | 0 |
| Me | ORPHA:455 | 9.84e-01 | 1.00e+01 | 1 | 0 |
| Me | ORPHA:52430 | 9.75e-03 | 4.19e-02 | 3 | 2 |
| Me | ORPHA:524 | NA | NA | NA | NA |
| Me | ORPHA:199340 | 4.11e-01 | 8.11e-01 | 4 | 0 |
| Me | ORPHA:892 | NA | NA | NA | NA |
| Me | ORPHA:79396 | 9.87e-01 | 1.00e+01 | 1 | 0 |
| Me | ORPHA:154 | 9.11e-01 | 1.00e+01 | 7 | 0 |
| Me | ORPHA:35878 | 1.37e-03 | 7.78e-03 | 1 | 1 |
| Me | ORPHA:217604 | 7.74e-01 | 1.00e+01 | 22 | 0 |
| Me | ORPHA:29072 | NA | NA | NA | NA |
| Nedd | ORPHA:85458 | 2.23e-02 | 7.58e-02 | 1 | 0 |
| NG | ORPHA:79241 | NA | NA | NA | NA |
| NG | ORPHA:324 | NA | NA | NA | NA |
| NG | ORPHA:228349 | 1.43e-02 | 5.23e-02 | 2 | 0 |
| NG | ORPHA:82 | 8.04e-01 | 1.00e+01 | 1 | 0 |
| NG | ORPHA:56970 | NA | NA | NA | NA |
| NG | ORPHA:79264 | 1.43e-02 | 5.23e-02 | 2 | 0 |
| NG | ORPHA:584 | NA | NA | NA | NA |
| NG | ORPHA:26106 | NA | NA | NA | NA |
| NG | ORPHA:232 | NA | NA | NA | NA |
| NG | ORPHA:93616 | 4.49e-02 | 1.36e-01 | 2 | 0 |
| NG | ORPHA:803 | 9.70e-01 | 1.00e+01 | 3 | 0 |
| NG | ORPHA:903 | 1.26e-01 | 3.17e-01 | 3 | 0 |
| NG | ORPHA:1308 | 8.44e-01 | 1.00e+01 | 1 | 0 |
| NG | ORPHA:204 | NA | NA | NA | NA |
| NG | ORPHA:454700 | NA | NA | NA | NA |
| NG | ORPHA:60 | NA | NA | NA | NA |
| NG | ORPHA:848 | NA | NA | NA | NA |
| NG | ORPHA:827 | 9.94e-01 | 1.00e+01 | 1 | 0 |
| NG | ORPHA:97369 | 9.94e-01 | 1.00e+01 | 11 | 0 |
| NG | ORPHA:77259 | 9.23e-01 | 1.00e+01 | 2 | 0 |
| NG | ORPHA:466 | NA | NA | NA | NA |
| NG | ORPHA:768 | NA | NA | NA | NA |
| Ni | ORPHA:139441 | 5.54e-02 | 1.65e-01 | 1 | 0 |
| Ni | ORPHA:524 | NA | NA | NA | NA |
| Ni | ORPHA:37553 | NA | NA | NA | NA |

| | | | | | |
|---|---|---|---|---|---|
| Ni | ORPHA:848 | NA | NA | NA | NA |
| Ni | ORPHA:255241 | 9.90e-01 | 1.00e+01 | 1 | 0 |
| Ni | ORPHA:91387 | NA | NA | NA | NA |
| Ni | ORPHA:2573 | NA | NA | NA | NA |
| Ni | ORPHA:444092 | NA | NA | NA | NA |
| Ni | ORPHA:3071 | 3.56e-01 | 7.63e-01 | 1 | 0 |
| Ni | ORPHA:68364 | NA | NA | NA | NA |
| Ni | ORPHA:79276 | NA | NA | NA | NA |
| Ni | ORPHA:2457 | NA | NA | NA | NA |
| Ni | ORPHA:330050 | 2.74e-05 | 2.10e-04 | 1 | 0 |
| OGa | ORPHA:898 | 5.35e-01 | 9.83e-01 | 59 | 2 |
| OGa | ORPHA:89937 | NA | NA | NA | NA |
| OGa | ORPHA:559 | 7.50e-01 | 1.00e+01 | 4 | 0 |
| OGa | ORPHA:411602 | NA | NA | NA | NA |
| OGl | ORPHA:276152 | 3.38e-06 | 3.29e-05 | 0 | 1 |
| OGl | ORPHA:276161 | 3.38e-06 | 3.29e-05 | 0 | 1 |
| OGl | ORPHA:603 | 7.81e-01 | 1 | 1 | 0 |
| OGl | ORPHA:178330 | 8.70e-01 | 1.00e+01 | 1 | 0 |
| OGl | ORPHA:93616 | 8.70e-01 | 1.00e+01 | 1 | 0 |
| OGl | ORPHA:740 | NA | NA | NA | NA |
| OG | ORPHA:903 | 2.84e-02 | 9.21e-02 | 9 | 0 |
| OG | ORPHA:89937 | NA | NA | NA | NA |
| OG | ORPHA:98879 | 1.00e+01 | 1.00e+01 | 4 | 0 |
| OG | ORPHA:849 | 1.17e-02 | 4.62e-02 | 2 | 0 |
| PC | OMIM:616214 | 2.52e-07 | 3.68e-06 | 6 | 1 |
| PC | ORPHA:91378 | NA | NA | NA | NA |
| PC | ORPHA:100051 | NA | NA | NA | NA |
| PC | ORPHA:98879 | NA | NA | NA | NA |
| PC | ORPHA:99885 | 1.97e-04 | 1.44e-03 | 1 | 0 |
| PC | ORPHA:89937 | NA | NA | NA | NA |
| PC | ORPHA:82 | NA | NA | NA | NA |
| PC | ORPHA:743 | 4.81e-04 | 3.19e-03 | 1 | 0 |
| PC | ORPHA:98878 | 2.03e-01 | 4.85e-01 | 1 | 0 |
| PC | ORPHA:325 | 1.00e+01 | 1.00e+01 | 3 | 0 |
| Ph | ORPHA:99818 | 1.33e-11 | 3.87e-10 | 6 | 1 |
| Ph | ORPHA:803 | 1.00e+01 | 1.00e+01 | 304 | 18 |
| Ph | ORPHA:171680 | 2.09e-02 | 7.25e-02 | 22 | 1 |
| Ph | ORPHA:102011 | 2.09e-02 | 7.25e-02 | 22 | 1 |
| Ph | OMIM:192600 | 9.01e-02 | 2.44e-01 | 454 | 32 |
| Ph | ORPHA:154 | 1.86e-01 | 4.61e-01 | 194 | 11 |
| Ph | OMIM:211980 | 3.82e-02 | 1.19e-01 | 217 | 9 |
| Ph | ORPHA:2869 | NA | NA | NA | NA |
| Ph | ORPHA:524 | 1.00e+01 | 1.00e+01 | 17 | 0 |
| Ph | ORPHA:139441 | 8.14e-06 | 6.99e-05 | 21 | 0 |
| Ph | ORPHA:217604 | 1.00e+01 | 1.00e+01 | 437 | 33 |

| | | | | | |
|---|---|---|---|---|---|
| Ph | ORPHA:3095 | NA | NA | NA | NA |
| Ph | ORPHA:740 | 1.00e+01 | 1.00e+01 | 27 | 0 |
| Ph | ORPHA:892 | NA | NA | NA | NA |
| Ph | ORPHA:227535 | 1.01e-06 | 1.17e-05 | 15 | 0 |
| Ph | ORPHA:2309 | 1.00e+01 | 1.00e+01 | 79 | 4 |
| Ph | ORPHA:2995 | 3.44e-07 | 4.56e-06 | 25 | 1 |
| Ph | ORPHA:98733 | 1.00e+01 | 1.00e+01 | 82 | 5 |
| Ph | ORPHA:217569 | 3.72e-01 | 7.76e-01 | 504 | 34 |
| Ph | ORPHA:154 | 1.86e-01 | 4.61e-01 | 194 | 11 |
| Sul | ORPHA:98878 | NA | NA | NA | NA |
| SUMO | ORPHA:227535 | 7.73e-02 | 2.21e-01 | 1 | 0 |
| SUMO | ORPHA:171680 | 3.09e-01 | 6.73e-01 | 3 | 0 |
| SUMO | ORPHA:102011 | 3.09e-01 | 6.73e-01 | 3 | 0 |
| SUMO | ORPHA:902 | NA | NA | NA | NA |
| SUMO | ORPHA:447877 | NA | NA | NA | NA |
| SUMO | ORPHA:2309 | NA | NA | NA | NA |
| SUMO | ORPHA:84 | 3.61e-01 | 7.64e-01 | 19 | 0 |
| SUMO | ORPHA:300751 | 7.05e-03 | 3.30e-02 | 3 | 0 |
| SUMO | ORPHA:217604 | 7.05e-03 | 3.30e-02 | 3 | 0 |
| SUMO | ORPHA:284417 | 8.66e-01 | 1.00e+01 | 5 | 0 |
| SUMO | ORPHA:524 | 1.00e+01 | 1.00e+01 | 1 | 0 |
| SUMO | ORPHA:3103 | 1.04e-06 | 1.17e-05 | 4 | 3 |
| Ub | ORPHA:98306 | 1.99e-01 | 4.84e-01 | 22 | 1 |
| Ub | ORPHA:171881 | 2.85e-01 | 6.40e-01 | 1 | 0 |
| Ub | ORPHA:35878 | 1.04e-01 | 2.72e-01 | 11 | 2 |
| Ub | ORPHA:300751 | 7.57e-02 | 2.21e-01 | 8 | 0 |
| Ub | ORPHA:101000 | NA | NA | NA | NA |
| Ub | ORPHA:3322 | 1.35e-02 | 5.18e-02 | 11 | 1 |
| Ub | ORPHA:1359 | 9.56e-01 | 1.00e+01 | 9 | 0 |
| Ub | ORPHA:284435 | 6.02e-01 | 1.00e+01 | 5 | 0 |
| Ub | ORPHA:892 | NA | NA | NA | NA |
| Ub | ORPHA:457260 | 1.25e-08 | 2.60e-07 | 6 | 0 |
| Ub | ORPHA:300570 | 8.63e-02 | 2.40e-01 | 2 | 0 |
| Ub | ORPHA:100045 | 1.00e+01 | 1.00e+01 | 19 | 0 |
| Ub | ORPHA:447877 | NA | NA | NA | NA |
| Ub | ORPHA:139441 | 4.29e-01 | 8.36e-01 | 2 | 0 |
| Ub | OMIM:211980 | 7.31e-03 | 3.30e-02 | 17 | 0 |
| Ub | ORPHA:84 | 2.25e-03 | 1.17e-02 | 38 | 3 |
| Ub | ORPHA:1340 | 1.24e-03 | 7.56e-03 | 16 | 0 |
| Ub | ORPHA:144 | 6.51e-01 | 1.00e+01 | 5 | 1 |
| Ub | ORPHA:394 | NA | NA | NA | NA |
| Ub | ORPHA:51 | 3.45e-03 | 1.74e-02 | 33 | 2 |
| Ub | ORPHA:443909 | 1.21e-01 | 3.10e-01 | 7 | 1 |
| Ub | ORPHA:171680 | 7.69e-01 | 1.00e+01 | 5 | 0 |
| Ub | ORPHA:324 | NA | NA | NA | NA |

| | | | | | |
|---|---|---|---|---|---|
| Ub | ORPHA:2309 | 9.93e-01 | 1.00e+01 | 9 | 1 |
| Ub | ORPHA:102011 | 7.69e-01 | 1.00e+01 | 5 | 0 |
| Ub | ORPHA:740 | 8.72e-02 | 2.40e-01 | 9 | 0 |
| Ub | ORPHA:910 | 5.65e-01 | 1.00e+01 | 36 | 1 |
| Ub | ORPHA:99947 | 9.96e-01 | 1.00e+01 | 8 | 0 |
| Ub | ORPHA:277 | 1.00e+01 | 1.00e+01 | 7 | 0 |
